# Regulatory roles of three-dimensional structures of chromatin domains

**DOI:** 10.1101/2022.07.22.501196

**Authors:** Kelly Yichen Li, Qin Cao, Huating Wang, Danny Leung, Kevin Y. Yip

**Affiliations:** Department of Computer Science and Engineering, The Chinese University of Hong Kong, Shatin, New Territories, Hong Kong SAR; Sanford Burnham Prebys Medical Discovery Institute, La Jolla, CA 92037, USA; School of Biomedical Sciences, The Chinese University of Hong Kong, Shatin, New Territories, Hong Kong SAR; Shenzhen Research Institute, The Chinese University of Hong Kong, Shenzhen, China; Department of Orthopaedics and Traumatology, Li Ka Shing Institute of Health Sciences, The Chinese University of Hong Kong, Shatin, New Territories, Hong Kong SAR; Center for Neuromusculoskeletal Restorative Medicine (CNRM), CUHK InnoHK Centres, The Chinese University of Hong Kong, Shatin, New Territories, Hong Kong SAR; Division of Life Science, The Hong Kong University of Science and Technology, Clear Water Bay, Kowloon, Hong Kong SAR

## Abstract

Transcriptional enhancers usually, but not always, regulate genes within the same topologically associating domain (TAD). We hypothesize that this incomplete insulation is due to three-dimensional structures of corresponding chromatin domains in individual cells: Whereas enhancers and genes buried inside the “core” of a domain interact mostly with other regions in the same domain, those on the “surface” can more easily interact with the outside. Here we show that a simple measure, the intra-TAD ratio, can quantify the “coreness” of a region with respect to single-cell domains it belongs. We show that domain surfaces are permissive for high gene expression, and cell type-specific active cis-regulatory elements (CREs), active histone marks, and transcription factor binding sites are enriched on domain surfaces, most strongly in chromatin subcompartments typically considered inactive. These findings suggest a “domain surface CRE” model of gene regulation. We also find that disease-associated non-coding variants are enriched on domain surfaces.

## 2 Introduction

Mammalian genomes have a complex three-dimensional (3D) architecture, such that meter-long DNA sequences can be compactly fitted into the micrometer-scale nucleus. The 3D genome architecture enables loci far apart on the same chromosome or even different chromosomes to come to physical proximity, which is important at various functional levels [1–6].

Understandings of chromatin structures have been greatly expanded by two complementary classes of methods, respectively based on imaging and chromosome conformation capture (3C) [7, 8]. Imaging methods enable direct visualization of spatial organization of whole chromosomes or individually labeled DNA regions within a single cell. While some existing imaging methods are highly advanced in visualizing large-scale chromosome organizations [9–11] and the simultaneous spatial distribution of RNA species [12–14], high-resolution imaging (with *≤*50kb consecutive probes) of whole mammalian genomes with the genomic locations of all imaged loci resolved is still not achieved. In contrast, high-throughput 3C-based methods, such as Hi-C [15, 16], can detect chromatin interactions in the whole genome at high resolution, but these methods provide only information about pairs of genomic loci in close proximity. Recent “multiplex” methods such as Chromatin-Interaction Analysis via Droplet-based and Barcode-linked Sequencing (ChIA-Drop) [17] and Split-Pool Recognition of Interactions by Tag Extension (SPRITE) [18] can identify chromatin interactions between multiple (*≥*2) loci simultaneously, but they still do not provide spatial coordinates of genomic regions directly as imaging methods do. It is therefore necessary to combine imaging and 3C-based data to gain a more holistic view of genome organization.

From Hi-C data, multiple types of structural components that constitute the 3D genome architecture have been observed, including compartments and subcompartments of megabases long on average, topologically associating domains (TADs) of tens to hundreds of kilobases long on average, and individual chromatin loops [19]. TADs are consecutive genomic regions with a clear enrichment of chromatin interactions among loci within the TAD as compared to the background distribution [20, 21], and they can be identified using various methods [22]. Previous studies have proposed that TADs are formed through loop extrusion, where loop-extruding factors form progressively larger loops until stalled by boundary proteins, such as CTCF (CCCTC-binding factor) [23, 24]. Regions near TAD boundaries usually have properties different from other regions within the TAD, such as enrichment of housekeeping genes and presence of a pair of convergent CTCF binding sites [20, 24]. Functionally, promoters tend to interact with transcriptional regulatory elements such as enhancers within the same TAD, but this insulation is incomplete as revealed by high-resolution promoter interaction profiling [25–27].

For Hi-C experiments performed on a population of cells in a bulk sample, the resulting data only reflect average interaction frequencies across the cells. In contrast, single-cell Hi-C (scHi-C) and single-nucleus Hi-C (snHi-C) [28–30] detect chromatin interactions in individual cells. Despite challenges in data interpretation due to dropout and low signal-to-noise ratio caused by data sparsity, TAD-like domains have been observed in individual cells. With the help of data imputation, variability of domain boundary appearance and location has been quantified [31]. In corroboration with these observations from sequencing data, super-resolution imaging has also shown that TAD-like chromatin domains at the mesoscale (hundreds of nanometers wide) appear in single cells [9, 32, 33], which roughly match the size of TADs. However, the boundaries of these chromatin domains vary substantially across single cells and TAD boundaries appear to be statistical constructs that emerge by aggregating across single cells [9, 29, 34–36].

A potential molecular mechanism of TAD-like domains in individual single cells (to be referred to as “scDomains” hereafter) emerged from the observation in imaging studies that these domains can take a somewhat globular structure in the 3D space [9, 33, 37, 38]. Quantified by either distance to inter-chromatin compartment or voxel intensity, scDomains can be divided into exterior perichromatin regions and interior core regions [32, 33]. As compared to core regions, perichromatin regions are enriched for transcriptional activities and active histone marks, and depleted of repressive histone marks [32, 33]. These observations suggest that the 3D structures of scDomains may have functional significance.

Here we hypothesize that due to the 3D structures of scDomains, genomic regions closer to the “core” are more isolated from loci outside the domain, while those closer to the “surface” are more exposed to outside loci and thus have a higher chance of interacting with them (Figure 1a). We further hypothesize that despite cell-to-cell variability, information about this core versus surface distinction is partially preserved in TADs detected from bulk Hi-C data, which contributes to their incomplete insulation. We also hypothesize that the “coreness” of a region with respect to the 3D structure of a scDomain can be approximated from 3C-based and multiplex data, and it negatively correlates with the region’s molecular activities by affecting its opportunities to interact with protein factors from outside.

**Figure 1:**
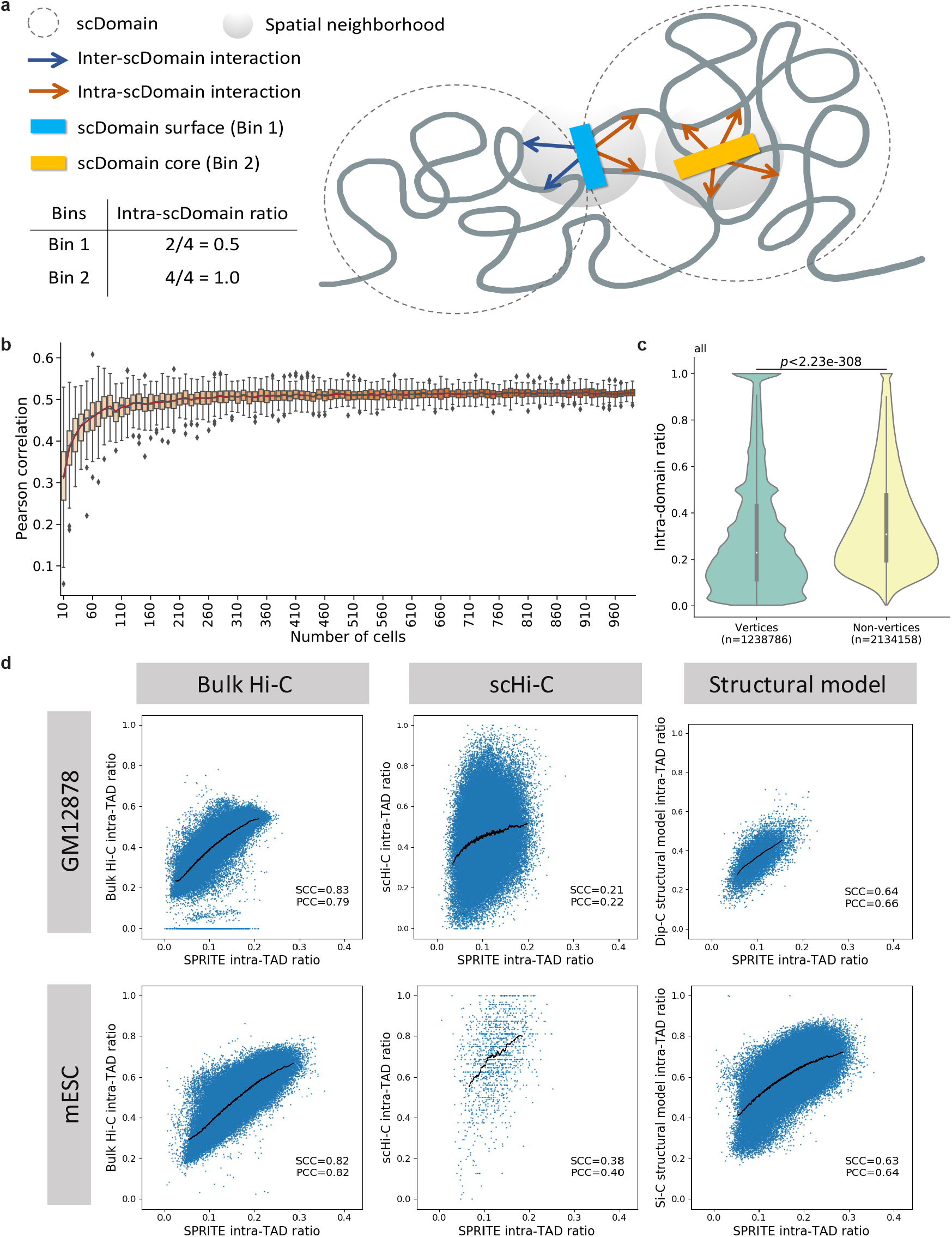
Calculation of intra-scDomain ratio and intra-TAD ratio using different types of data. (a) Schematic figure showing how the proportions of intra- and inter-scDomain interactions are related to the location of a genomic locus relative to the 3D structure of its scDomain. (b) Distribution of Pearson correlations between intra-TAD ratio and average intra-scDomain ratio in *x* single cells of IMR90 cells, for *x* equal to 10-990. Intra-TAD ratio was calculated using bulk Hi-C data [16]. Intra-scDomain ratio was calculated using single-cell imaging data [13]. For each value of *x*, we first sub-sampled *x* single cells and computed the average intra-scDomain ratio of each 50kb region over these cells. Then the vector of these average intra-scDomain ratios and the corresponding vector of intra-TAD ratios of the same 50kb regions were taken to compute a correlation. Finally, by repeating the sub-sampling of *x* cells 100 times, we obtained a distribution of correlation values. (c) Violin plots comparing the intra-scDomain ratios of regions on the convex hulls of scDomains (vertices) and other regions (non-vertices) defined using single-cell imaging data in IMR90 cells [13]. (d) Scatterplots comparing intra-TAD ratios calculated from SPRITE data [18] with those calculated from bulk Hi-C [16], scHi-C [34, 42], and structural models [34, 43]. Black curves represent moving averages. PCC: Pearson correlation coefficient. SCC: Spearman correlation coefficient.

## 3 Results

### 3.1 TAD-like domains in single cells exhibit 3D structural tendencies despite cell-to-cell variability

Considering the substantial cell-to-cell variability of scDomain boundaries, it is intriguing that TAD boundaries are usually fairly obvious in bulk Hi-C contact maps. To quantify the similarity between scDomains in different single cells and the corresponding bulk-aggregated TADs, we took a whole-chromosome tracing data set with individual 50kb regions on chromosome 21 of human IMR90 cells sequentially imaged at high resolution [13]. For each pair of 50kb genomic regions on chromosome 21 separated by a certain distance (from 100kb to 500kb), we labeled it as either “Same TAD” or “Different TADs” based on whether they belong to the same TAD called from IMR90 bulk Hi-C data. We then determined the number of cells in which these two regions also belong to the same scDomain, defined by the imaging data. The results (Figure S1) show that consistently across all distance thresholds, region pairs from the same TAD are significantly more often to belong to the same scDomain than distance-matched region pairs not from the same TAD. These results show that in terms of one-dimensional (1D) boundaries, scDomains display location tendencies that are consistent with the clear TAD boundaries. This is in line with previous findings that although chromatin interactions are dynamic and vary across different cells, 1D TAD boundaries tend to be more stable [31, 39, 40].

In a similar manner, we explored whether scDomains also display some tendencies in terms of their 3D structures. Using the same imaging data set, we defined a spatial neighborhood for each 50kb region (the “starting region”) based on its 3D coordinates and then computed the fraction of regions within this neighborhood coming from the same scDomain as the starting region (Methods). For a starting region buried inside the core of a scDomain 3D structure that is isolated from outside, this “intra-scDomain ratio” should be high (closer to 1); in contrast, for a starting region residing on the surface of a scDomain 3D structure that interacts with outside regions, its intra-scDomain ratio should be low (closer to 0) (Figure 1a). Analogously, for each 50kb region, we computed an “intra-TAD ratio” as the proportion of its Hi-C contacts coming from the same TAD. We found that the intra-scDomain ratio of a 50kb region averaged over a certain number of single cells correlates positively with its intra-TAD ratio, and the correlation reaches its maximum when averaging over 200 or more single cells (Figures 1b, S2a). These results show that despite cell-to-cell variability, the degree of insulation of a region provided by the 3D structure of its scDomain also displays some tendencies that are partially captured by bulk Hi-C data.

As a control, we shuffled the TADs and inter-TAD gaps while preserving the distributions of their 1D spans. Intra-TAD ratios computed from these shuffled TADs correlated much less strongly with average intra-scDomain ratios than real TADs (Figure S2b,c).

To further establish the relationship between intra-scDomain ratio and the “coreness” of a region relative to the 3D structure of its scDomain, we considered shape-related features. It has been shown by super-resolution imaging that TAD-like domains in single cells can take mostly globular shapes that are sometimes elongated [33], but a systematic analysis of the shapes of many scDomains has been missing. To avoid making any assumptions about scDomain shapes, we determined the convex hull of each scDomain based on the 3D coordinates of its constituent 50kb regions. Regardless of the actual shape of a scDomain, regions that define the convex hull (called the vertices) should be residing on the exterior of the 3D structure, while non-vertices can be on the exterior or in the interior. If intra-scDomain ratio represents the coreness of a region, the vertex regions should have lower intra-scDomain ratios than non-vertices on average. This was indeed the case based on the chromosome tracing data (Figure 1c). To make sure that this is not a bias caused by scDomain compactness or size, we also repeated the analysis with only regions in either a particular chromosome compartment or scDomains of a particular size range. In all cases, vertex regions had significantly lower intra-scDomain ratios than non-vertex regions (Figure S3).

Collectively, we have shown that a genomic region’s coreness with respect to the 3D structure of its scDomain exhibits some level of consistency across single cells and this tendency is partially captured by the intra-TAD ratio computed from bulk Hi-C data.

### 3.2 Intra-TAD ratios computed from different types of data show consistency

In order to study functional roles of 3D structures of scDomains, we need the capability to estimate the coreness of a genomic region relative to its scDomain in a variety of cell types with available functional data. Since intra-TAD ratios computed from bulk Hi-C data correlate with average intra-scDomain ratios in IMR90 cells (Figures 1b, S2a), we explored the possibility to compute similar intra-TAD ratios using other types of data as a practical proxy of intra-scDomain ratios.

Considering the ability to capture multiplex interactions from individual cells, we first explored SPRITE data. We computed intra-TAD ratios for all equal-sized genomic bins that tile the whole genome based on SPRITE data produced from the GM12878 human lymphoblastoid cells and mouse embryonic stem cells (mESCs) [18]. In general, the distribution of intra-TAD ratios remains highly stable for different sizes of the genomic bins (Figure S4). We developed and applied a data pre-processing procedure to ensure the robustness of the intra-TAD ratios and their analyses (Methods). Since TAD boundaries approximate average scDomain boundaries (Figure S1), and a 1D boundary of a scDomain is expected to be on the exterior of its 3D structure (for otherwise the segment between this “boundary” and the surface of the structure should also belong to this scDomain, thus invalidating the region as the boundary), we would expect intra-TAD ratios to be lower at TAD boundaries. Indeed, we observed that bins overlapping 1D TAD boundaries have significantly lower intra-TAD ratios than other bins within TADs, in both GM12878 and mESC (Figure S5a-d,i-l) (*p <* 2.23 *×* 10*^−^*^308^ in al cases, two-sided Mann-Whitney U test). To make sure that low intra-TAD ratios do not appear only at the 1D TAD boundaries, we examined the change of intra-TAD ratio across different bins within a TAD. We found that while the 1D TAD boundaries consistently had the lowest intra-TAD ratios, as expected, many other bins also had low intra-TAD ratios, including some “valleys” with particularly low intra-TAD ratios (Figure S5e-h,m-p), which likely correspond to genomic loci that tend to stay on domain surfaces. Some of these valleys overlap genes with high expression levels and regions with active chromatin signals, which suggest their potential functional significance (Figure S6).

Next, we compared intra-TAD ratios computed from SPRITE and bulk Hi-C. We computed intra-TAD ratios using bulk Hi-C data produced from GM12878 cells [16] and mESCs [41]. Since SPRITE can detect longer-distance chromatin interactions among regions in each barcoded complex [18], conceptually the intra-TAD ratios computed from SPRITE and bulk Hi-C data can be different (Figure S7). However, we still found their values highly consistent (Pearson correlation coefficient (PCC)=0.83 and Spearman correlation coefficient (SCC)=0.79 for GM12878; PCC=0.82 and SCC=0.82 for mESC) (Figure 1d), suggesting that 3D structures of scDomains are consistently captured by both multiplex SPRITE data and pairwise Hi-C data.

We also computed intra-TAD ratios using scHi-C data produced from GM12878 cells [34] and mESCs [42]. Despite the sparsity of scHi-C data, intra-TAD ratios computed from SPRITE and scHi-C data still correlated positively (Figure 1d).

We further used structural models derived from scHi-C data of GM12878 [34] and mESC [43] to determine the fraction of proximal loci of each genomic bin that are within the same TAD in the structural models, which can be considered an independent way of computing intra-TAD ratio. These structurally-derived intra-TAD ratios were found to be highly correlated with the intra-TAD ratios computed from SPRITE data (PCC=0.64 and SCC=0.66 for GM12878; PCC=0.63 and SCC=0.64 for mESC) (Figure 1d). When intra-TAD ratios computed from SPRITE data were used to color genomic loci in the structural models, the interior regions of TADs were indeed enriched for colors of large intra-TAD ratios (Figure S8).

In the above comparisons, for a region involved in multiple SPRITE clusters, we computed its average intra-TAD ratio across these clusters. We also repeated the same comparisons based on data from individual SPRITE clusters without averaging over different clusters (Figure S9). The resulting intra-TAD ratios computed from SPRITE data still correlate quite strongly with those computed from bulk Hi-C data, scHi-C data, and structural models.

In addition, given the variability of scDomains across single-cells, loci from two adjacent TADs could appear in the same scDomain frequently. To make sure that the intra-TAD ratio calculation is not dominated by such adjacent TADs, we checked the SPRITE data and found that contacts among loci from adjacent TADs constitute only a small proportion of all the inter-TAD contacts (Figure S10a,c). Besides, excluding contacts from adjacent TADs resulted in intra-TAD ratios highly correlated with the original intra-TAD ratios (Figure S10b,d), thus confirming that they are not dominated by contacts from adjacent TADs.

Taken together, these results show that the intra-TAD ratio, as a proxy of intra-scDomain ratio, can be computed using a variety of data. Considering the multiplex nature of SPRITE, in the following analyses we compute intra-TAD ratios using SPRITE data unless otherwise stated.

### 3.3 3D structural properties of chromatin domains differ across chromosome subcompartments

Previous studies have classified genomic regions roughly into the A (active) and B (inactive) compartments [15], and further divided them into different subcompartments [16, 44]. These subcompartments have been shown to be associated with various epigenomic signals and molecular activities, such as gene expression, histone modifications, and DNA replication timing [16, 44]. To see if the 3D structures of chromatin domains in different subcompartments have different properties, we compared the distributions of intra-TAD ratios of genomic bins in five subcompartments in GM12878 (A1, A2, B1, B2 and B3) [44]. We found that the intra-TAD ratio has different distributions in the different subcompartments, generally larger in the B2 and B3 subcompartments than in the A1, A2 and B1 subcompartments (Figure S11a). This could be related to previous observations that TAD-like domains in A1 and A2 are more dispersed [16, 37, 45], which allow core regions in these domains to be more accessible to regions outside the TAD and thus their lower intra-TAD ratios. We also found that TADs in B2 and B3 tend to be larger than TADs in the other subcompartments in terms of 1D genomic span (Figure S11b), which may also contribute to their larger intra-TAD ratios since core regions in a large TAD are expected to be more insulated from outside. This is supported by the fact that intra-TAD ratio is positively correlated with TAD size in all subcompartments (Figure S12). A recent study has demonstrated that DNA in smaller nucleosome clutches is more accessible and active, which is in line with our finding here since TADs may represent clusters of clutches [45]. The generally smaller TADs in the B1 subcompartment as compared to those in B2 and B3 likely explain their different distributions of intra-TAD ratios despite all three subcompartments belong to the inactive B compartment.

### 3.4 Genes on chromatin domain surfaces tend to have higher expression

Chromatin structures and gene regulation are coordinated during cell differentiation and other biological processes [2–4, 46], but how such an interplay is mediated by the 3D structures of chromatin domains has not been investigated in detail. To study whether 3D structures of chromatin domains are related to gene expression, we downloaded RNA-seq data produced from GM12878 cells and mESCs by ENCODE [47]. As expected, genes in A1 and A2 subcompartments have significantly higher expression than those in B1, B2, and B3 sub-compartments (Figure S13). Partitioning genes into equal-size groups based on intra-TAD ratios of their transcription start sites (TSSs), a clear trend is seen for genes with larger intra-TAD ratios to have lower expression levels (Figures 2a, S14a). The overall Spearman correlation between intra-TAD ratio and gene expression is −0.24 for GM12878 and −0.13 for mESCs. The same negative correlation is also observed when the SPRITE cluster size or the genomic bin size is changed to other values (Figures S15, S16). In contrast, the negative correlation largely disappears when the locations of TADs are randomly shuffled in GM12878 and mESC (Figures S17, S18). These findings suggest that genes with high expression levels tend to reside on domain surfaces. As to be shown below, this may make these genes more exposed to interactions coming from outside the domain such as transcription factor binding. To further characterize this dependency, we compared the intra-TAD ratios of the most highly expressed genes (top 1,000) and the most lowly expressed genes (bottom 1,000), and found a significant difference between them (Figures 2b, S14b). Since housekeeping genes tend to have high expression, we compared the intra-TAD ratios of TSSs of housekeeping genes [48] with those of other genes, and found that housekeeping genes have significantly smaller intra-TAD ratios (Figures 2c, S14c), even when TAD 1D boundaries and their neighboring regions were excluded (Figure S19).

**Figure 2:**
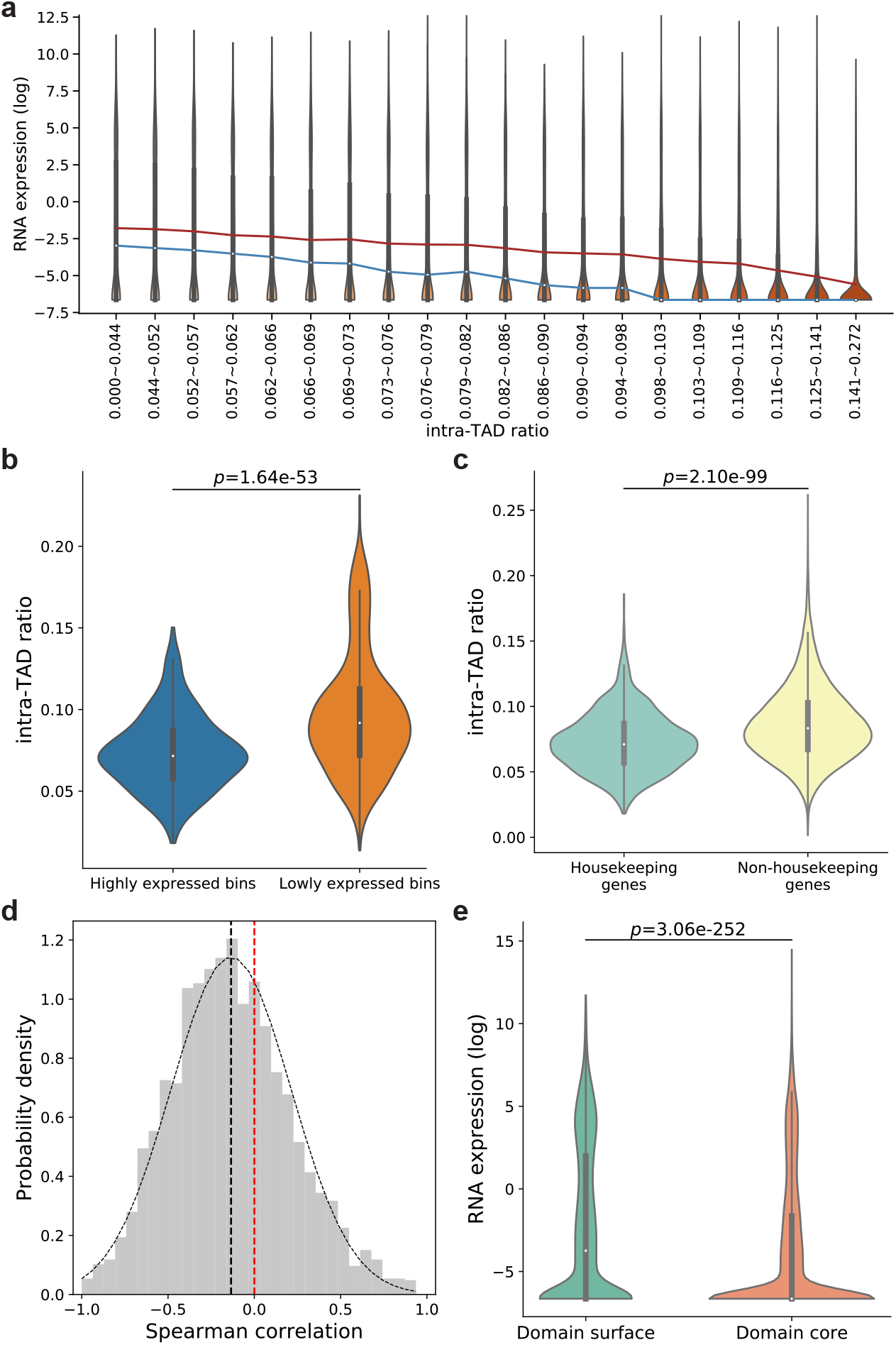
Inverse relationship between intra-TAD ratio and gene expression in GM12878 cells. (a) Violin plots of gene expression (log_2_(FPKM+0.01)) of TSS bins in groups with increasing intra-TAD ratio and similar number of bins. The blue and red lines connect median and mean values of the different groups, respectively. (b) Violin plots of intra-TAD ratios of the highly expressed bins and lowly expressed bins, defined as the genomic bins containing TSSs of genes whose FPKM values rank within top and bottom 1,000, respectively. (c) Violin plots of intra-TAD ratio of housekeeping genes and non-housekeeping genes. (d) Distribution of Spearman correlations between gene expression and intra-TAD ratio of genomic bins in individual TADs. Black dotted curve shows the fitted normal distribution. The vertical black dotted line shows the mean of Spearman correlations. The vertical red dotted line shows zero Spearman correlation. (e) Violin plots of gene expression levels (log_2_(FPKM+0.01)) in domain surface TSS bins and domain core TSS bins. *p*-values in all panels are calculated using the two-sided Mann-Whitney U test.

To rule out the possibility that the inverse relationship between intra-TAD ratio and gene expression is simply caused by TAD size or differences between different chromosome subcompartments, we computed the correlation between intra-TAD ratio and gene expression level across different genes in the same TAD for each TAD containing at least 5 genes. In this case, any effects due to TAD size and chromosome subcompartment would be eliminated. The resulting distribution of correlation values collected from these TADs has a mean significantly smaller than zero (Figures 2d, S14d, *p* = 4.61 *×* 10*^−^*^89^ for GM12878 and *p* = 3.59 *×* 10*^−^*^200^ for mESC, two-sided t-test). As a further confirmation, when we plotted these correlation values against TAD sizes for each chromosome subcompartment separately, we did not observe any significant correlation between them (Figure S20). These results show that the inverse relationship between gene expression level and intra-TAD ratio is not due to TAD size or subcompartment alone.

We also used an intra-TAD ratio threshold to classify genes into “domain core genes” and “domain surface genes” (Methods), and found that domain surface genes have significantly higher expression levels than domain core genes (*p* = 3.06 *×* 10*^−^*^252^ for GM12878 and *p <* 2.23 *×* 10*^−^*^308^ for mESC, two-sided Mann-Whitney U test) (Figures 2e, S14e).

We repeated the above analyses in different subcompartments separately and reached the same conclusions highly consistently across subcompartments (Figure S21). To ensure that the above results are not dominated by lowly expressed genes, we repeated the analyses but excluded TSS bins with extremely low expression (FPKM*<*0.02). The overall trends are highly consistent with those considering all genes (Figures S22, S23).

### 3.5 Chromatin signals are correlated with chromatin domain 3D structures

Chromatin signals such as chromatin accessibility and histone modifications have been shown to characterize chromosome subcompartments [16, 44], but whether they are associated with chromatin domain 3D structures remains unclear. We computed the correlation between intra-TAD ratio and each of 12 chromatin signals in GM12878, including chromatin accessibility signals from ATAC-seq and enrichment of 11 histone marks from ChIP-seq [47, 49] (Figure 3a). When considering all genomic bins together, ATAC-seq signals (Figure 3a,b) and most active histone marks (Figure 3a,c) are negatively correlated with intra-TAD ratio, while some repressive histone marks are positively correlated with intra-TAD ratio (Figure 3a,d). When considering genomic bins of different chromosome sub-compartments separately, stronger correlations are seen in the B2 and B3 subcompartments (Figure 3a). These observations are consistent with the negative correlation between gene expression and intra-TAD ratio shown above, that active chromatin signals are enriched on domain surfaces. Similar trends are also observed from the correlations between epigenomic signals and intra-TAD ratio in mESCs in the A and B compartments (Figure S24). Repeating the calculations with shuffled TADs, the correlations are much weaker (Figure S25).

**Figure 3:**
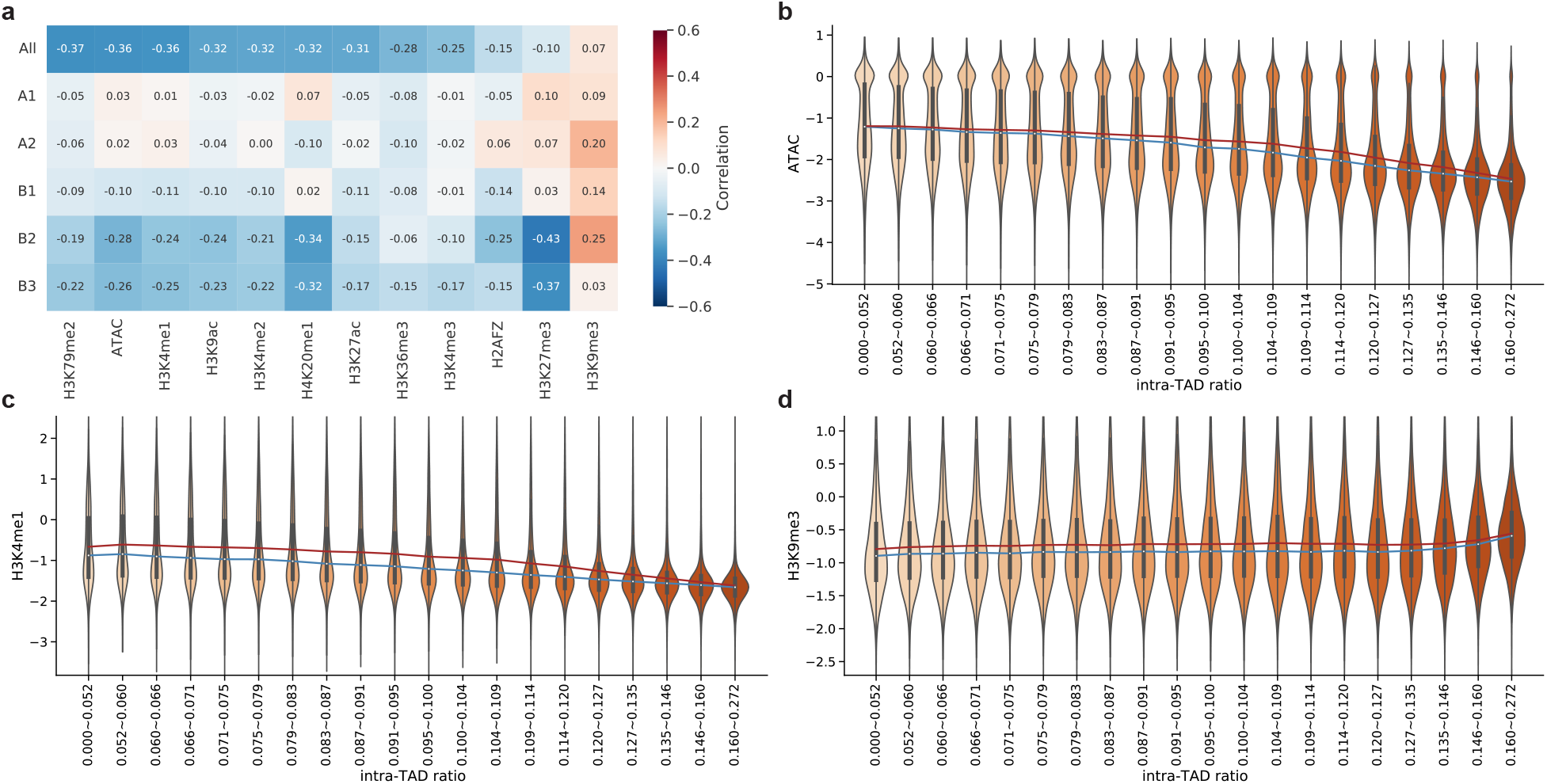
Correlation between intra-TAD ratio and chromatin signals. (a) Spearman correlation between intra-TAD ratio and ATAC-seq signals and 11 histone modifications across either all genomic bins or only bins in a genomic subcompartment. The columns are sorted by the correlation values when all TADs are considered (in the “All” row). (b)-(d) Violin plots of ATAC-seq signals (b), H3K4me1 (c), and H3K9me3 (d) in groups with increasing intra-TAD ratio and similar number of bins. y-axis shows log_2_(signal+0.01). The blue and red lines connect median and mean values of the different groups, respectively.

Interestingly, the correlation between the repressive mark H3K27me3 and intra-TAD ratio is slightly positive in A1 and A2 but quite negative in B2 and B3 (Figure 3a, S26). This is different from the universal positive correlation between the repressive mark H3K9me3 and intra-TAD ratio across all subcompartments. This may be related to differences between facultative and constitutive heterochromatin, that H3K27me3-marked regions can more easily be reactivated if they are on domain surfaces. Alternatively, this may also be due to the different molecular weights (MW) of the enzyme complexes that deposit H3K27me3 and H3K9me3. The H3K27me3 modifier polycomb repressive complex 2 (PRC2) (MW: *∼*300 kDa) has a larger molecular weight than H3K9me3 modifiers such as SETDB1 (MW: *∼*140 kDa), SETDB2 (MW: *∼*80 kDa), SUV39H1 (MW: *∼*50 kDa) and SUV39H2(MW: *∼*50 kDa) [33]. As a result, the large TADs in B2 and B3 subcompartments may impose stronger hindrance to the deposition of H3K27m3e to their core regions than to H3K9me3.

To further explore the effect of H3K9me3 and H3K27me3 on intra-TAD ratio, we built a linear regression model with H3K9me3 signal, H3K27me3 signal, and the interaction term between them as the explanatory variables, and intra-TAD ratio as the response variable. Consistent with the correlation analysis above, both H3K9me3 and H3K27me3 had positive model coefficients in the A1, A2, and B1 subcompartments, while the coefficients of H3K27me3 were much more negative than H3K9me3 in the B2 and B3 subcompartments (Figure S27). Surprisingly, the coefficients of the interaction term were consistently opposite in sign to the coefficients of the two independent variables, suggesting that the distribution of the two marks when they appear together with respect to chromatin domain 3D structures is different from the distribution when they appear separately. The exact reason for this discrepancy requires further investigations.

### 3.6 Cell type-specific ***cis***-regulatory elements and transcription factors are enriched on chromtain domain surfaces

As shown above, despite the different properties of individual chromatin domains, domain surfaces are generally more active than domain cores, especially for the domains in the B2 and B3 subcompartments. We hypothesized two alternative models for the domain surface genes to be regulated by *cis*-regulatory elements (CREs) in the same domain, namely having the CREs (such as enhancers) also residing on domain surfaces or being buried in the domain cores (Figure 4a). The former, “domain surface CRE” model, provides a better explanation for the CREs to be easily accessible by transcription factors, while the latter, “domain core CRE” model, provides a better explanation for the insulation of the CREs in a TAD from regulating genes outside the TAD. To test these models, we downloaded different categories of candidate CREs (cCREs) active in GM12878 from ENCODE [50] and examined their locations with respect to chromatin domain 3D structures based on intra-TAD ratios (Methods). Several categories of cCREs were found enriched on domain surfaces (Figure 4b) and depleted from domain cores (Figure S28a), most notably proximal enhancer-like signatures (pELS) and promoter-like signatures (PLS). This is not simply due to 1D TAD boundaries, as the enrichment of cCREs on domain surfaces and depletion of them from domain cores still hold when cCREs at and near TAD boundaries are excluded from the analysis (Figures 4c, S28b). To check whether the enrichment of cCREs on domain surfaces is related to cell type-specific activities or it simply reflects the locations of CREs regardless of their activities, we also downloaded cell type-agnostic cCREs and cCREs active in other cell types (Methods). When cCREs active in GM12878 were excluded from these sets, the remaining cCREs were no longer enriched on the surfaces of GM12878 domains (Figure 4d). Together, these results demonstrate that cCREs are enriched on domain surfaces in a cell type-specific manner.

**Figure 4:**
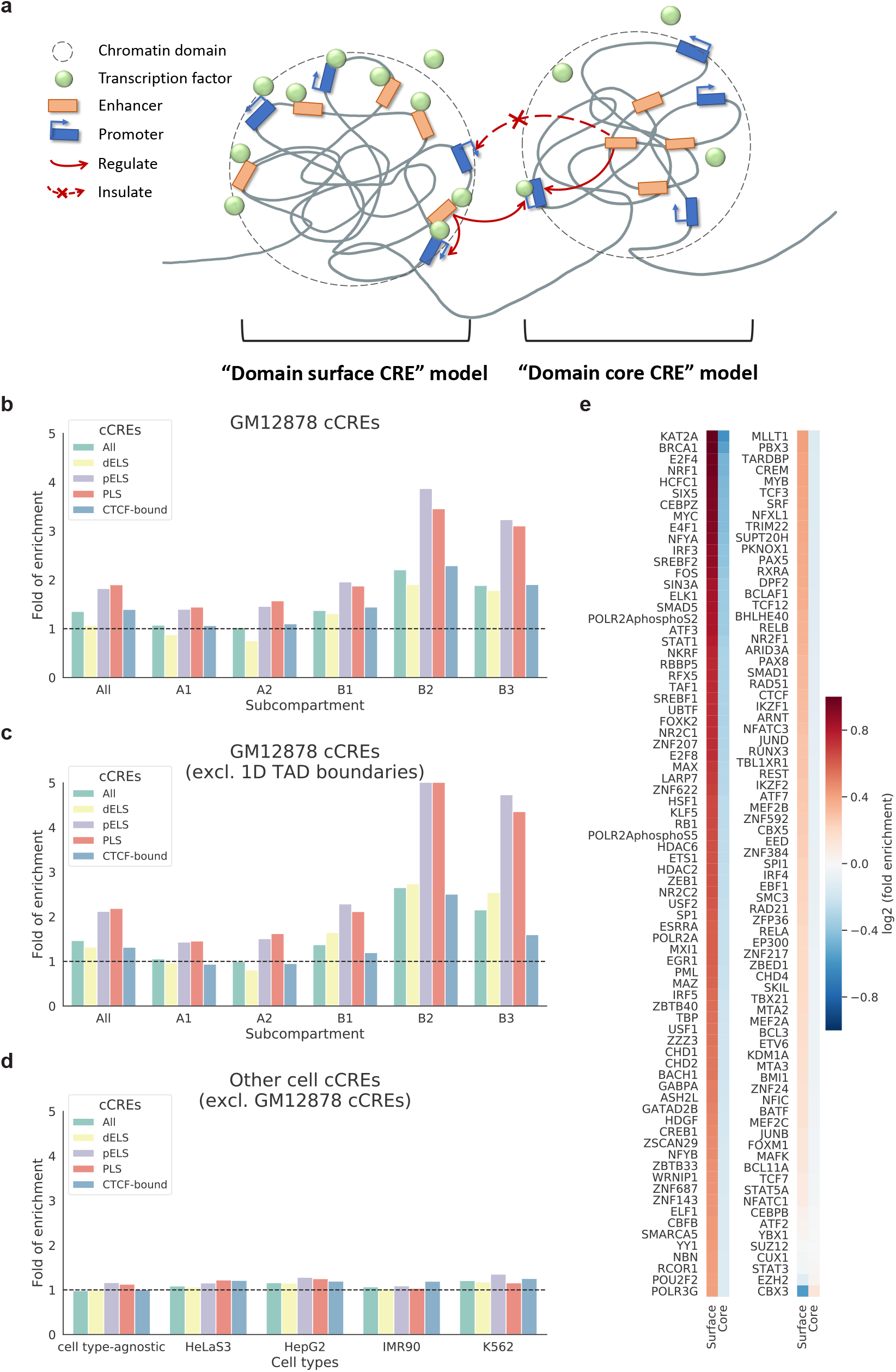
Enrichment of CREs and transcription factors on chromatin domain surfaces. (a) A schematic diagram showing the “domain surface CRE” model and the “domain core CRE” model. (b) Fold of enrichment of cCREs on domain surfaces in GM12878. (c) Fold of enrichment of cCREs on domain surfaces in GM12878 when regions around TAD boundaries are excluded. (d) Fold of enrichment of cCREs on domain surfaces when the cCREs are cell type-agnostic or defined in other cell types and the GM12878-specific cCREs are excluded. (e) Heatmap showing log2(enrichment fold) of transcription factor binding sites on domain surfaces and in domain cores in GM12878. cCREs All: all categories of cCREs; dELS: distal enhancer-like signatures; pELS: proximal enhancer-like signatures; PLS: promoter-like signatures; CTCF-bound: bound by CTCF.

To see whether residing on domain surfaces allows CREs to be easily bound by transcription factors, we next examined the enrichment of cell type-specific transcription factor binding sites on domain surfaces and in domain cores, based on ChIP-seq peaks of 153 TFs in GM12878 produced by ENCODE [47, 49]. Strikingly, the binding sites of almost all the TFs are enriched on domain surfaces and depleted from domain cores (Figures 4e, S29), consistent with the finding that enhancers and promoters are enriched on domain surfaces. The two most prominent exceptions are CBX3, which is associated with heterochromatin marked with H3K9me3, and EZH2, which is the enzymatic subunit of the polycomb repressive complex 2 that catalyzes H3K27me3. In contrast, the binding sites of the lysine acetyltransferase KAT2A, which is involved in the acytelation of H3K9 are strongly enriched on domain surfaces.

We also repeated the analysis using shuffled TADs and found that the enrichment of TFs on domain surfaces defined by real TADs is significantly stronger than that by shuffled TADs (Figure S30).

To check whether the enrichment of TFs on domain surfaces is cell type-specific, we also downloaded ChIP-seq peak locations of TFs in four other cell types. When the TF peaks in GM12878 were excluded from these sets, the remaining peaks had reduced enrichment on the GM12878 domain surfaces. Furthermore, when the cCREs in GM12878 were also excluded from these sets, the enrichment largely disappeared (Figures S31, S32, S33, S34). These results further support the “domain surface CRE” model, that the localization of CREs on domain surfaces allows TFs to bind them more easily in a cell type-specific manner.

### 3.7 Chromatin domain 3D structures correlate with replication timing and nuclear spatial compartmentalization

Previous studies have demonstrated that DNA replication timing has strong association with spatial genome organization. In general, loci in the active A compartment tend to replicate early, whereas loci in the repressive B compartment tend to replicate late [51]. Although TAD boundaries often align with boundaries of replication domains, the relationship between chromatin domain 3D structures and replication timing remains obscure [51].

To see whether replication timing is correlated with chromatin domain 3D structures, we analyzed Repli-seq data produced from GM12878 cells by ENCODE [47] and found that replication timing signal (higher value corresponding to earlier replication) negatively correlated with intra-TAD ratio when considering all the TADs genome-wide, and even more so for the TADs in the B subcompartments (Figures 5a, S35). These results suggest that 3D structures of chromatin domains play a stronger role in replication timing for genomic loci in the B subcompartments and that early replication tends to occur in regions on the surfaces of these domains.

**Figure 5:**
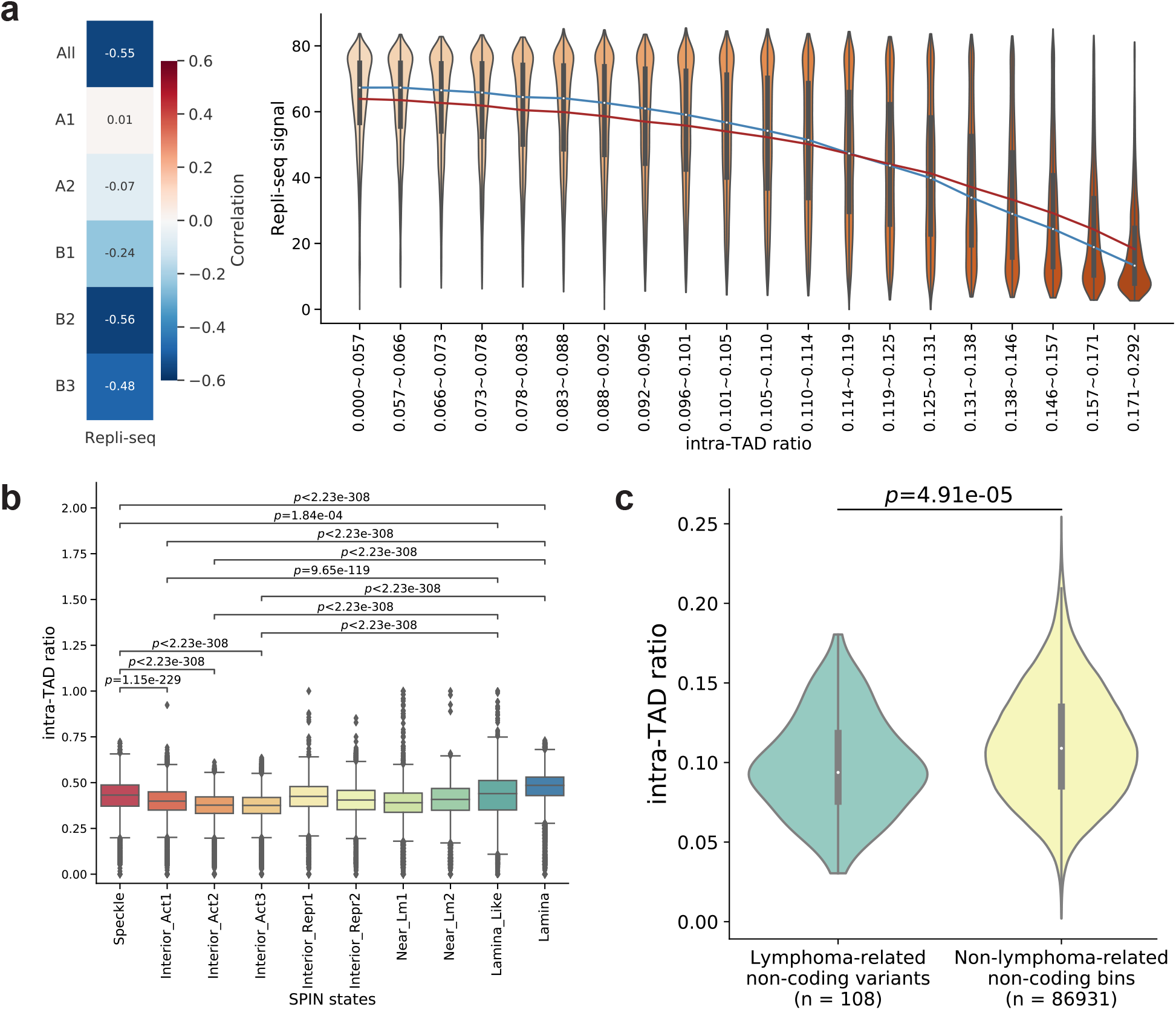
Intra-TAD ratio correlates with replication timing, nuclear compartments, and disease variants. (a) Replication timing is correlated with intra-TAD ratio. left panel: Spearman correlation between intra-TAD ratio and Repli-seq signals; right panel: Violin plots of Repliseq signals in groups with increasing intra-TAD ratio and similar number of bins. The blue and red lines connect median and mean values of the different groups, respectively. (b) Distribution of intra-TAD ratios of genomic bins in individual SPIN states. Speckle: genomic regions predicted to be associated with nuclear speckles; Interior Act1, Interior Act2, Interior Act3: predicted active interior regions; Interior Repr1, Interior Repr2: predicted repressive interior regions; Near Lm1, Near Lm2: regions predicted to be near nuclear lamina; Lamina Like, and Lamina: regions predicted to be associated with nuclear lamina. (c) Violin plots of intra-TAD ratio in GM12878 of lymphoma-related non-coding variants and non-lymphoma-related non-coding bins.

Nuclear compartments, such as nuclear speckles and lamina, participate in chromosome packaging and potentially associate with genome function [18, 51, 52]. Given the correlation between chromatin domain 3D structure and replication timing, we wondered whether chromatin domain 3D structure is also related to nuclear compartments. A recent study has identified several chromatin states (called Spatial Position Inference of the Nuclear genome, “SPIN” states) that correspond to nuclear compartmentalization [53]. We compared the intra-TAD ratios of genomic loci belonging to different SPIN states in human K562 myelogenous leukemia cells (Figure 5b). In general, the inactive and heterochromatic lamina and lamina-like regions have the highest intra-TAD ratios, significantly higher than the intra-TAD ratios of the active interior regions. It has been shown that an active inter-chromosomal interaction hub is organized around nuclear speckles [18]. We found that genomic regions near nuclear speckles have lower intra-TAD ratios than those close to the nuclear lamina but higher intra-TAD ratios than some active interior regions. This may be due to spatial constraints imposed on chromatin structures by the nuclear speckles, which require further investigations.

### 3.8 Chromatin domain 3D structure is related to disease-associated genetic variants

Beyond the correlation of chromatin domain 3D structures with the various molecular- and cellular-level features shown above, we further explored whether they are also related to higher-level phenotypes. Specifically, we studied the locations of disease-associated genetic variants relative to chromatin domain 3D structures. Genome-wide association studies (GWAS) have identified millions of variants associated with various diseases. A large proportion of these variants are outside coding sequences and tend to locate in the regulatory regions [54]. As such, they may tend to reside on domain surfaces in the relevant cell types. To test this hypothesis, we downloaded lymphoma-related variants from the GWAS Catalog [55]. We first found that these variants are strongly enriched in the cCREs in GM12878 (2.84 fold of enrichment, *p* = 1.60 *×* 10*^−^*^35^, hypergeometric test), as expected. We then specifically focused on the lymphoma-associated non-coding variants, and found that they have significantly smaller intra-TAD ratios as compared to all other non-coding regions (Figure 5c), indicating that indeed these variants tend to locate on domain surfaces in this B lymphocyte cell line.

## 4 Discussion

In this study, we have shown that the intra-TAD ratio serves as a proxy for chromatin domain 3D structures. It enabled us to discover various molecular features that are correlated with chromatin domain 3D structures, including the enrichment of highly expressed genes, CREs, active histone marks, and TF binding sites on domain surfaces. Most of these correlations are cell type-specific and are more prominent in the B2 and B3 sub-compartments, which contain larger and more inactive TADs. We have also shown that genomic loci with different replication timing and near different nuclear compartments have different preferences of their localization relative to chromatin domain 3D structures. Finally, we have also shown that disease-associated non-coding variants tend to reside more on domain surfaces.

The plots of gene expression against intra-TAD ratio (Figures 2a, S14a) show that there is a clear depletion of genes with both a high intra-TAD ratio and a high gene expression level. On the other hand, genes with a low intra-TAD ratio are not guaranteed to have a high expression level. This relationship resembles findings about promoter DNA methylation in gene expression control: whereas a gene with high promoter methylation is repressed, a gene with low promoter methylation is permitted to express but its expression level depends on other factors such as transcription factor binding [56]. Analogously, a low intra-TAD ratio appears to be a condition permissive for a gene to be highly expressed, but whether it actually has a high expression level depends on additional factors.

The strong enrichment of cell type-specific CREs and TF binding sites on domain surfaces led us to propose the “domain surface CRE” model, in which CREs and genes are all on domain surfaces, which make it easy for them to interact with each other and with transcription factors coming from outside the TAD. Indeed, we found that in general most TFs have their binding sites enriched on domain surfaces, although variations were observed across chromatin subcompartments.

One question associated with the “domain surface CRE” model is why CREs tend to regulate genes in the same TAD, given that they are also exposed to genes outside (Figure 4a). One possibility is that genes in the same TAD as a CRE have more stable proximity with it, while the proximity between the CRE and genes outside the TAD could be more dynamic. As a result, a CRE can occasionally get into contact with genes outside its TAD when proximity requirements are satisfied, resulting in the incomplete insulation of CREs [27].

In this study, we have demonstrated the use of multiplex data from SPRITE to study chromatin domain 3D structures, which offer both semi-single-cell information (by having each cluster expected to come from a single cell but different clusters in the same data set coming from a population of cells) and longer-range interactions that are not captured by the proximity ligation of Hi-C. The fairly strong correlation between the intra-TAD ratios computed from SPRITE and bulk Hi-C suggest that chromatin domain 3D structures are reasonably stable across single cells and that binary interactions from Hi-C could be sufficient for computing the intra-TAD ratio. This opens the opportunity for large-scale study of intra-TAD ratios in different cell types, given the much more abundant published bulk Hi-C data than multiplex data. On the other hand, to further investigate the stability of chromatin domain 3D structures across single cells, single-cell SPRITE data [57] could be utilized. By comparing intra-TAD ratios computed from bulk Hi-C, scHi-C, SPRITE, and single-cell SPRITE, differences in intra-TAD ratios caused by cell-cell variation and those caused by multiplexing can be separately analyzed.

A common limitation of all these “all-against-all” sequencing methods is that due to the very large number of chromosome interactions genome-wide, it is practically impossible to achieve sub-kilobase data resolution even with very deep sequencing. Instead, by using digitonin cell permeabilization, micrococcal nuclease (MNase) digestion, and direct sequencing of ligation junctions, it is possible to probe interactions of distinct viewpoints (the Micro Capture-C method [58]) or tiled regions (the Tiled-MCC method [59]) at near-base pair resolution, which helps explore nanoscale regulatory interactions. Based on the data produced from these new methods so far, it has been found that active loop extrusion is not essential for enhancer-promoter interactions but it contributes to their specificity and robustness, while local insulation of interactions among regulatory elements depends on CTCF [59]. We hypothesize that these findings are related to scDomains, whose boundary locations and correspondingly 3D structures are partially specified by loop extrusion and CTCF binding sites, and CTCF binding can create physical barriers between nearby regions in the 3D space. Further investigations are required to elucidate these mechanisms in detail.

Our analysis of lymphoma-associated non-coding variants shows that there is a trend for them to reside on domain surfaces. A difficulty of this analysis was the small number of variants available from the GWAS Catalog, which prohibited us from further focusing on only the non-coding variants within GM12878 cCREs since that would leave too few variants to remain and thus decrease statistical power. Similarly, it would be interesting to investigate whether the coreness of a region relative to its chromatin domain provides non-redundant information about disease risk on top of its chromatin state, but the same issue with statistical power pertains. Nevertheless, our initial results here suggest that similar analyses can also be performed for other diseases and corresponding cell types. By pooling data from related diseases, such as a pan-cancer analysis, more significant enrichment of disease-associated non-coding variants on domain surfaces may be found.

In some previous work, efforts have been devoted to deriving useful 1D features (i.e., a single numeric value for each genomic locus) from Hi-C contact maps and to linking them with chromatin organization and gene regulation [60]. As the most well-known examples, the first principal component of the normalized Hi-C contact map was introduced to partition genomic loci into the A and B compartments [15], and both the directionality index (DI) [20] and insulation score (IS) [61] were used to identify TAD boundaries. The intra-TAD ratio we introduce in this work can also be considered a type of 1D feature. The previously proposed open chromatin index (OCI) [62] and distal-to-local ratio (DLR) [63] bear some similarity to the intra-TAD ratio, in that they are both based on the comparison of Hi-C interactions that involve proximal regions versus distal regions. Yet to the best of our knowledge, the current study is the first one that uses 1D features derived from Hi-C contact maps to study chromatin domain 3D structures.

### 4.1 Limitations of the study

There are several limitations of this study. First, although the whole-chromosome tracing data set has enabled us to study scDomain structures, it only provides high-resolution data (at 50kb resolution) for one chromosome, while the whole-genome data from the same study do not have the resolution required by our analyses. Second, data involved in our study of chromatin domain structures (imaging, SPRITE, bulk Hi-C, and scHi-C) were only obtained from the same cell types as the functional data but not the same single cells. We were therefore not able to directly relate chromatin domain structures and functional signals at the single cell level. Third, all the data we have used in this study were obtained from cell lines, which may differ from *in vivo* situations. Fourth, the structural models we used were originally derived from a limited number of cells, and thus may not have captured sufficient information about cell-to-cell variability. Fifth, the different types of data we used all provide only a snapshot of the cells but do not provide information about dynamic changes over time. Sixth, the analyses performed for studying relationships between intra-TAD ratio and functional signals were correlative in nature. Causal relationships, such as whether change of location of a gene from the core to the surface of a scDomain would lead to increase of its expression, are yet to be confirmed. Finally, some of the correlations between the intra-TAD ratio and chromatin signals are weak. While the binding sites of several proteins that associate with specific histone marks (CBX3, EZH2, and KAT2A) have provided supports for the biological relevance of these correlations, a more systematic analysis involving binding sites of more histone writers would be required.

## 5 Methods

### 5.1 Quantifying the location tendencies of scDomain boundaries

High-resolution imaging data of 50kb genomic bins in chromosome 21 of IMR90 cells produced by Su et al. [13] were downloaded from Zenodo at https://doi.org/10.5281/zenodo. 3928890. Processed bulk Hi-C data produced by Rao et al. [16] were also downloaded from the same link. scDomains were identified from the imaging data following the code provided by Su et al. [13], available at https://github.com/ZhuangLab/Chromatin_Analysis_2020_cell. Briefly, pairwise distances between all pairs of 50kb bins in each individual cell were calculated from the imaging data. For each 50kb bin in each cell, upstream and downstream loci within a fixed window size were selected to calculate the insulation score. With this sliding window calculation, loci exhibiting local maxima of insulation scores were identified as potential domain boundaries. Boundaries that separated domains exhibiting similar patterns were then removed.

TADs called from the bulk Hi-C data were downloaded from the 3D Genome Browser [64] at http://3dgenome.fsm.northwestern.edu/. For each pair of 50kb bins separated by a certain genomic distance, we divided them into two groups based on whether they belonged to the same TAD or not. In each group, we counted the number of cells in which the two bins belonged to the same scDomain.

### 5.2 Quantifying the 3D structural tendencies of scDomains

For each 50kb bin in chromosome 21 of an IMR90 cell, we defined its spatial neighborhood as a sphere centered at this bin (based on its coordinates from the imaging data) with a radius of 500nm, which is the distance threshold that provides the best correlation with Hi-C contact data [13]. We then calculated the intra-scDomain of the bin as the fraction of bins in its neighborhood that come from its scDomain.

Using the Hi-C contact matrix of chromosome 21 in IMR90 provided by Su et al. [13], the intra-TAD ratio of each 50kb bin was calculated as the total intra-TAD contacts divided by the total contacts involving this bin.

To compare the intra-scDomain ratio computed from imaging data with the intra-TAD ratio computed from bulk Hi-C data, we randomly sampled a certain number of cells, among all single cells imaged, and took the average of the intra-scDomain ratio of each 50kb bin in these sampled cells. The resulting vector of average intra-scDomain ratios was used to compute Pearson correlation and Spearman correlation with the vector of intra-TAD ratios. For each number of cells, we repeated the random sampling independently 100 times to obtain a distribution of correlation values.

We also created shuffled TADs with the same size (i.e., 1D span) distribution and inter-TAD gap pattern as the actual TADs. This was done by first collecting the real sizes of the TADs and the gaps between them and putting them in a list following their original order. We then randomly permuted the list of TAD sizes and gap sizes, together with the label of whether the value is a TAD or a gap. We repeated this random permutation 100 times to produce 100 sets of shuffled TAD locations. We then used these shuffled TAD locations to compute intra-TAD ratios and calculated their correlations with average intra-scDomain ratios (over 200 randomly sampled cells), in the same way that we did with the real TADs. With the 3D coordinates of 50kb genomic bins in each scDomain, we identified the convex hull for each scDomain using the ConvexHull function implemented in the scipy.spatial module of Python [65]. We then compared the intra-scDomain ratio of the 50kb genomic bins forming the convex hull (“vertices”) with other bins (“non-vertices”).

### 5.3 Calculation of intra-TAD ratio using SPRITE data

SPRITE multi-contact read clusters from GM12878 cells and mESCs were downloaded from the 4D Nucleome data portal (https://data.4dnucleome.org/) [66] (data processed in GRCh38 and GRCm38) and Gene Expression Omnibus (http://www.ncbi.nlm.nih.gov/geo/) [67] (GEO; accession: GSE114242) (data processed in GRCh37). All sequencing reads within the read clusters were overlapped with genomic bins of a fixed size (5kb, 10kb, 25kb, or 50kb). TAD calls were downloaded from the 3D Genome Browser (http://3dgenome.fsm.northwestern.edu/) [64]. For a genomic bin *i* involved in a read cluster *j*, its normalized intra-TAD ratio in the read cluster, *r_ij_*, was defined as *N_ij_/*(*C_j_ −* 1), where *N_ij_* is the number of intra-TAD interactions between bin *i* and other bins involved in this cluster and *C_j_* is the total number of bins in cluster *j*. The overall intra-TAD ratio of bin *i*, *r_i_*, is then the average of all its intra-TAD ratios in the read clusters that it is involved in. For calculations that considered only intra-chromosomal interactions, we first partitioned the reads in each cluster into sub-clusters according to their chromosomes, and then computed intra-TAD ratios of each sub-cluster separately.

In order to compare with intra-TAD ratios calculated from bulk Hi-C data, scHi-C data, and structural models, which were processed using different reference genome builds in the case of GM12878, we computed intra-TAD ratios using SPRITE data based on both GRCh38 and GRCh37. In the case of mESC, all calculations were based on GRCm38.

In the analyses that involved classifying bins into “domain surface bins” and “domain core bins”, domain surface bins were defined in each TAD separately as bins with an intra-TAD ratio smaller than or equal to the *q_k_*-th quantile of intra-TAD ratio within TAD *k*, where 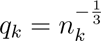 and *n_k_* is the total number of bins in TAD *k*. This definition of *q_k_* is based on the ratio between surface area (*S*) and volume (*V*) of a sphere, which is proportional to 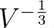. The remaining TAD regions were defined as domain core bins.

### 5.4 Pre-processing of SPRITE data when computing intra-TAD ratios

Since SPRITE clusters involving too few or too many reads are potentially problematic, we tested different ranges of cluster size (i.e., number of reads per cluster) following the original SPRITE paper [18], and found 2-1,000 to be a reasonable range that provides a balance between filtering out problematic clusters and maintaining the diversity of intra-TAD ratios (Figure S4). Previous studies have shown that inter-chromosomal interactions provide additional information about chromatin organizations [18, 44] and considering them led to more consistent intra-TAD ratios across different ranges of cluster sizes compared with using intra-chromosomal interactions alone to compute intra-TAD ratios (Figure S4). Considering these observations and the fact that intra-TAD ratios remain fairly stable across bin sizes (Figure S4), in our default setting for computing the intra-TAD ratio, we used a small bin size of 10kb to obtain fine-grained information and considered both intra-chromosomal and inter-chromosomal interactions based on SPRITE clusters with 2-1,000 reads. We also filtered out genomic bins with less than 100 SPRITE clusters involving them.

### 5.5 Calculation of intra-TAD ratio using bulk Hi-C data

Processed in situ Hi-C data (in .hic format) produced from GM12878 cells [16], K562 cells [16], and mESCs [41] were downloaded from the 4D Nucleome data portal (https://data.4dnucleome.org/). To extract whole-genome intra- and inter-chromosomal contact matrices, we first added SCALE normalization vectors to the original .hic data using the addNorm function of Juicer Tools (version 1.22.01). The contact matrices were generated by Juicer Tools (version 1.22.01) dump at 10kb, 25kb, and 50kb resolution. The intra-TAD ratio of each genomic bin was then calculated as the intra-TAD contacts divided by the total contacts involving this bin in a Hi-C contact matrix.

### 5.6 Calculation of intra-TAD ratio using single-cell Hi-C data

Single-cell Hi-C contacts from GM12878 cells and mESCs were downloaded from GEO (accessions: GSE117876 and GSE80280, respectively). The Hi-C contacts were converted to .pairs format (https://github.com/4dn-dcic/pairix/blob/master/pairs_format_specification.md#standard-format) if they were originally not. The intra-TAD contacts and total contacts of a given genomic bin were extracted using the Pyhton package pypairix (v0.3.7), and its intra-TAD ratio was then calculated as their ratio. We used 10kb bins for GM12878 scHi-C data and 50kb bins for mESC scHi-C data due to the sparsity of the latter data set. Finally, for each bin, the average intra-TAD ratio across all single cells was compared with its intra-TAD ratio computed using SPRITE data.

### 5.7 Calculation of intra-TAD ratio using structural models

Dip-C [34] structures at 20kb resolution for GM12878 cells were downloaded from GEO (accession: GSE117876) and Si-C [43] structures at 10kb resolution for mESCs were downloaded from https://github.com/TheMengLab/Si-C_3D_structure. To calculate the intra-TAD ratio of a genomic bin according to a 3D structure, we first found the *k* = 10 nearest neighbors of it in the structure based on their Euclidean distances using the Python package sklearn (v0.23.2). Among these *k* nearest neighbors, we counted the number of them belonging to the same TAD as the genomic bin of interest and divided it by *k* to become the intra-TAD ratio. For Dip-C structures, which also contained the diploid chromosome information, genomic bins were regarded as intra-TAD neighbors only if they were from the same TAD and the same chromosome haplotype. To obtain the intra-TAD ratio at 10kb resolution for the 20kb bins in Dip-C data, we separated each 20kb bin into two virtual 10kb bins and assigned the same intra-TAD ratio to them. Finally, for each genomic bin, its intra-TAD ratios over all the available structural models (including cells and configurations) were averaged and compared with its intra-TAD ratio computed using SPRITE data.

### 5.8 Discretization of a range of values into intervals

In order to illustrate the correlation between two variables *X* and *Y*, we divided all values of *X* into distinct intervals with approximately equal numbers of values in each interval. By plotting the distribution of *Y* within each interval, we could observe how *Y* changes as *X* increases.

### 5.9 Violin plots

We used the violinplot function inside the seaborn module of Python [68] to create violin plots. By default, it estimates the data distribution based on the data points observed, which could result in non-zero probability densities of values outside the observed maximum and minimum values. To avoid the impression that values were observed outside the actual observed range of values, we disabled such data extrapolation by setting the parameter cut=2.

### 5.10 RNA-seq data analysis

Processed RNA-seq data from GM12878 cells were downloaded from GEO (accessions: GSE88583 and GSE88627). GENCODE [69] (v24) was used to annotate the genes and extract TSSs.

Replicates were averaged and gene expression levels were quantified by fragments per kilobase of exons per million fragments (FPKM). Processed RNA-seq data from mESCs were downloaded from GEO (accession: GSE90277) and processed in a similar way with GENCODE (vM4) annotations.

### 5.11 ATAC-seq and histone ChIP-seq data analysis

Processed ATAC-seq data and ChIP-seq data for H2AFZ, H3K4me1, H3K4me2, H3K4me3, H3K9ac, H3K9me3, H3K27ac, H3K27me3, H3K36me3, H3K79me2, and H4K20me1 (in .bigwig format) from GM12878 cells were downloaded from the ENCODE data matrix (https://www.encodeproject.org/). For each type of data, the averaged signal (fold change over control) in each genomic bin was computed using bigWigAverageOverBed.

### 5.12 Enrichment of *cis*-regulatory elements

Candidate *cis*-regulatory elements (cCREs) in GM12878, HeLaS3, HepG2, IMR90, and K562 cells as well as cell type-agnostic cCREs were downloaded from ENCODE (https://www. encodeproject.org/). All cCREs were overlapped with genomic bins at 5kb, 10kb, 25kb, and 50kb resolution. Regions either with no cCRE labels or labeled with “Low-DNase” were filtered out since they are unlikely CREs, and the remaining cCREs were intersected with all TAD regions, domain surface regions, and domain core regions. Suppose the number of all TAD bins is *a*, the number of all domain surface bins is *b*, the number of all filtered cCRE bins within TAD regions is *c*, and the number of cCRE bins on domain surfaces is *d*, the domain surface enrichment fold was defined as 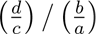. Enrichment *p*-values of cCREs on domain surfaces were calculated by fitting a hypergeometric distribution using the hypergeom.sf() function inside the scipy.stats module of Python [65]. Domain core enrichment folds and *p*-values were calculated similarly.

For each of the four categories of cCREs (dELS, pELS, PLS and CTCF-bound), a genomic bin was labeled with a certain category if it overlapped with a region of this subtype, regardless of its overlap with regions of other categories. To test the effect of TAD boundaries on surface cCRE enrichment, we excluded TAD boundary bins and the neighboring 10% bins of the whole TAD from the domain surface bins and repeated the analysis. To test cell-type specificity of surface cCRE enrichment, we excluded all cCREs defined in GM12878 from cell type-agnostic cCREs or cCREs in HeLaS3, HepG2, IMR90, or K562, and repeated the analysis.

### 5.13 Enrichment of transcription factors

ChIP-seq peaks of 153 transcription factors (TFs) obtained from GM12878 cells were downloaded from ENCODE (https://www.encodeproject.org/). All TF peaks were overlapped with genomic bins at 5kb, 10kb, 25kb, and 50kb resolution and intersected with domain surfaces and domain cores. Enrichment of TFs on domain surfaces and in domain cores was calculated in the same way as described above for cCREs.

The molecular weights of TFs were downloaded from UniProt (https://www.uniprot.org/ [70]). Only the “reviewed” entry was kept if a TF had multiple entries. For TFs with multiple “reviewed” entries, we manually selected one of them to define the molecular weight. We only considered each TF protein as a single sub-unit without considering other proteins in the possible TF complexes.

### 5.14 Repli-seq data analysis

Wavelet-smoothed signals of Repli-seq data (in .bigWig format) from GM12878 cells were downloaded from ENCODE (https://www.encodeproject.org/). Averaged signals in each genomic bin at 5kb, 10kb, 25kb, and 50kb resolution were obtained using bigWigAverageOverBed.

### 5.15 SPIN state analysis

SPIN annotations of genomic regions in K562 cells were downloaded from https://github.com/ma-compbio/SPIN [53]. Since SPRITE data were not available for K562 cells, intra-TAD ratios were calculated using bulk Hi-C data downloaded from the 4D Nucleome data portal (https://data.4dnucleome.org/ and processed in the same way for other Hi-C data, as described above.

### 5.16 Disease variants analysis

Lymphoma-related variants were downloaded from GWAS Catalog [55] with trait ID of “EFO 0000574”. The variants overlapped with any human transcript defined by GENCODE (v29) were excluded. The remaining variants were considered non-coding variants and were overlapped with 10kb bins. Intra-TAD ratios (calculated in GM12878) of these lymphoma-related non-coding variants were compared with those of all the non-coding 10kb bins.

## 6 Acknowledgment

We thank Aniruddha Deshpande, Genevìeve Fourel, Chiara Nicoletti, and Pier Lorenzo Puri for their comments on this work. This project was supported by Hong Kong Research Grants Council General Research Fund 14107420. QC is supported by National Natural Science Foundation of China Youth Program 32100515. KYY was additionally supported by Hong Kong Research Grants Council Collaborative Research Funds C4015-20E, C4045-18W, C4057-18E and C7044-19G, General Research Fund 14203119, and Theme-based Research Scheme T12-712/21-R, the Hong Kong Epigenomics Project (EpiHK), and the Chinese University of Hong Kong Young Researcher Award, Outstanding Fellowship and Project Impact Enhancement Fund. KYY is currently supported by the National Institute of Health P30 CA030199-41, U54 AG079758-01, and R21 AG075483-01S1.

## 7 Contributions

KYY conceived the study. KYL, QC and KYY developed the methodology and designed the analyses. KYL performed data processing and analyses. KYL, QC, HW, DL and KYY interpreted the results. KYL and KYY wrote the manuscript. All authors read and approved the final version of the manuscript.

## S1 Supplementary information

### S1 Supplementary figures

**Figure S1:**
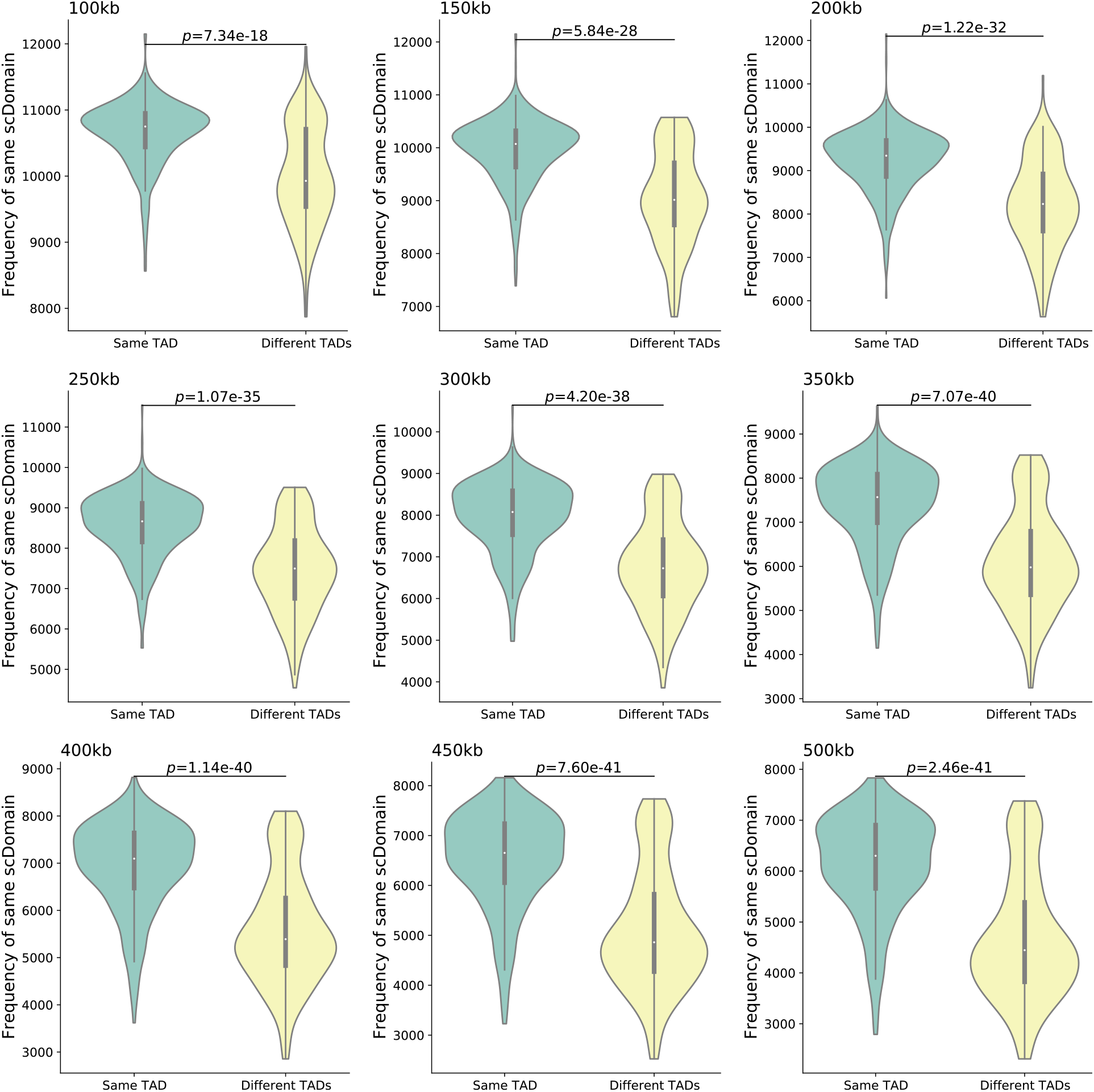
Consistency between scDomains and TADs of IMR90 cells. Each panel includes pairs of 50kb regions separated by a certain genomic distance (100kb-500kb). Each region pair is classified as either belonging to the same TAD or not. The frequency (i.e., number of cells among a total of *∼*12,000 cells) that the two regions belonging to the same scDomain is then counted. Finally, these frequency values from all region pairs are collected and visualized using violin plots. *p*-values are computed using the two-sided Mann–Whitney U test.

**Figure S2:**
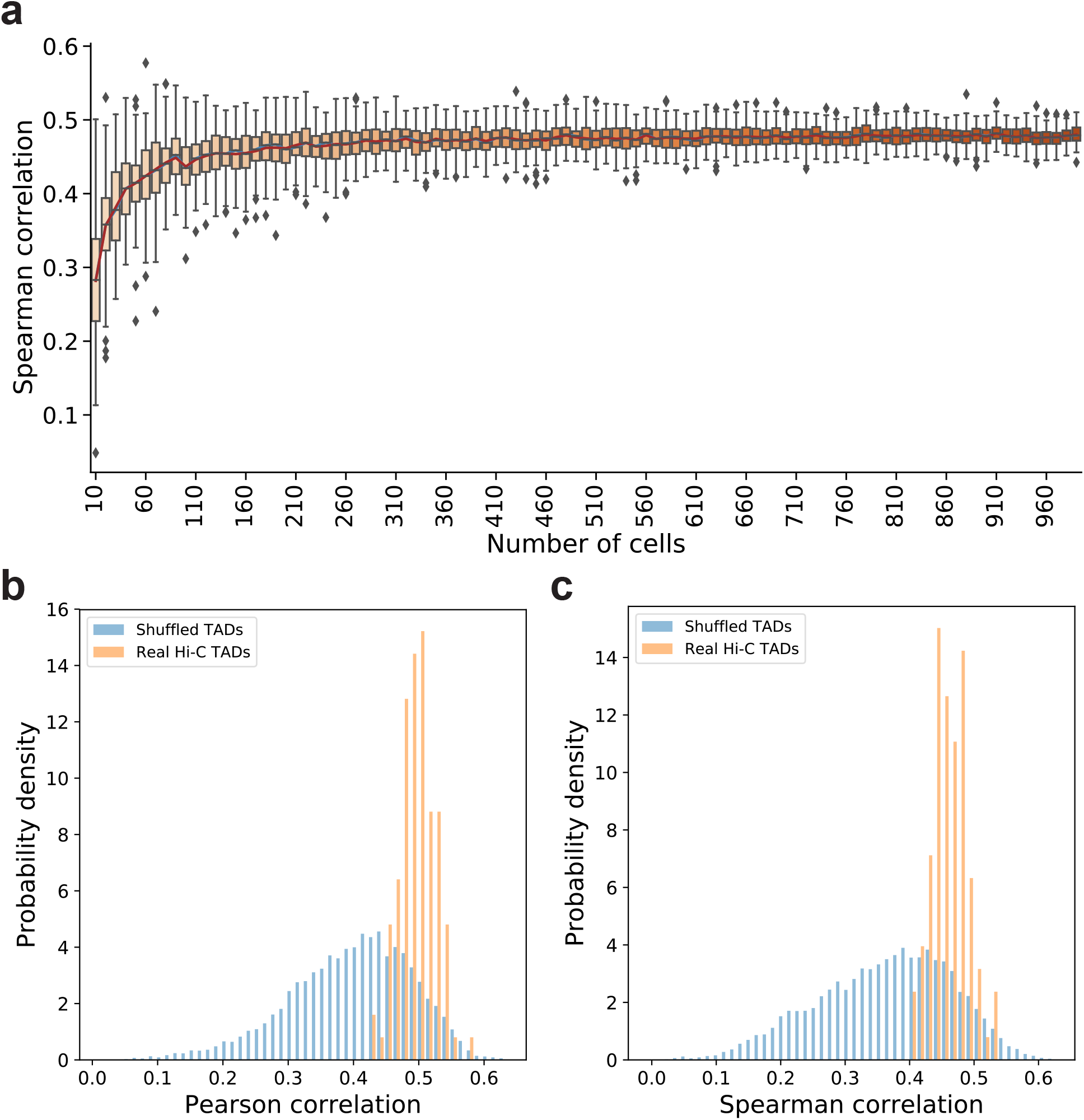
Consistency between average intra-scDomain ratios and intra-TAD ratios of IMR90 cells. (a) Distribution of Spearman correlations between intra-TAD ratio and average intra-scDomain ratio in *x* single cells, for *x* equal to 10-990. For each value of *x*, we first sub-sampled *x* single cells and computed the average intra-scDomain ratio of each 50kb region over these cells. Then the vector of these average intra-scDomain ratios and the corresponding vector of intra-TAD ratios of the same 50kb regions were taken to compute a correlation. Finally, by repeating the sub-sampling of *x* cells 100 times, we obtained a distribution of correlation values. Histogram of Pearson (b) and Spearman (c) correlations when considering *x* = 200 cells, based on real TADs or shuffled TADs.

**Figure S3:**
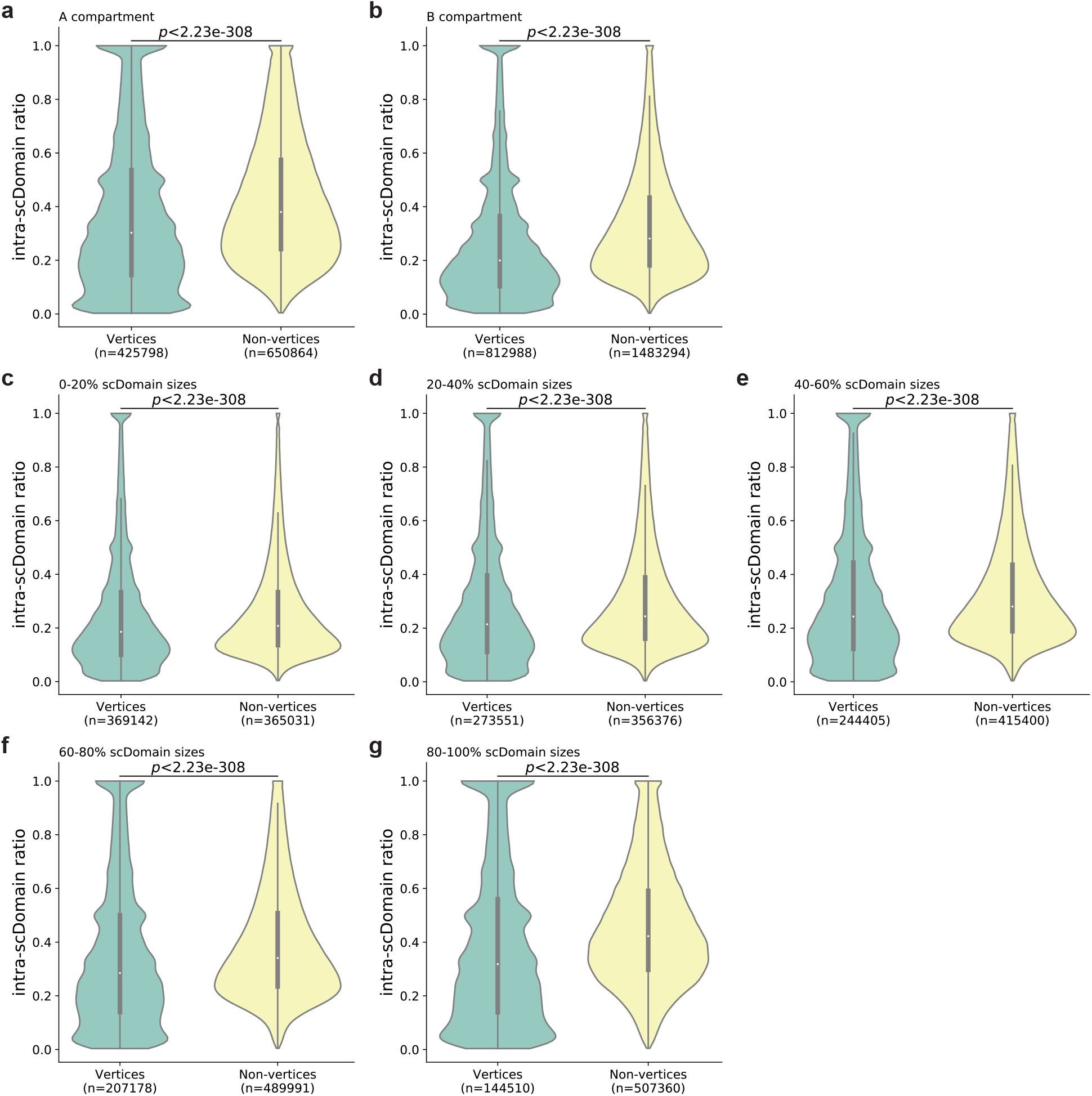
Comparing the intra-scDomain ratios of regions on the convex hulls of scDomains (vertices) and other regions (non-vertices). In each panel, a violin plot is shown for all included vertices from all imaged single cells, and a separate violin plot is shown for all included non-vertices from all imaged single cells. Different subsets of regions were included in the different panels, namely only regions in the A (a) and B (b) compartments, and only regions within a certain size range according to the number of constituent 50kb regions, namely 0-20 percentiles (c), 20-40 percentiles (d), 40-60 percentiles (e), 60-80 percentiles (f), and 80-100 percentiles (g), where a smaller percentile corresponds to scDomains with fewer constituent regions. *p*-values in all panels are computed using the two-sided Mann–Whitney U test.

**Figure S4:**
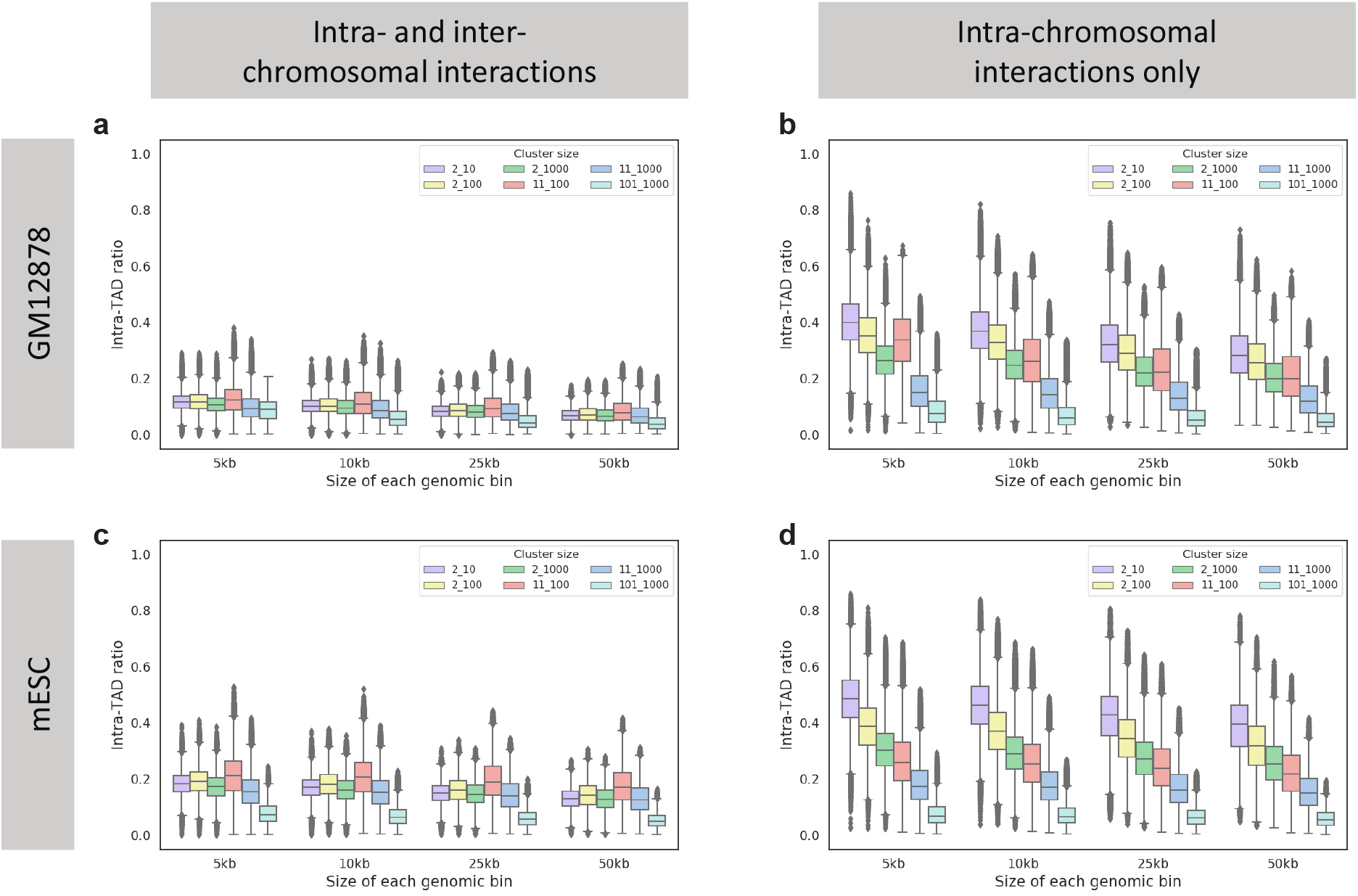
Effects of technical parameters on the intra-TAD ratio. The different panels show the intra-TAD ratios calculated using data from GM12878 cells (a,b) and mESCs (c,d), with both intra- and inter-chromosomal interactions (a,c) or only intra-chromosomal interactions (b,d). In each panel, the different groups show the intra-TAD ratios calculated with a specific size of genomic bins (5kb, 10kb, 25kb, or 50kb). Within a group, the different box plots correspond to different subsets of SPRITE clusters involved in the calculation of intra-TAD ratios selected based on their sizes, where *x y* means the inclusion of only SPRITE clusters with at least *x* and at most *y* reads.

**Figure S5:**
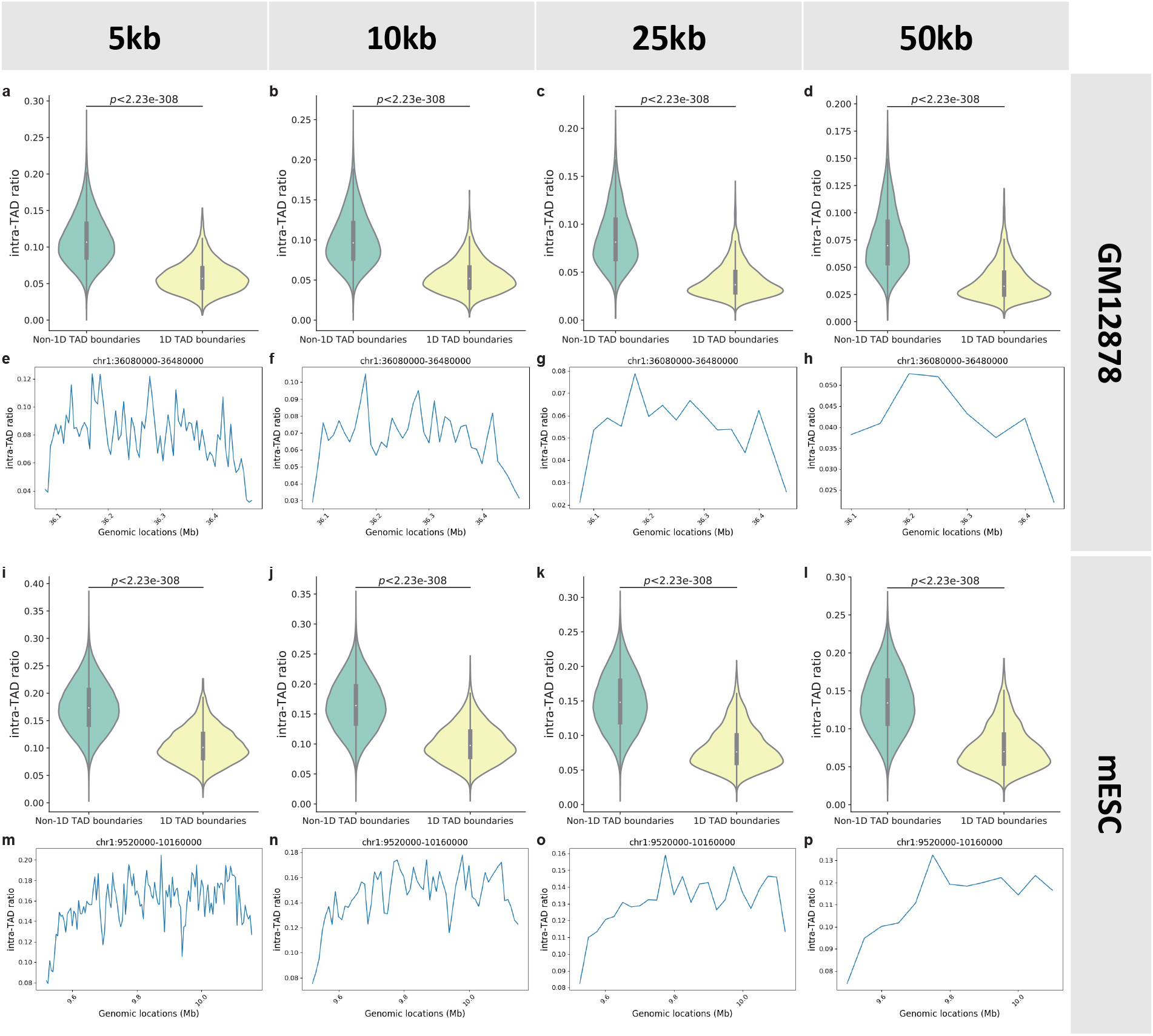
Comparing the intra-TAD ratios of 1D TAD boundaries and other TAD bins. The different panels show the intra-TAD ratios in GM12878 cells (a-h) and mESCs (i-p), at 5kb (a,e,i,m), 10kb (b,f,j,n), 25k (c,g,k,o), and 50kb (d,h,l,p) bin sizes. The violin plots (a-d, i-l) compare the distributions of intra-TAD ratios of genomic bins that overlap 1D TAD boundaries and those that do not. The line plots (e-h, m-p) show the change of intra-TAD ratio along the genomic span of example TADs.

**Figure S6:**
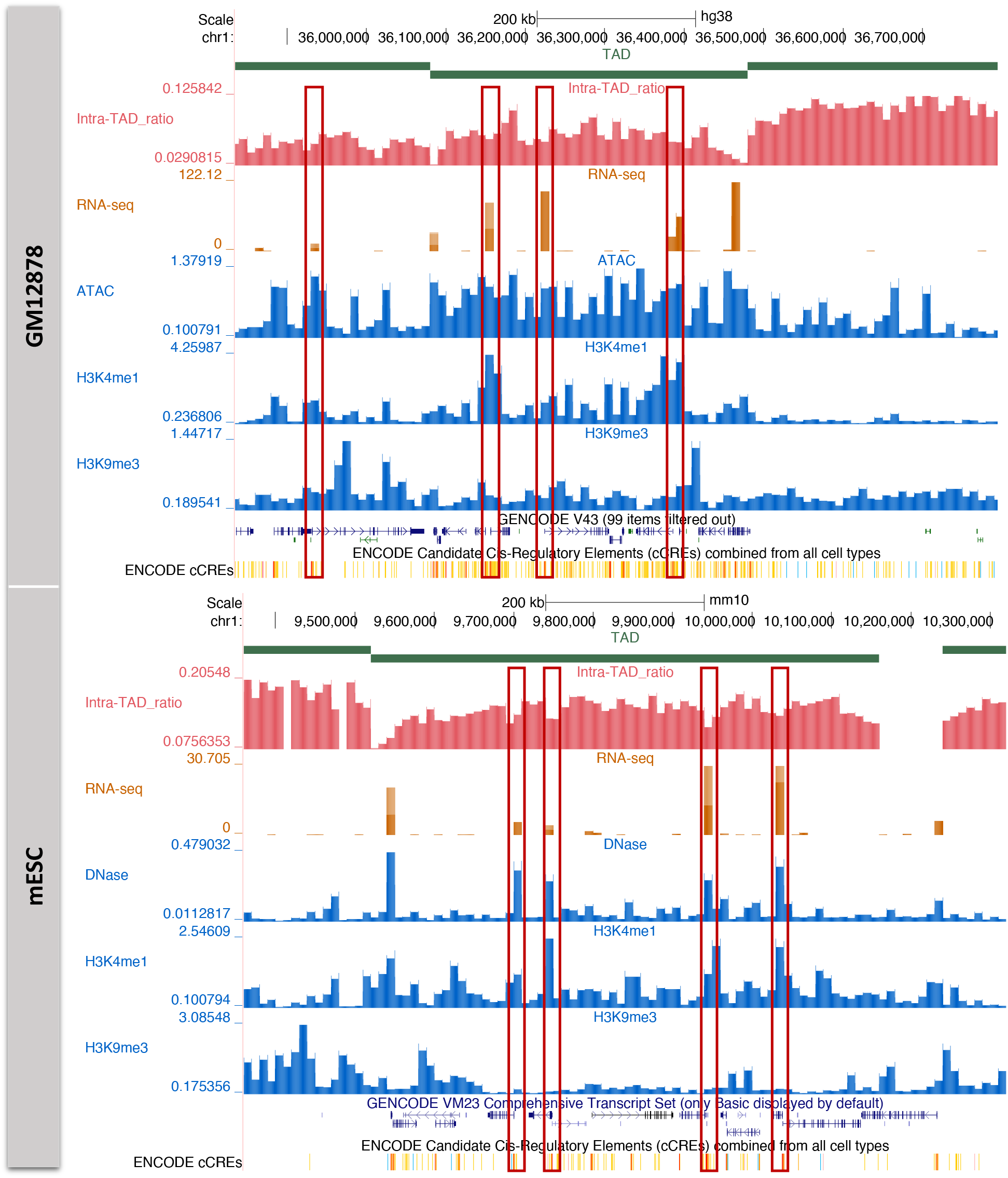
Genomic tracks showing TAD locations, intra-TAD ratios, RNA expression levels, signals of chromatin accessibility, H3K4me1, and H3K9me3, and gene annotation in GM12878 (a) and mESC (b). In each panel, the red boxes show example regions inside a TAD that are not close to 1D TAD boundaries but have a local dip of intra-TAD ratio, a corresponding high level of RNA expression, some enrichment of active chromatin signals and some depletion of inactive repressive chromatin signals.

**Figure S7:**
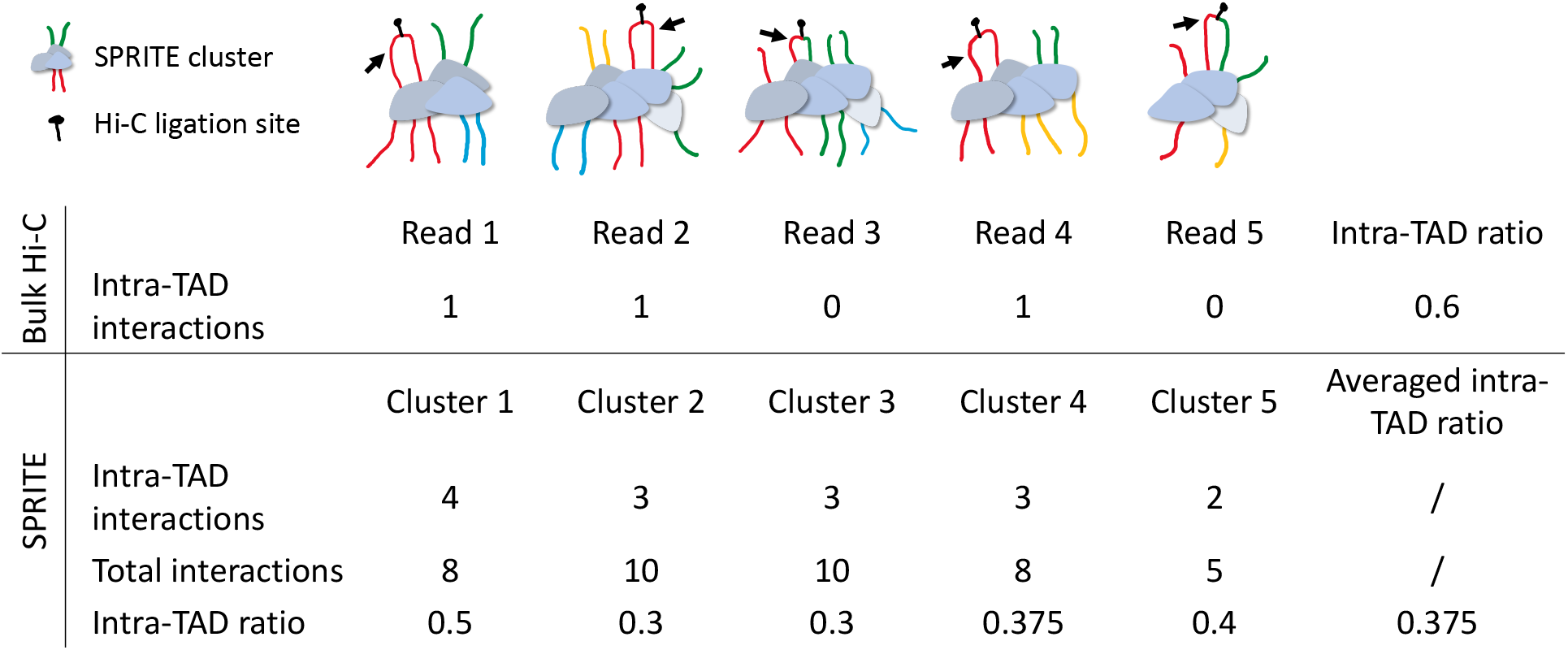
Calculations of intra-TAD ratio from SPRITE and bulk Hi-C data. Black arrows indicate the target DNA fragment of which the intra-TAD ratio is computed. DNA fragments from the same TAD are shown in the same color. Hi-C captures only one DNA fragment ligated to the target fragment at a time, while SPRITE captures the whole complex of multiple DNA fragments at the same time.

**Figure S8:**
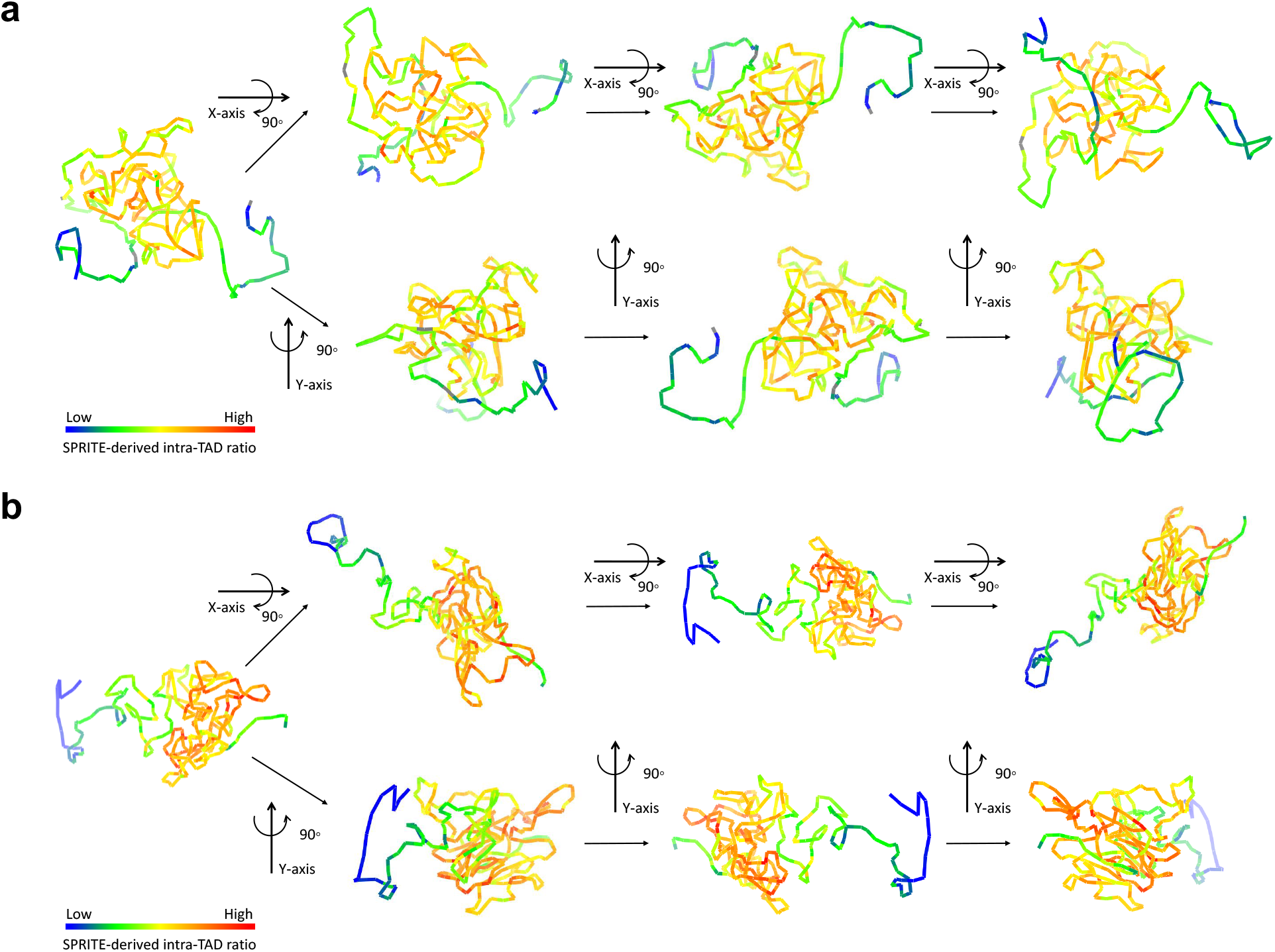
Visualization of intra-TAD ratios in structural models of two example TADs. The structural models of the first example TAD (chr18:77920000-80200000) (a) and the second example TAD (chr16:73000000-75560000) (b) were taken from Meng et al. [43]. Each genomic bin is colored by its SPRITE-derived intra-TAD ratio. Gray indicates missing data.

**Figure S9:**
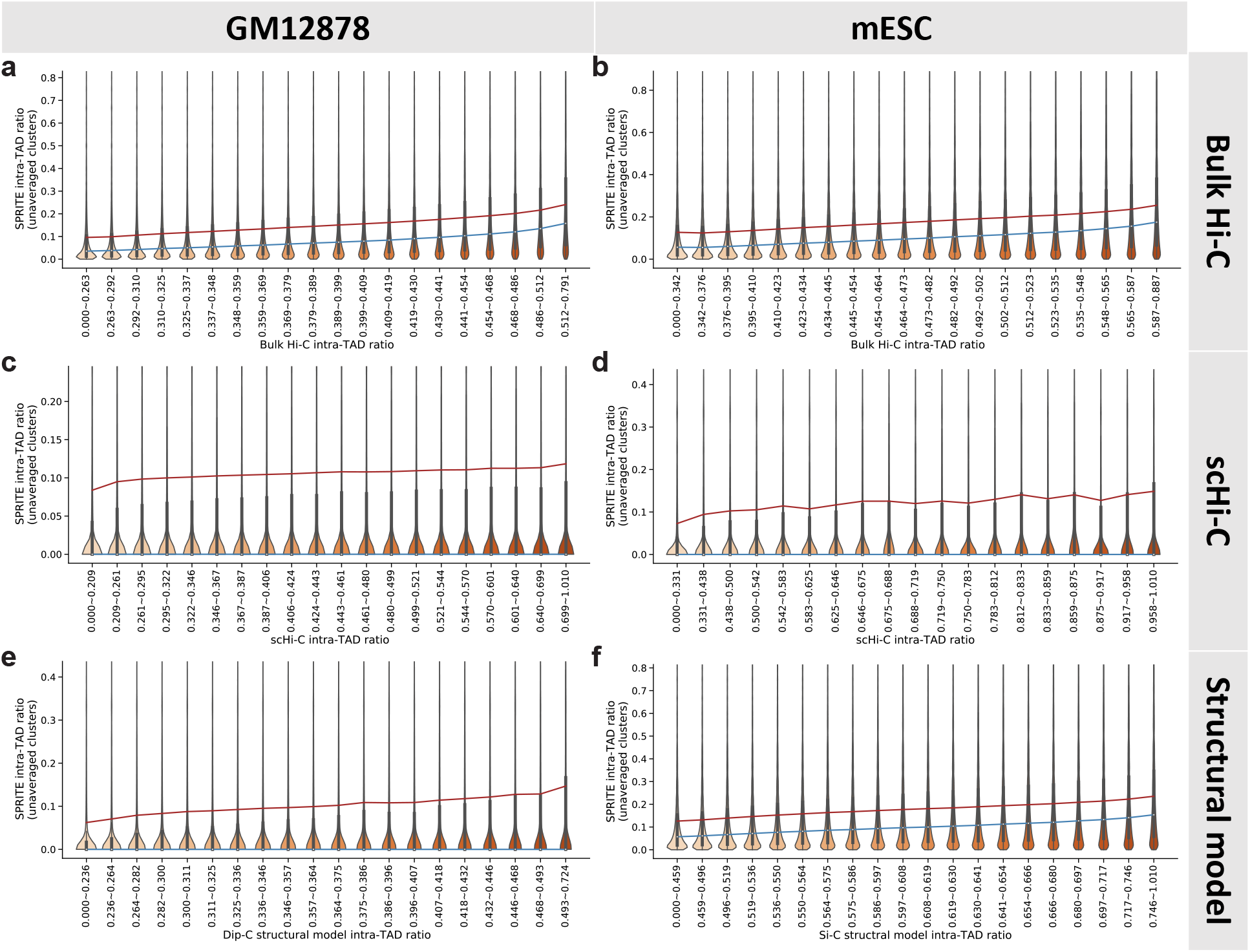
Comparing intra-TAD ratios calculated from SPRITE data without averaging over SPRITE clusters and intra-TAD ratios calculated from other types of data. Genomic bins were put into discrete classes based on their intra-TAD ratios computed from bulk Hi-C [16] (a,b), scHi-C [34, 42] (c,d), or structural models [34, 43] (e,f). The intra-TAD ratios computed from individual SPRITE clusters of the bins belonging to each class were then visualized using a violin plot. In all panels, the blue and red lines connect median and mean values of the different groups, respectively. PCC: Pearson correlation. SCC: Spearman correlation.

**Figure S10:**
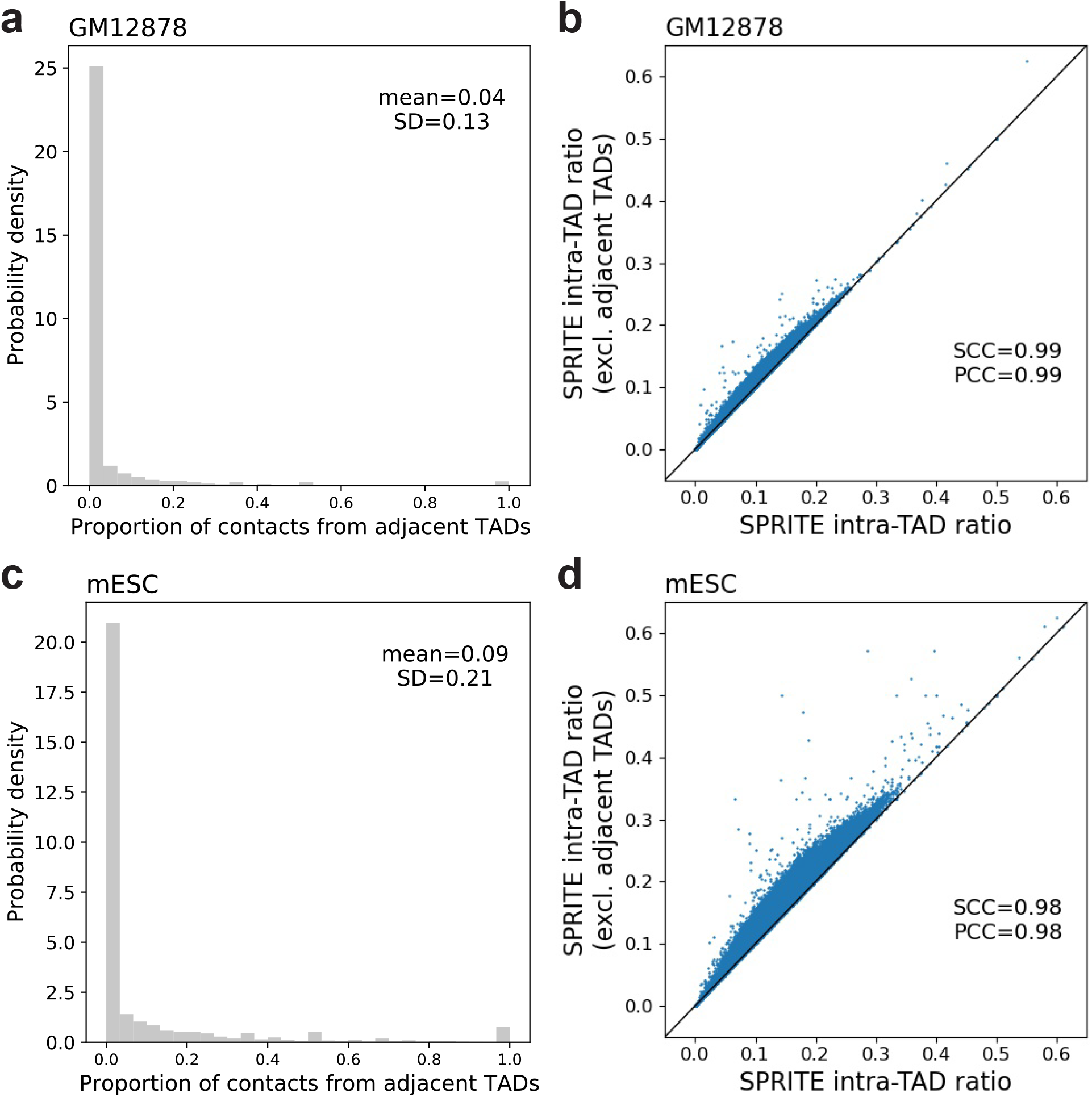
Effects of contacts from adjacent TADs in SPRITE data on the intra-TAD ratio. (a,c) Distribution of the proportion of contacts from adjacent TADs among all the inter-TAD contacts in GM12878 (a) and mESC (c). (b,d) Scatter plots of the intra-TAD ratio when interactions between adjacent TADs are excluded against the intra-TAD ratio when all the interactions are considered in GM12878 (b) and mESC (d).

**Figure S11:**
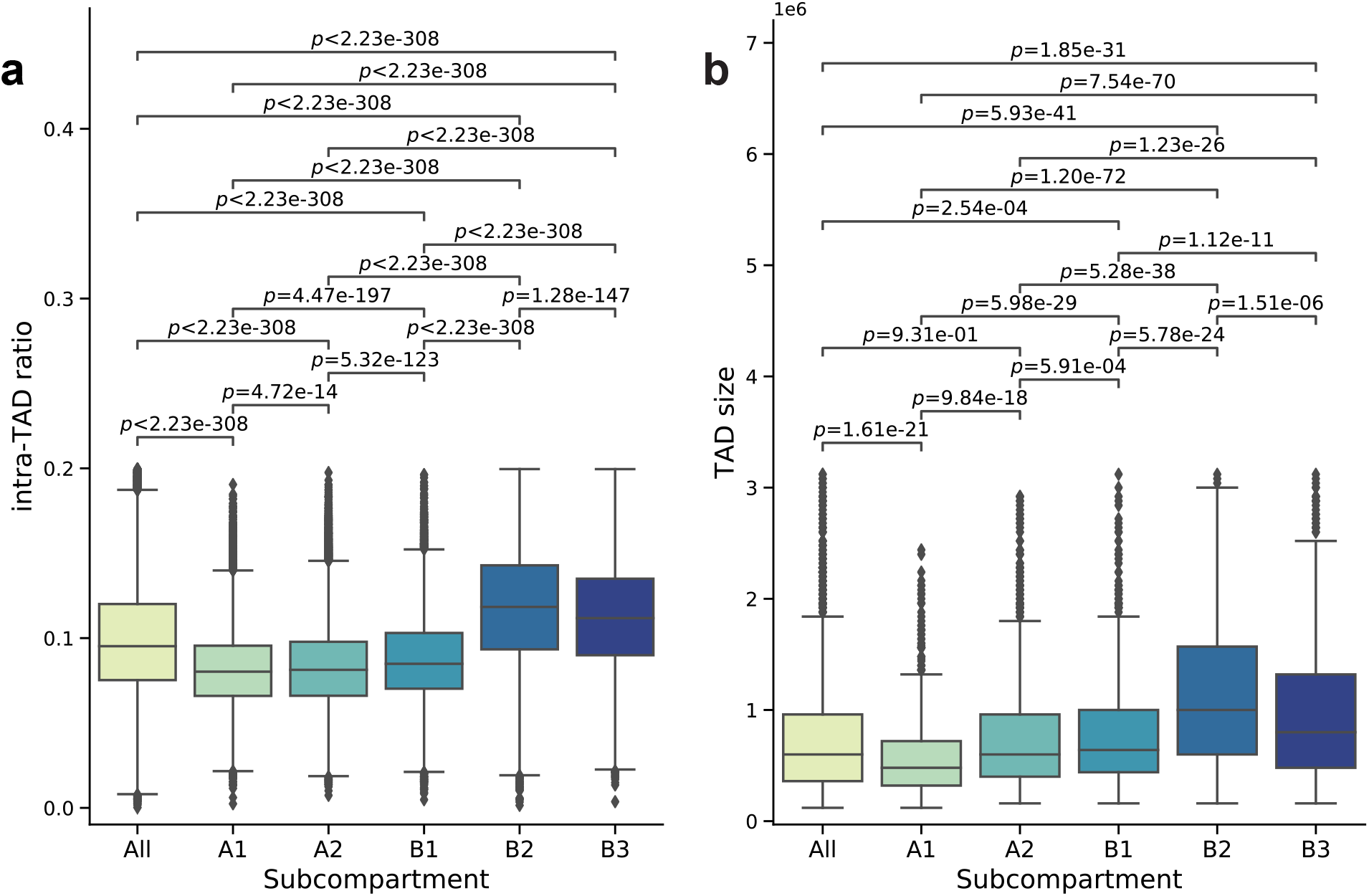
Distribution of intra-TAD ratio (a) and TAD size (b) of either all TADs or only TADs in a genomic subcompartment. The size of a TAD is defined as the length of its 1D genomic span.

**Figure S12:**
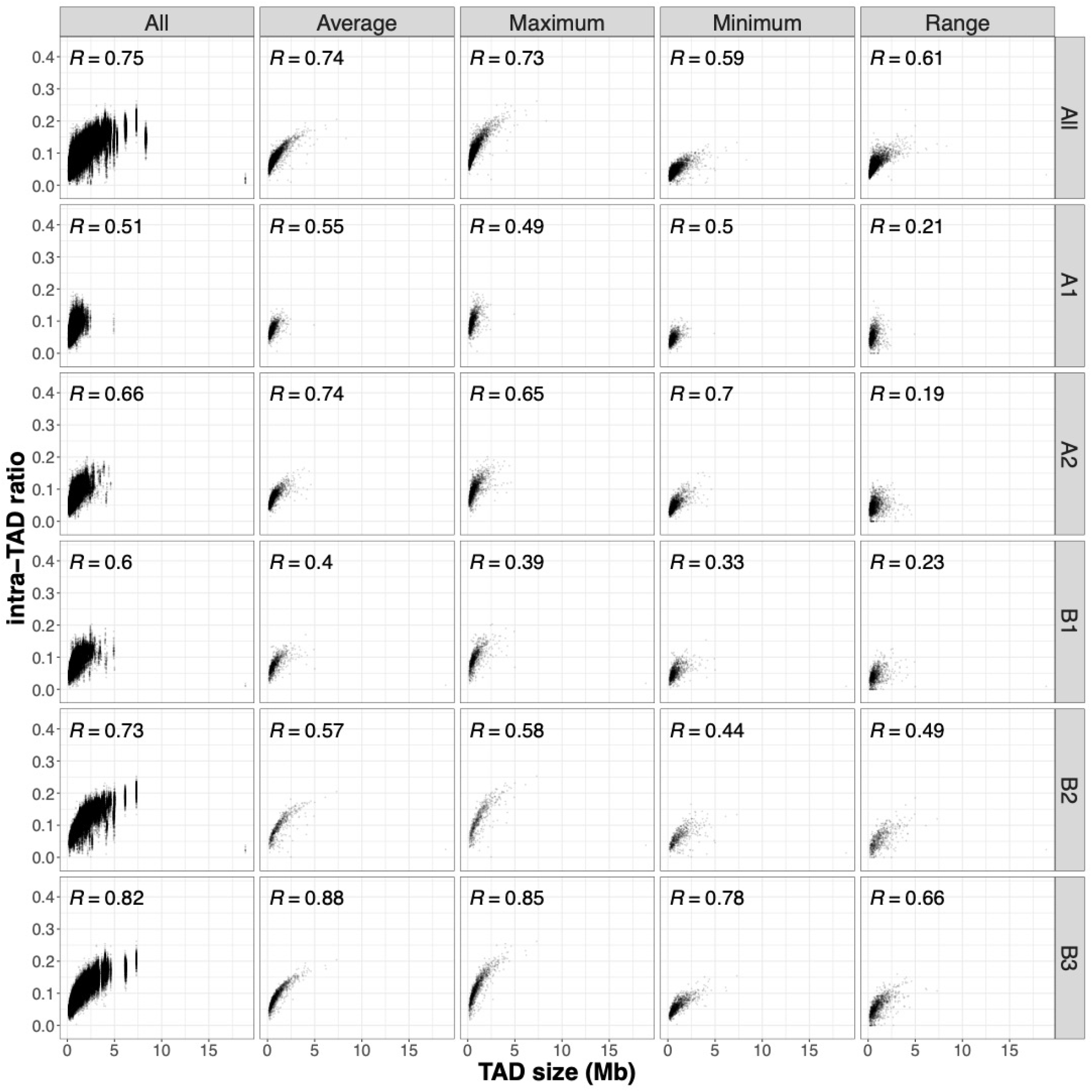
Relationship between intra-TAD ratio and TAD size. Each scatterplot compares TAD size with intra-TAD ratio based on either all TADs or only TADs in a genomic sub-compartment (rows) and a specific way to aggregate intra-TAD ratios of all genomic bins in a TAD (columns).

**Figure S13:**
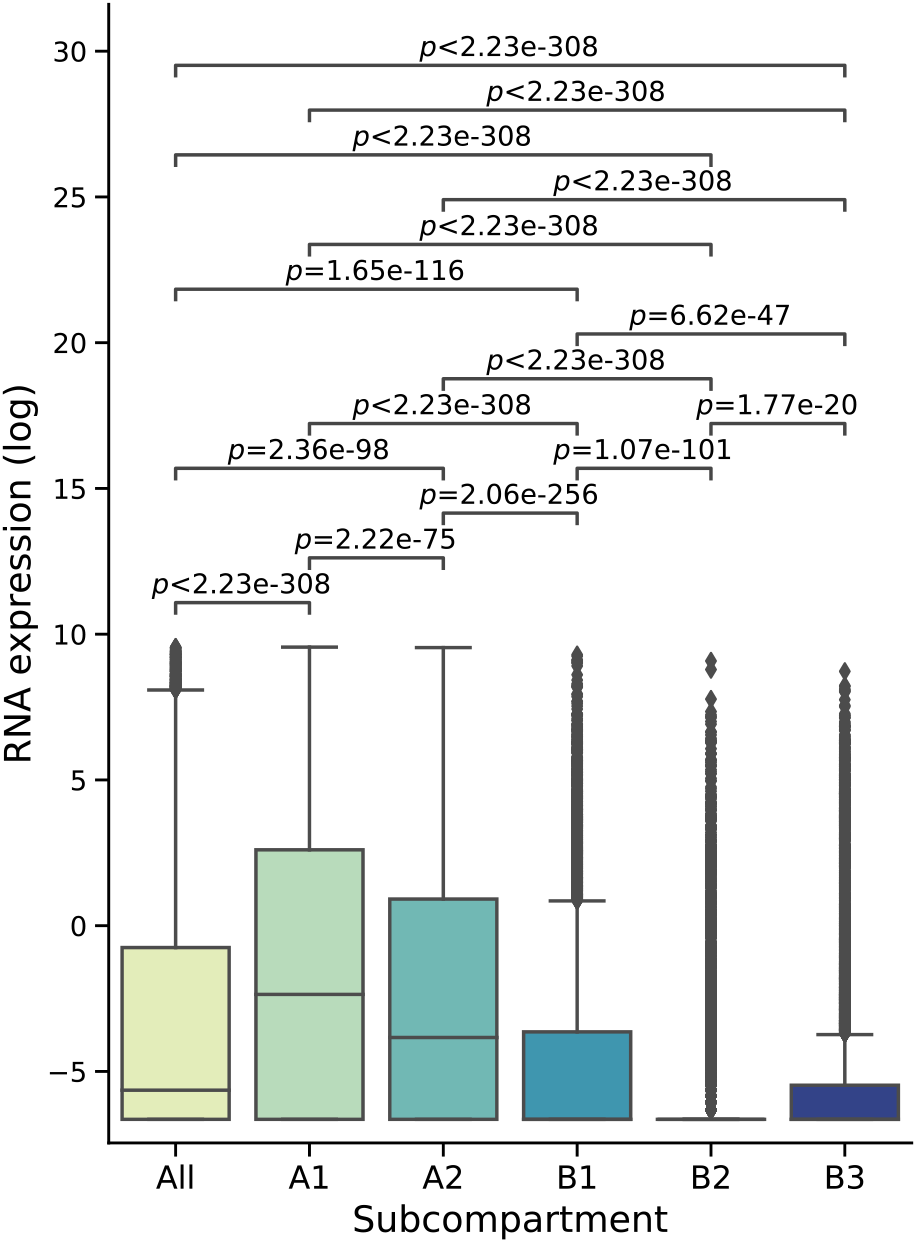
Distribution of expression levels of genes either in the whole genome or in a genomic subcompartment. RNA expression is quantified by log_2_(FPKM+0.01).

**Figure S14:**
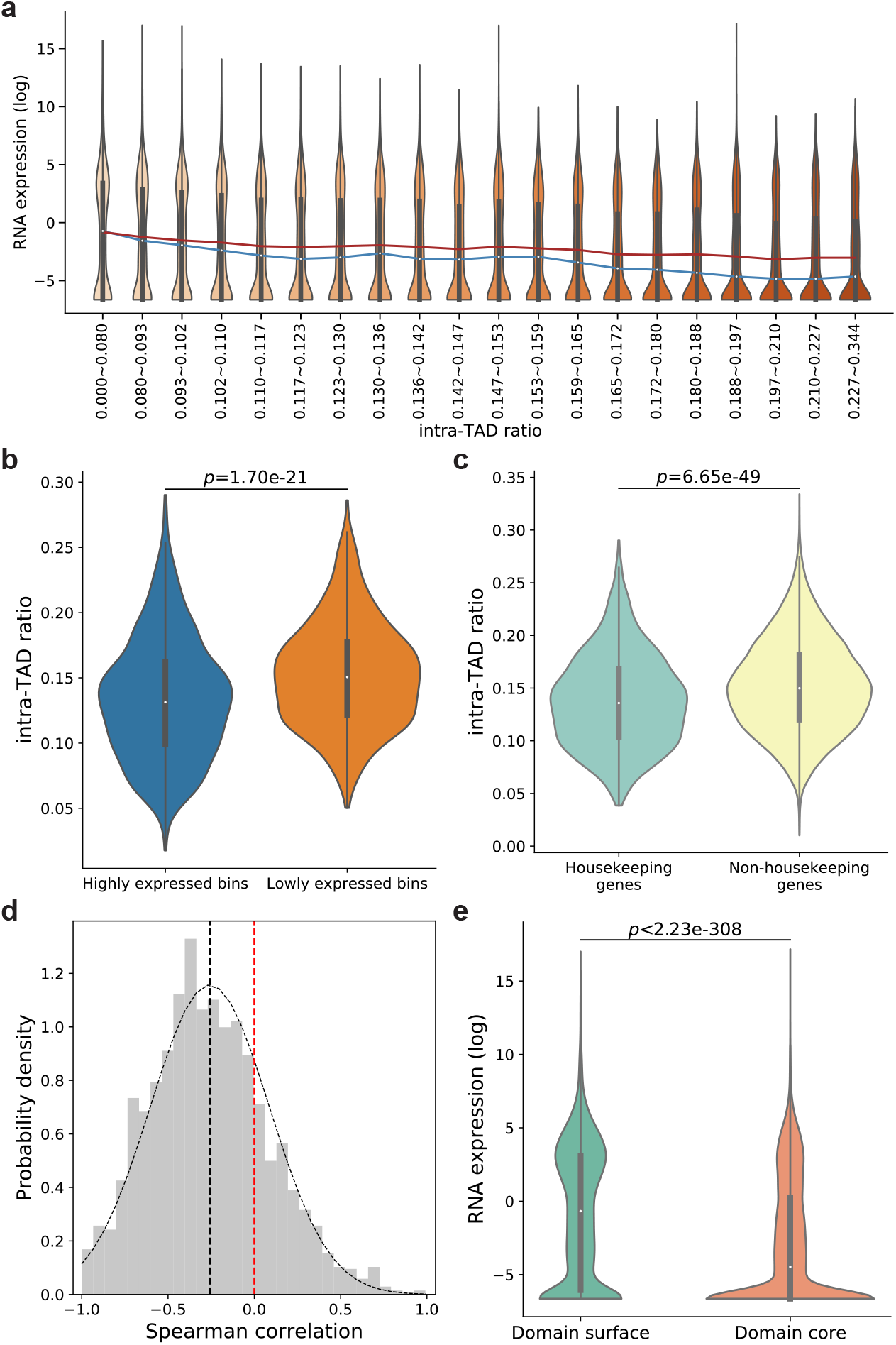
Inverse relationship between intra-TAD ratio and gene expression in mESCs. (a) Violin plots of gene expression (log_2_(FPKM+0.01)) of TSS bins in groups with increasing intra-TAD ratio and similar number of bins. The blue and red lines connect median and mean values of the different groups, respectively. (b) Violin plots of intra-TAD ratios of the highly expressed bins and lowly expressed bins, defined as the genomic bins containing TSSs of genes whose FPKM values rank within top and bottom 1,000, respectively. (c) Violin plots of intra-TAD ratio of housekeeing genes and non-housekeeping genes. (d) Distribution of Spearman correlations between gene expression and intra-TAD ratio of genomic bins in individual TADs. Black dotted curve shows the fitted normal distribution. The vertical black dotted line shows the mean of Spearman correlations. The vertical red dotted line shows zero Spearman correlation. (e) Violin plots of gene expression levels (log_2_(FPKM+0.01)) in domain surface bins and domain core bins. *p*-values in all panels are calculated using the two-sided Mann-Whitney U test.

**Figure S15:**
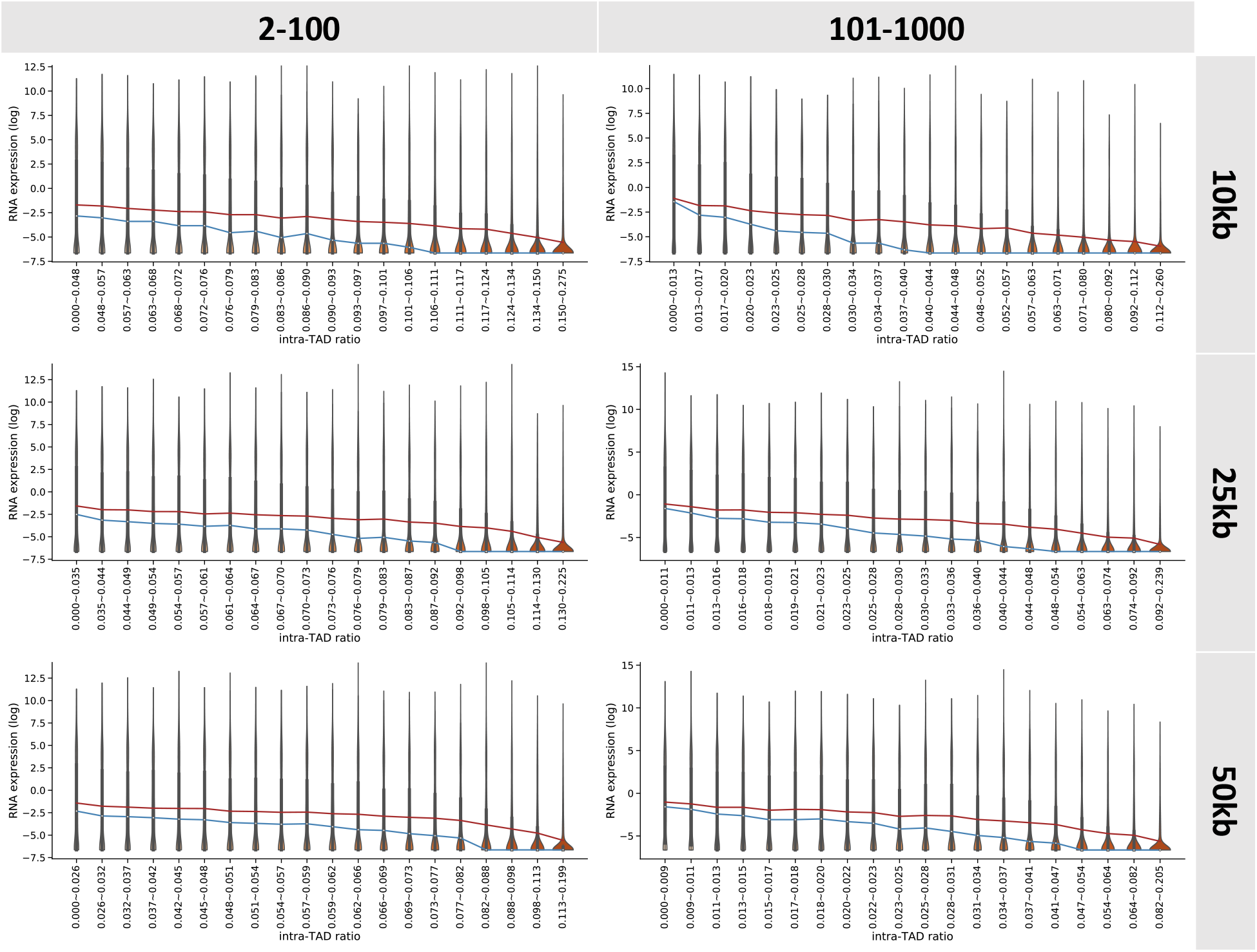
Inverse relationship between intra-TAD ratio and gene expression in GM12878 with different bin sizes (rows) and SPRITE cluster sizes (columns). Violin plots show gene expression (log_2_(FPKM+0.01)) of TSS bins in groups with increasing intra-TAD ratio and similar number of bins. The blue and red lines connect median and mean values of the different groups, respectively.

**Figure S16:**
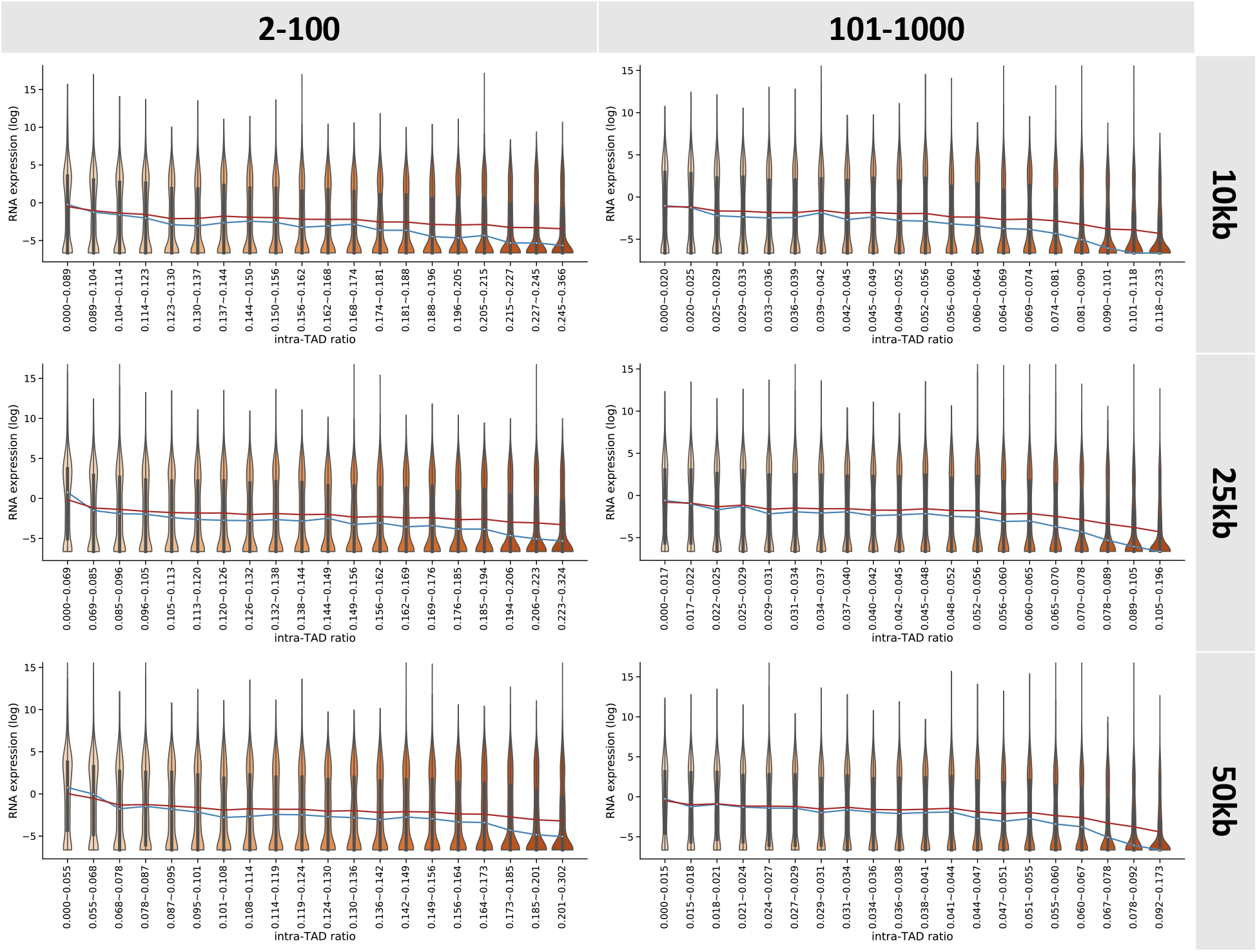
Inverse relationship between intra-TAD ratio and gene expression in mESC with different bin sizes (rows) and SPRITE cluster sizes (columns). Violin plots show gene expression (log_2_(FPKM+0.01)) of TSS bins in groups with increasing intra-TAD ratio and similar number of bins. The blue and red lines connect median and mean values of the different groups, respectively.

**Figure S17:**
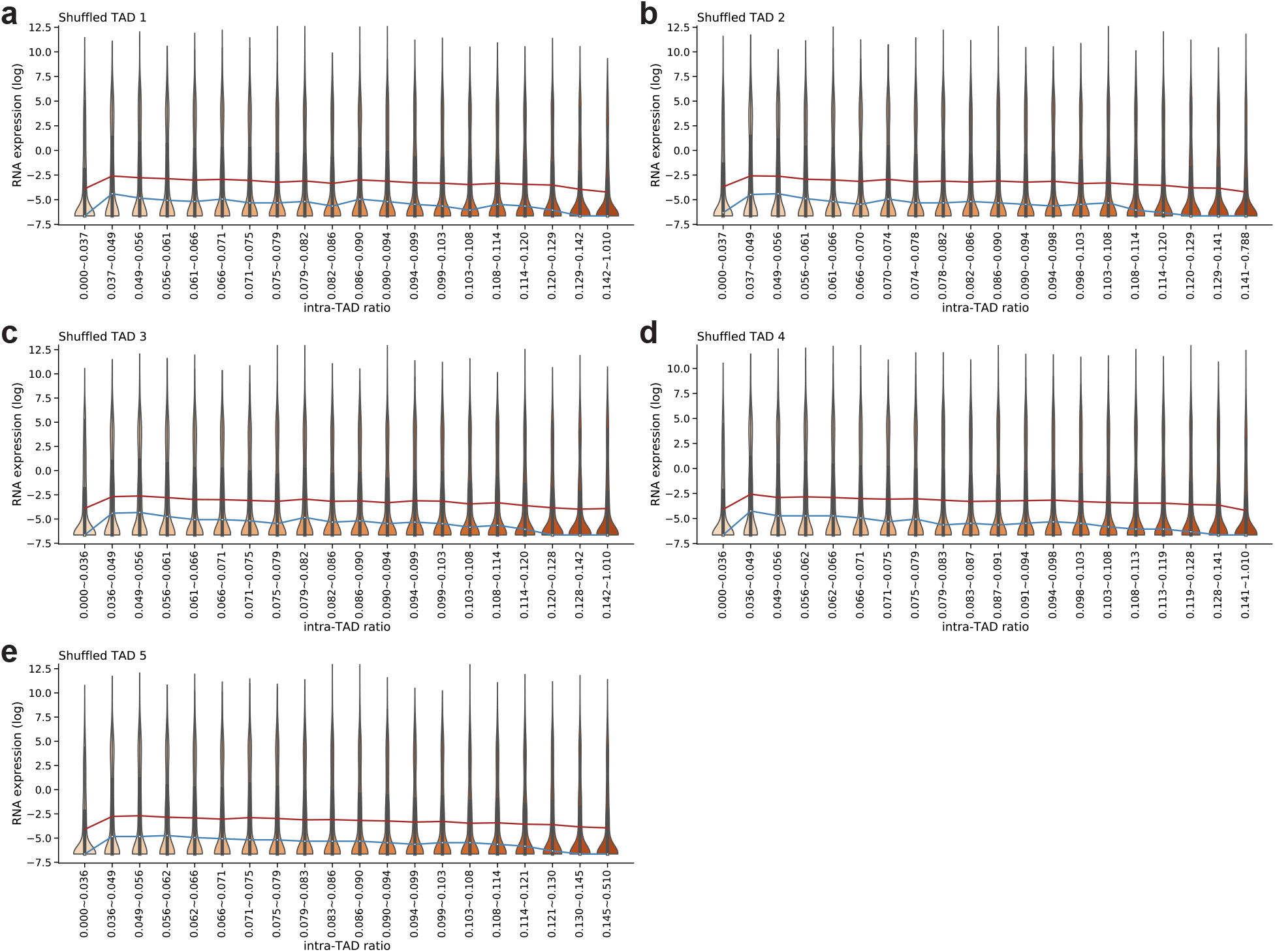
Relationship between intra-TAD ratio and gene expression for shuffled TADs in GM12878. Violin plots show gene expression (log_2_(FPKM+0.01)) of TSS bins in groups with increasing intra-TAD ratio and similar number of bins, with intra-TAD ratio being calculated using five sets of shuffled TADs. The blue and red lines connect median and mean values of the different groups, respectively.

**Figure S18:**
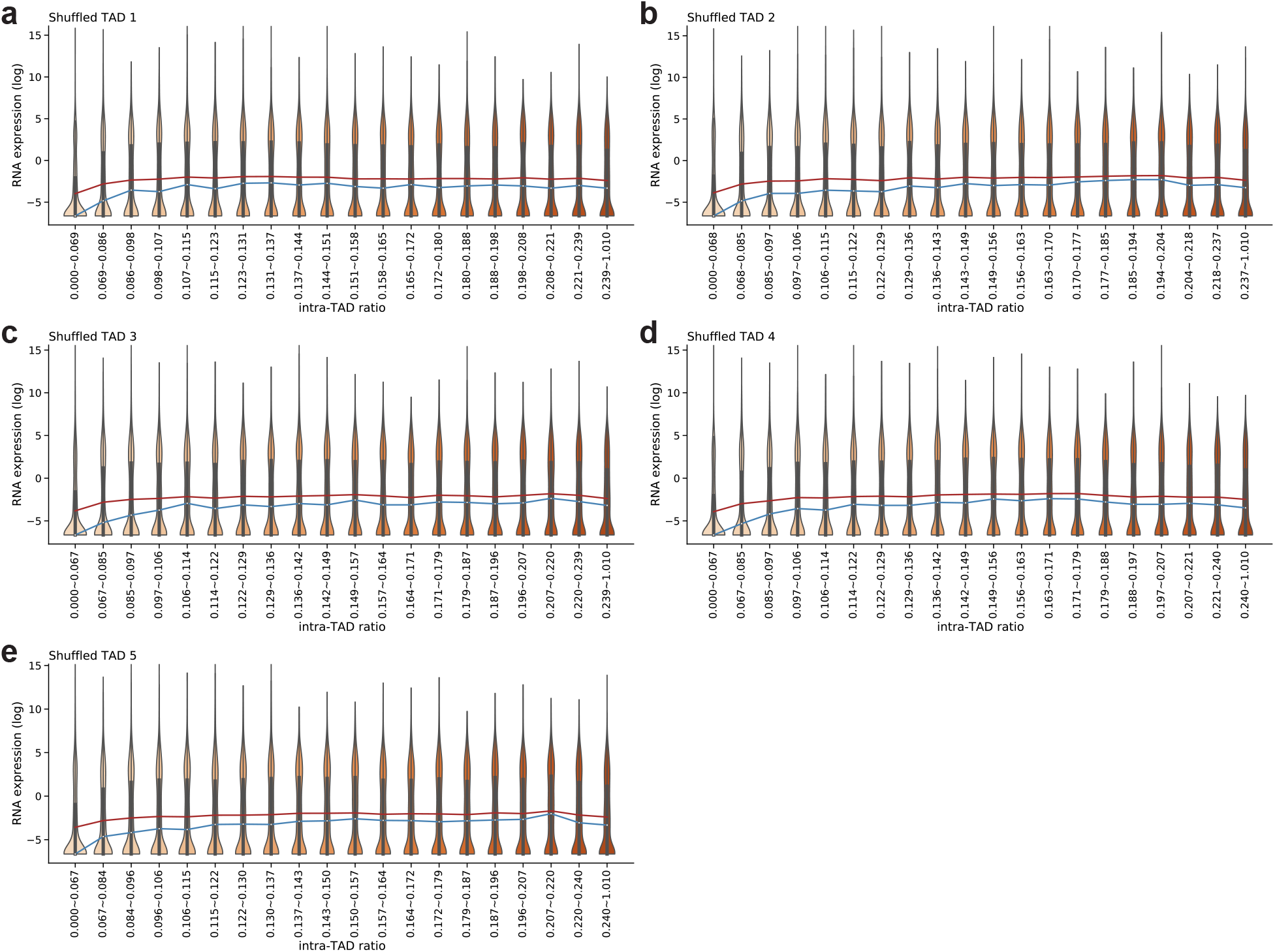
Relationship between intra-TAD ratio and gene expression for shuffled TADs in mESC. Violin plots of gene expression (log_2_(FPKM+0.01)) of TSS bins in groups with increasing intra-TAD ratio and similar number of bins, with intra-TAD ratio being calculated using five sets of shuffled TADs. The blue and red lines connect median and mean values of the different groups, respectively.

**Figure S19:**
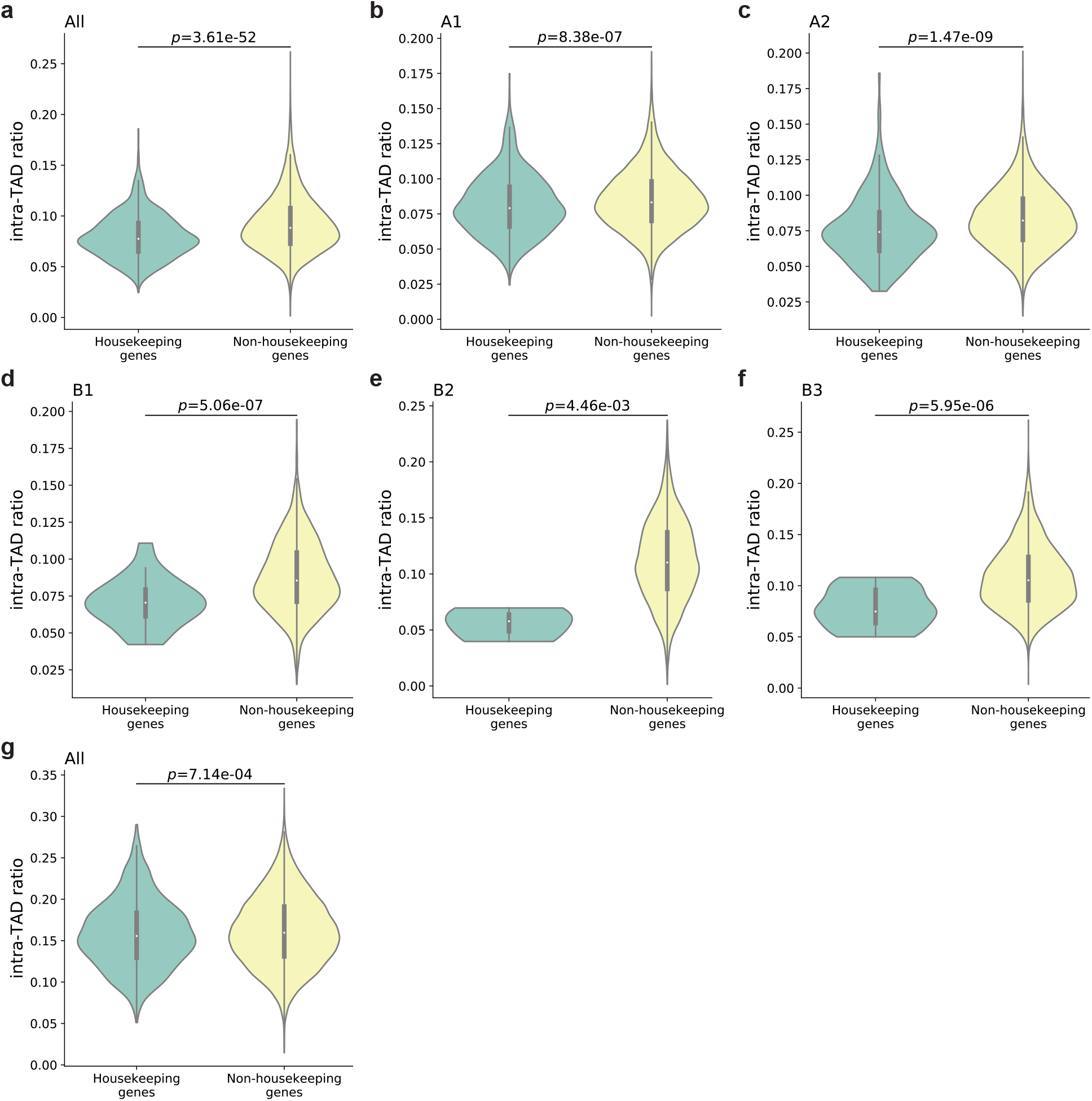
Violin plots of intra-TAD ratio of housekeeping genes and non-housekeeping genes when TAD boundaries and their neighboring regions were excluded in GM12878 (a-f) and mESC (g). *p*-values in all panels are calculated using the two-sided Mann-Whitney U test.

**Figure S20:**
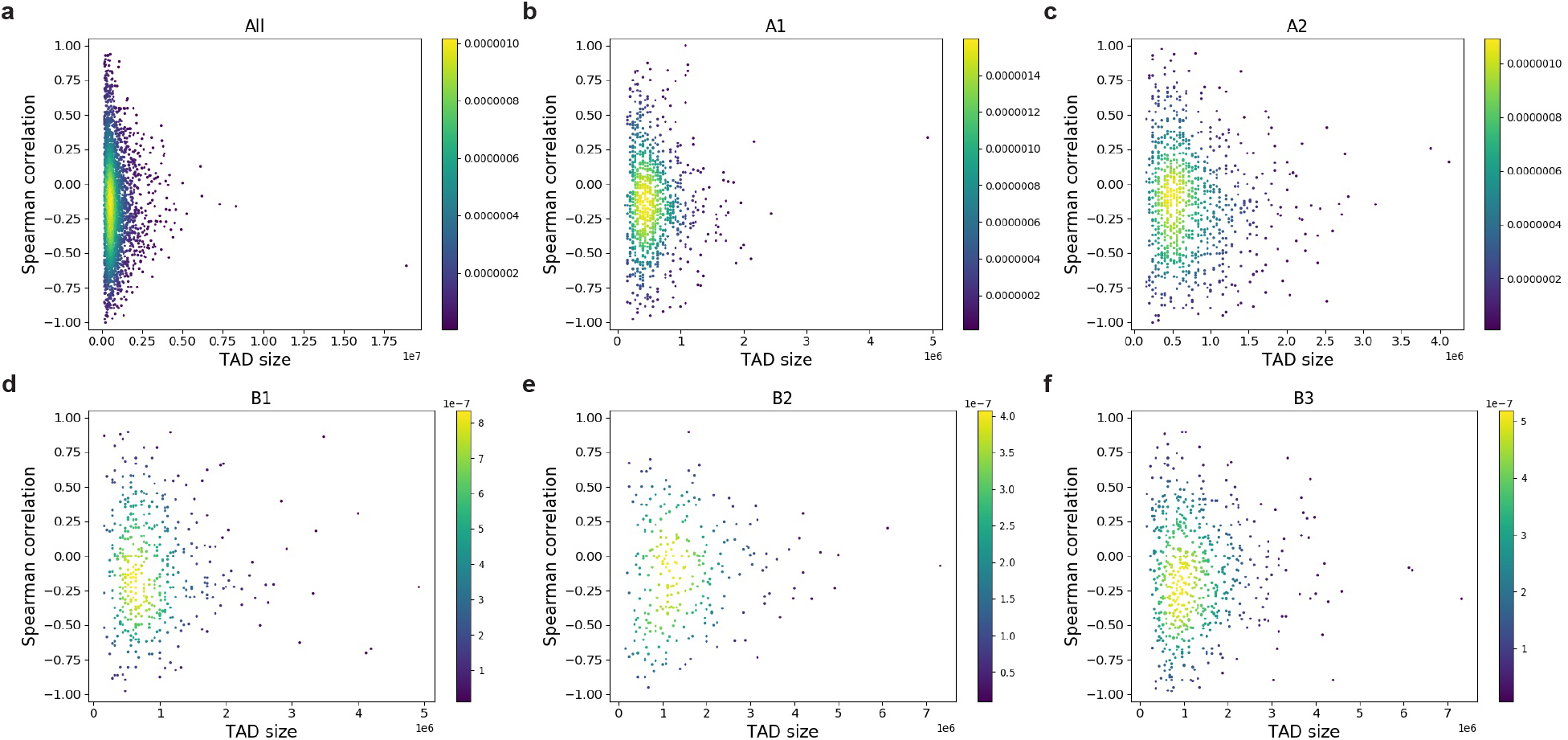
Density plots showing the Spearman correlation between intra-TAD ratio and gene expression of individual TADs versus TAD size for either all genomic genes (a) or genes in a genomic subcompartment (b-f).

**Figure S21:**
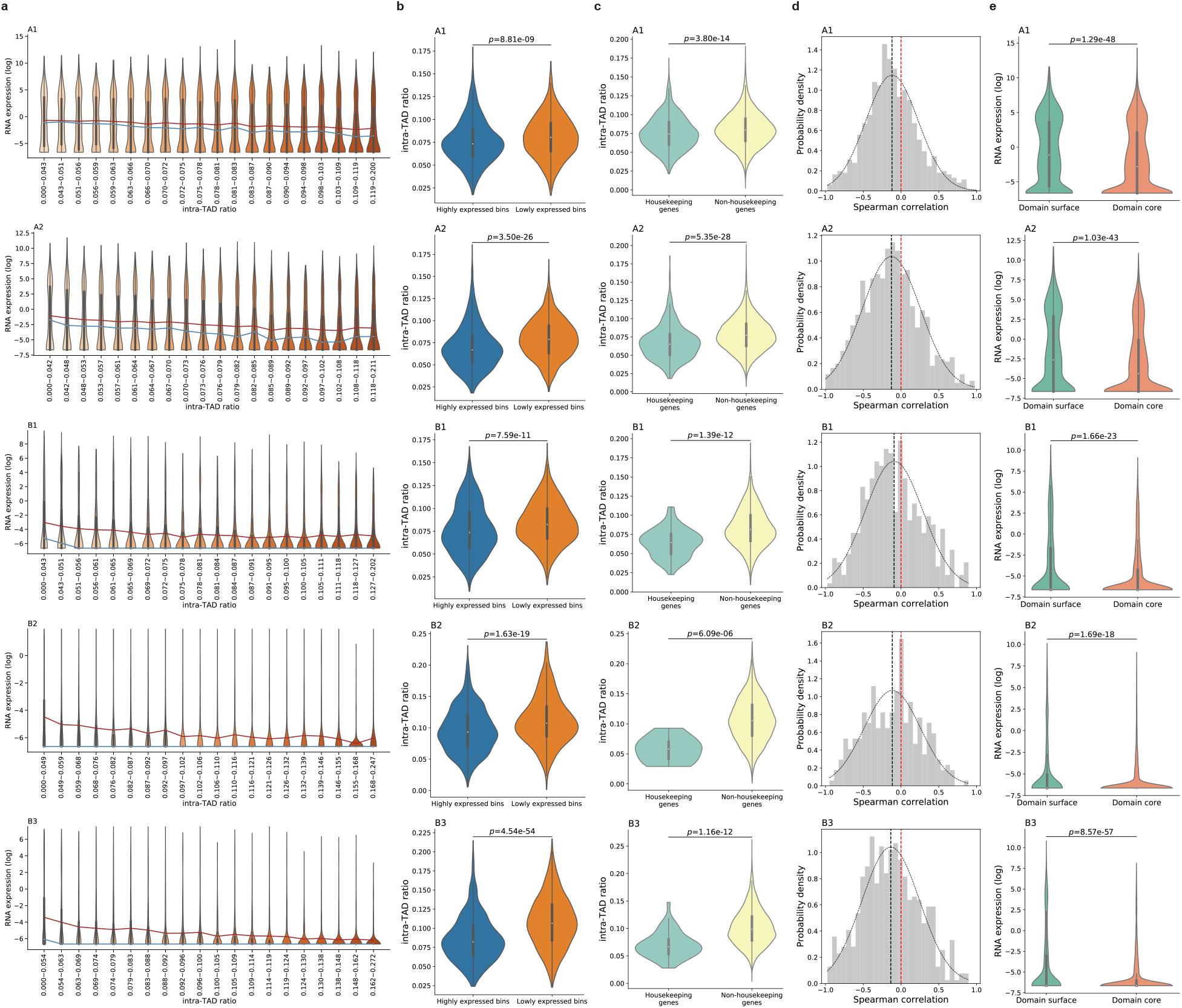
Intra-TAD ratio and gene expression have an inverse relationship in GM12878 for genes in individual subcompartments. (a) Violin plots of gene expression (log_2_(FPKM+0.01)) of TSS bins in groups with increasing intra-TAD ratio and similar number of bins. The blue and red lines connect median and mean values of the different groups, respectively. (b) Violin plots of intra-TAD ratios of the highly expressed bins and lowly expressed bins, defined as the genomic bins containing TSSs of genes whose FPKM values rank within top and bottom 1,000, respectively. (c) Violin plots of intra-TAD ratio of housekeeing genes and non-housekeeping genes. (d) Distribution of Spearman correlations between gene expression and intra-TAD ratio of genomic bins in individual TADs. Black dotted curve shows the fitted normal distribution. The vertical black dotted line shows the mean of Spearman correlations. The vertical red dotted line shows zero Spearman correlation. (e) Violin plots of gene expression levels (log_2_(FPKM+0.01)) in domain surface bins and domain core bins. *p*-values in all panels are calculated using the two-sided Mann-Whitney U test.

**Figure S22:**
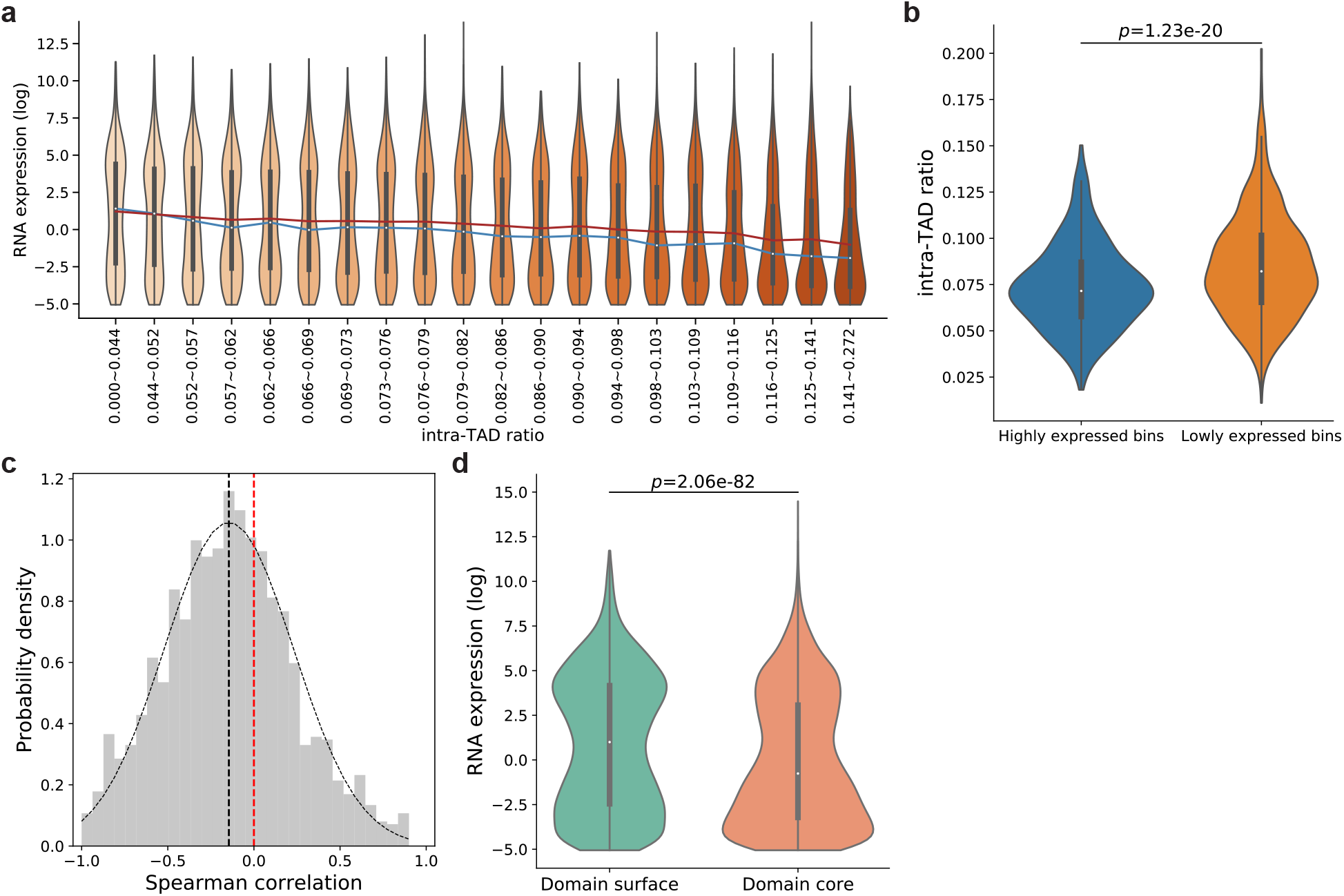
Intra-TAD ratio and gene expression have an inverse relationship in GM12878 even when genes with extremely low expression (RPKM*<*0.02) are ignored. (a) Violin plots of gene expression (log_2_(FPKM+0.01)) of TSS bins in groups with increasing intra-TAD ratio and similar number of bins. The blue and red lines connect median and mean values of the different groups, respectively. (b) Violin plots of intra-TAD ratios of the highly expressed bins and lowly expressed bins, defined as the genomic bins containing TSSs of genes whose FPKM values rank within top and bottom 1,000 after TSS bins with extremely low expression (RPKM*<*0.02) are excluded, respectively. (c) Distribution of Spearman correlations between gene expression and intra-TAD ratio of genomic bins in individual TADs. Black dotted curve shows the fitted normal distribution. The vertical black dotted line shows the mean of Spearman correlations. The vertical red dotted line shows zero Spearman correlation. (d) Violin plots of gene expression levels (log_2_(FPKM+0.01)) in Domain surface bins and Domain core bins. *p*-values in all panels are calculated using the two-sided Mann-Whitney U test.

**Figure S23:**
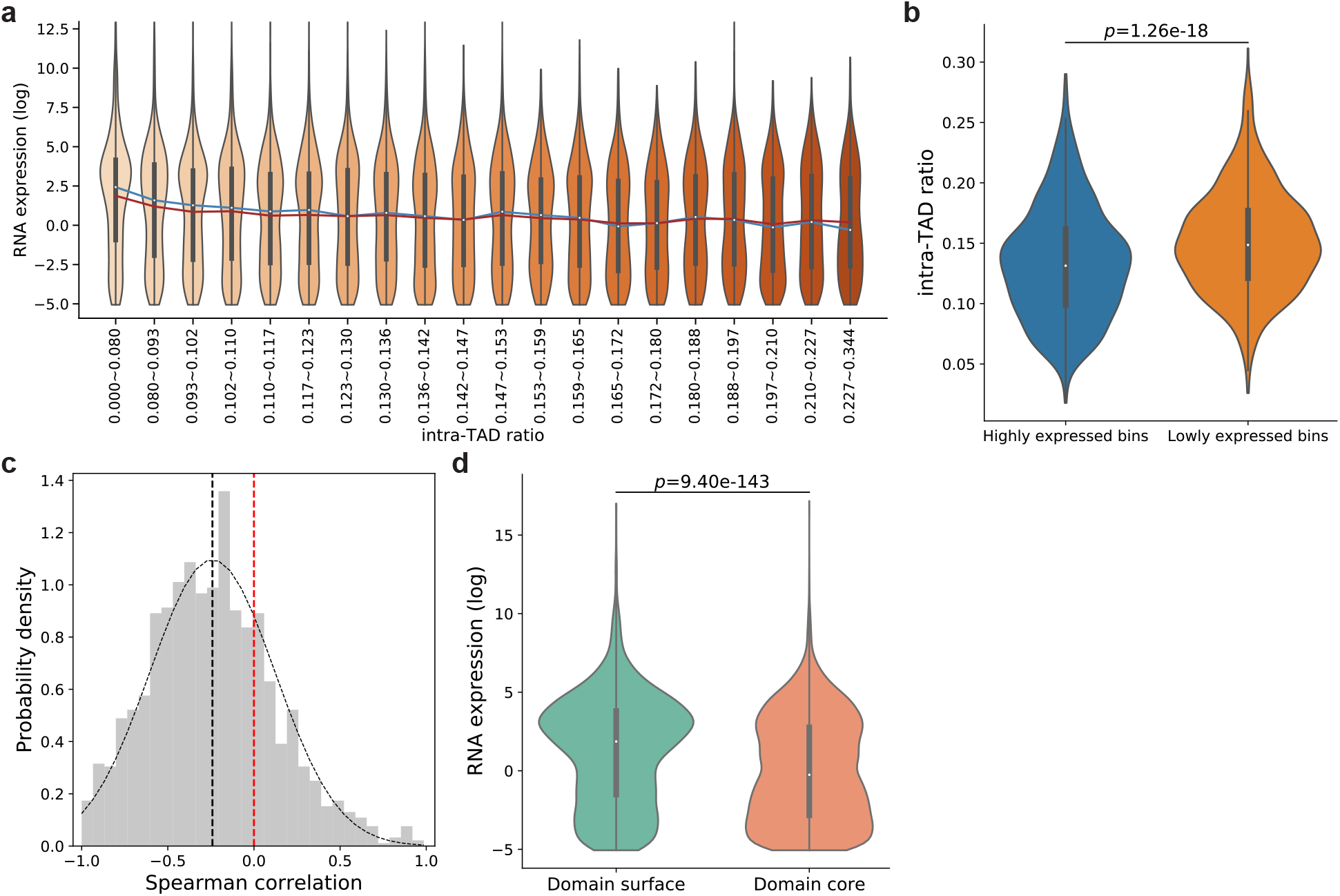
Intra-TAD ratio and gene expression have an inverse relationship in mESC even when genes with extremely low expression (RPKM*<*0.02) are ignored. (a) Violin plots of gene expression (log_2_(FPKM+0.01)) of TSS bins in groups with increasing intra-TAD ratio and similar number of bins. The blue and red lines connect median and mean values of the different groups, respectively. (b) Violin plots of intra-TAD ratios of the highly expressed bins and lowly expressed bins, defined as the genomic bins containing TSSs of genes whose FPKM values rank within top and bottom 1,000 after TSS bins with extremely low expression (RPKM*<*0.02) are excluded, respectively. (c) Distribution of Spearman correlations between gene expression and intra-TAD ratio of genomic bins in individual TADs. Black dotted curve shows the fitted normal distribution. The vertical black dotted line shows the mean of Spearman correlations. The vertical red dotted line shows zero Spearman correlation. (d) Violin plots of gene expression levels (log_2_(FPKM+0.01)) in Domain surface bins and Domain core bins. *p*-values in all panels are calculated using the two-sided Mann-Whitney U test.

**Figure S24:**
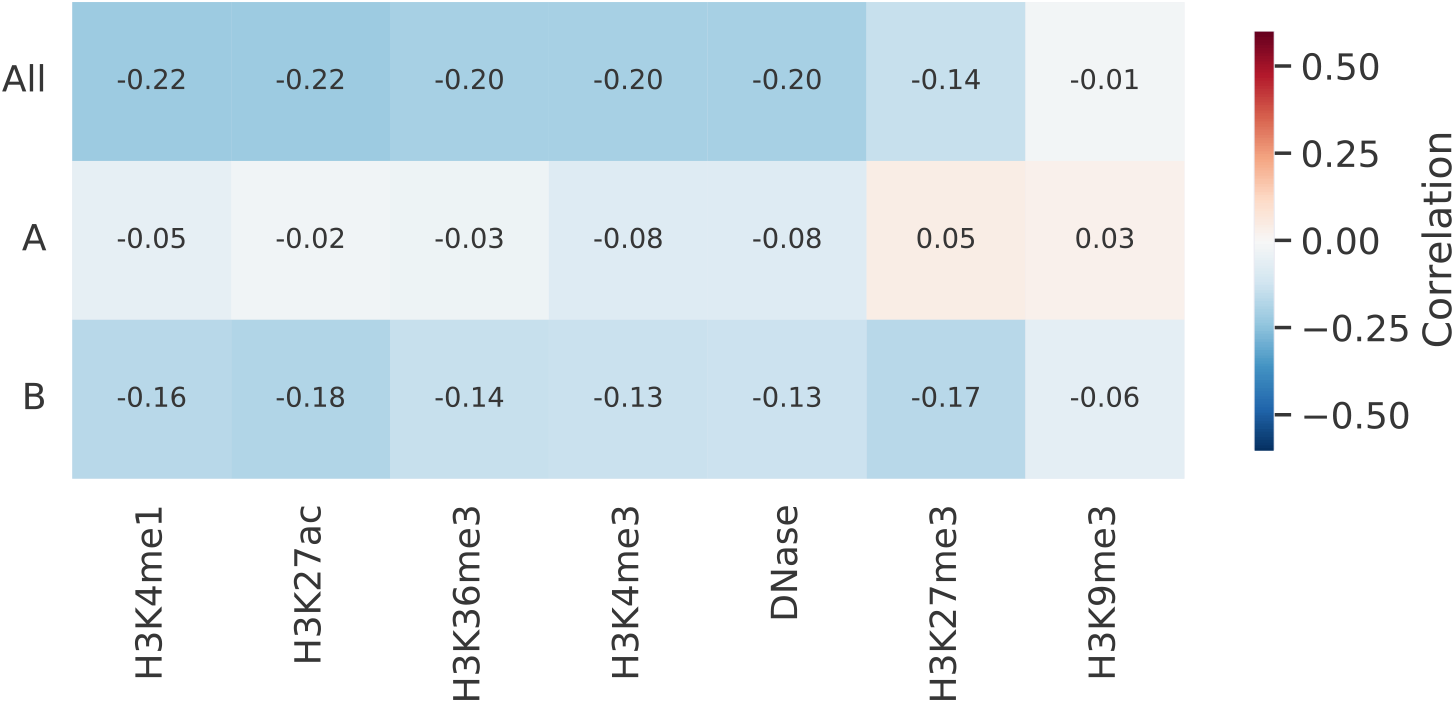
Spearman correlation between intra-TAD ratio and DNase-seq signals and 6 histone modifications across either all genomic bins or only bins in a genomic compartment in mESC. The columns are sorted by the correlation values when all TADs are considered (in the “All” row).

**Figure S25:**
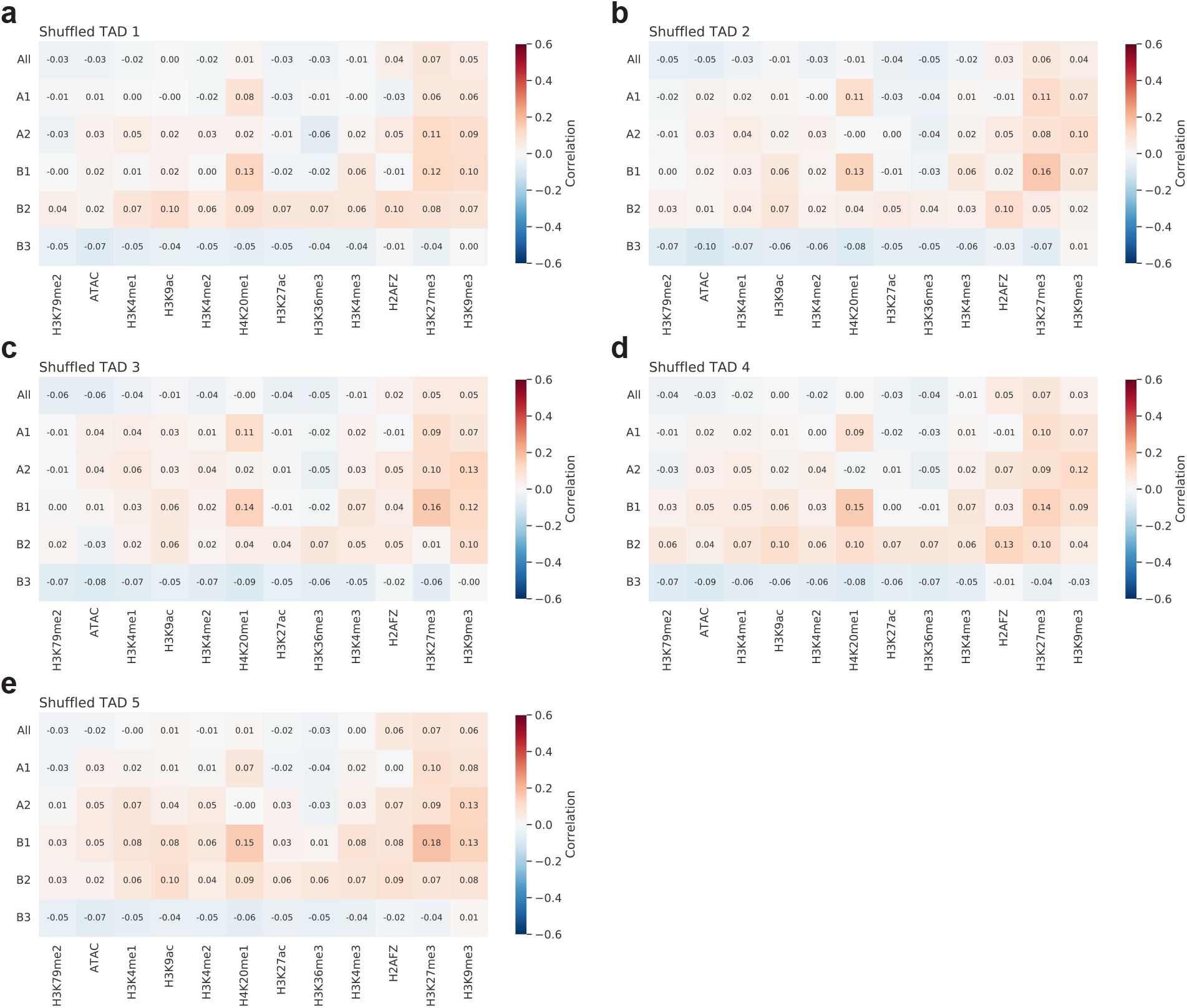
Spearman correlation between intra-TAD ratio and ATAC-seq signals and 11 histone modifications across either all genomic bins or only bins in a genomic subcompartment, with intra-TAD ratio being calculated using five sets of shuffled TADs. In each panel, the columns are sorted by the correlation values when all TADs are considered (in the “All” row).

**Figure S26:**
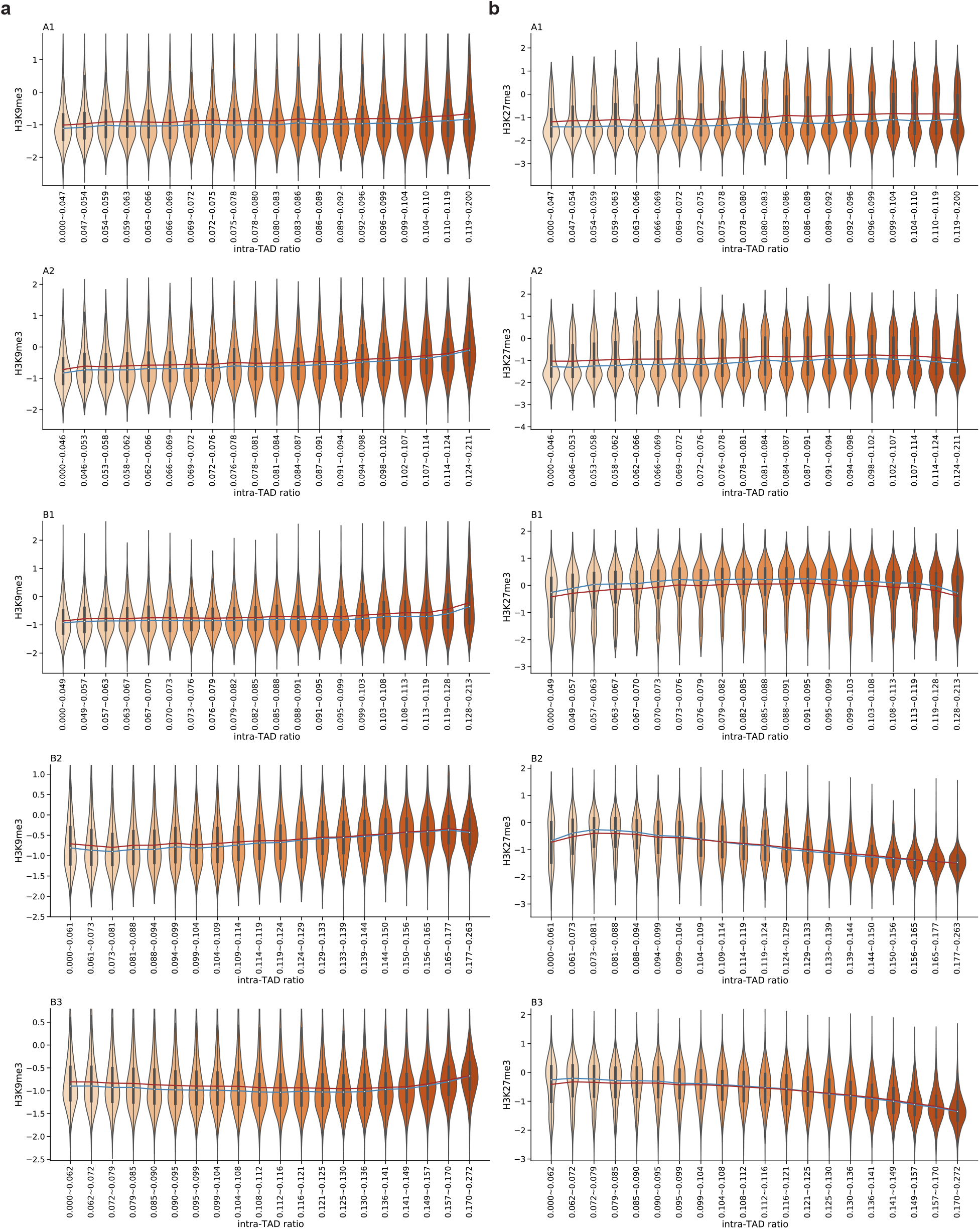
Violin plots of H3K9me3 (a) and H3K27me3 (b) signals in groups with increasing intra-TAD ratio and similar number of bins in individual subcompartments. y-axis shows log_2_(signal+0.01). The blue and red lines connect median and mean values of the different groups, respectively.

**Figure S27:**
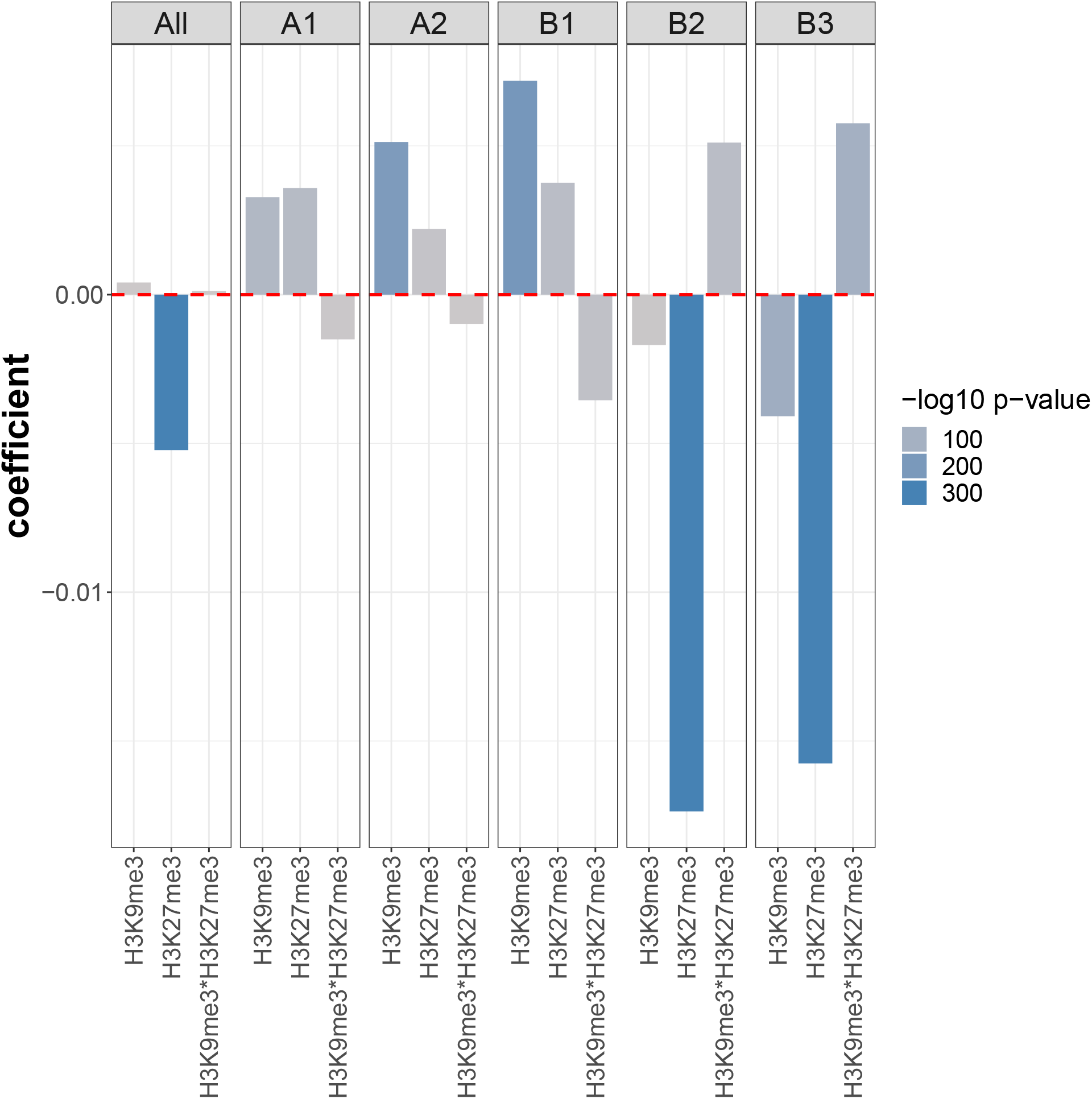
Coefficients of the linear regression models with H3K9me3, H3K27me3, and their interaction term as explanatory variables and intra-TAD ratio as the response variable, considering either all TADs or only TADs in a genomic subcompartment. The explanatory variables were all standardized individually before fitting the linear regression models.

**Figure S28:**
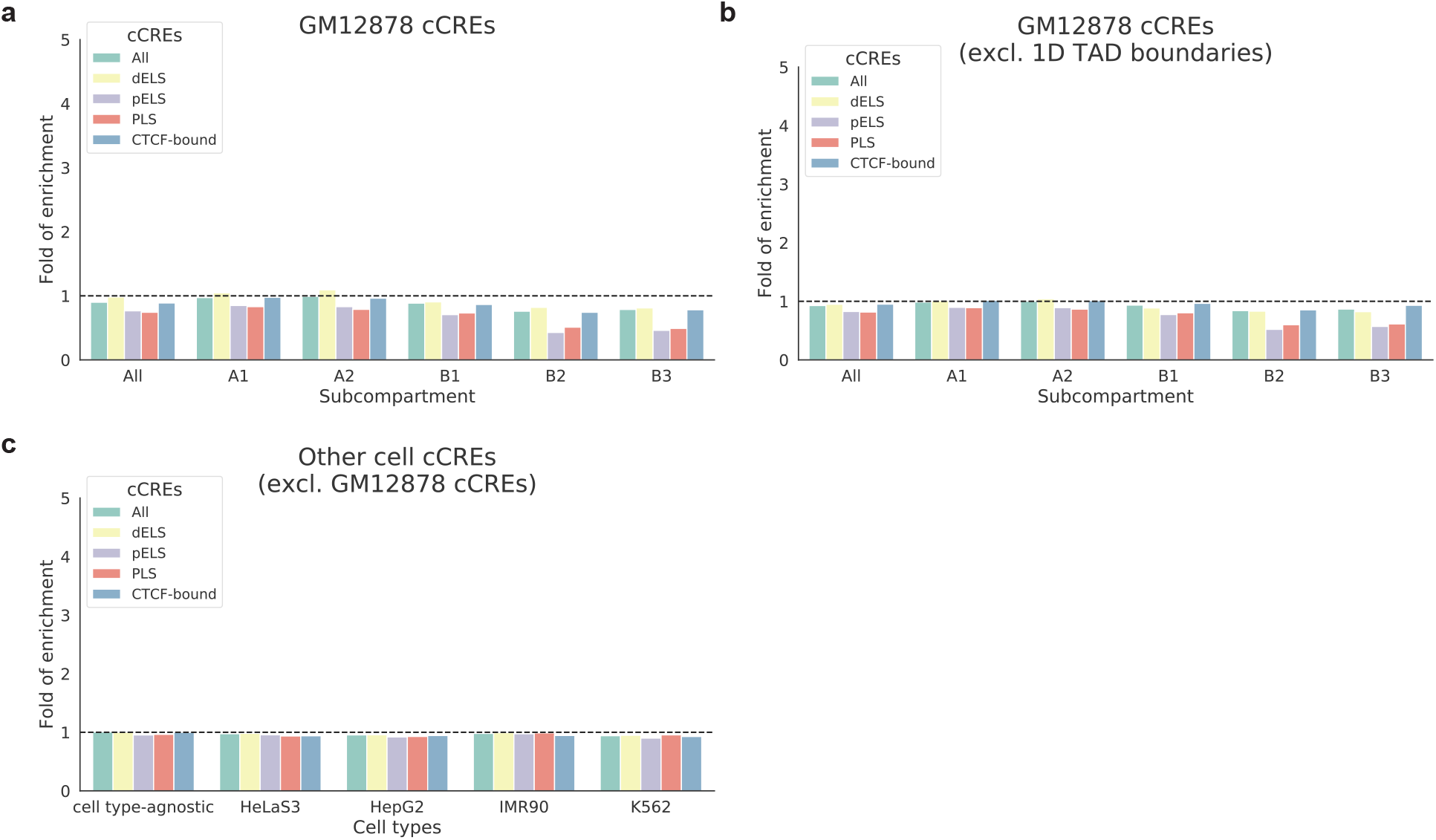
Depletion of CREs in chromatin domain cores. (a) Fold of enrichment of cCREs on domain cores in GM12878. (b) Fold of enrichment of cCREs on domain cores in GM12878 when the regions around TAD boundaries are excluded. (c) Fold of enrichment of cCREs on domain cores when the cCREs are cell type-agnostic or defined in other cell types and the GM12878-specific cCREs are excluded. cCREs All: all categories of cCREs; dELS: distal enhancer-like signatures; pELS: proximal enhancer-like signatures; PLS: promoter-like signatures; CTCF-bound: bound by CTCF.

**Figure S29:**
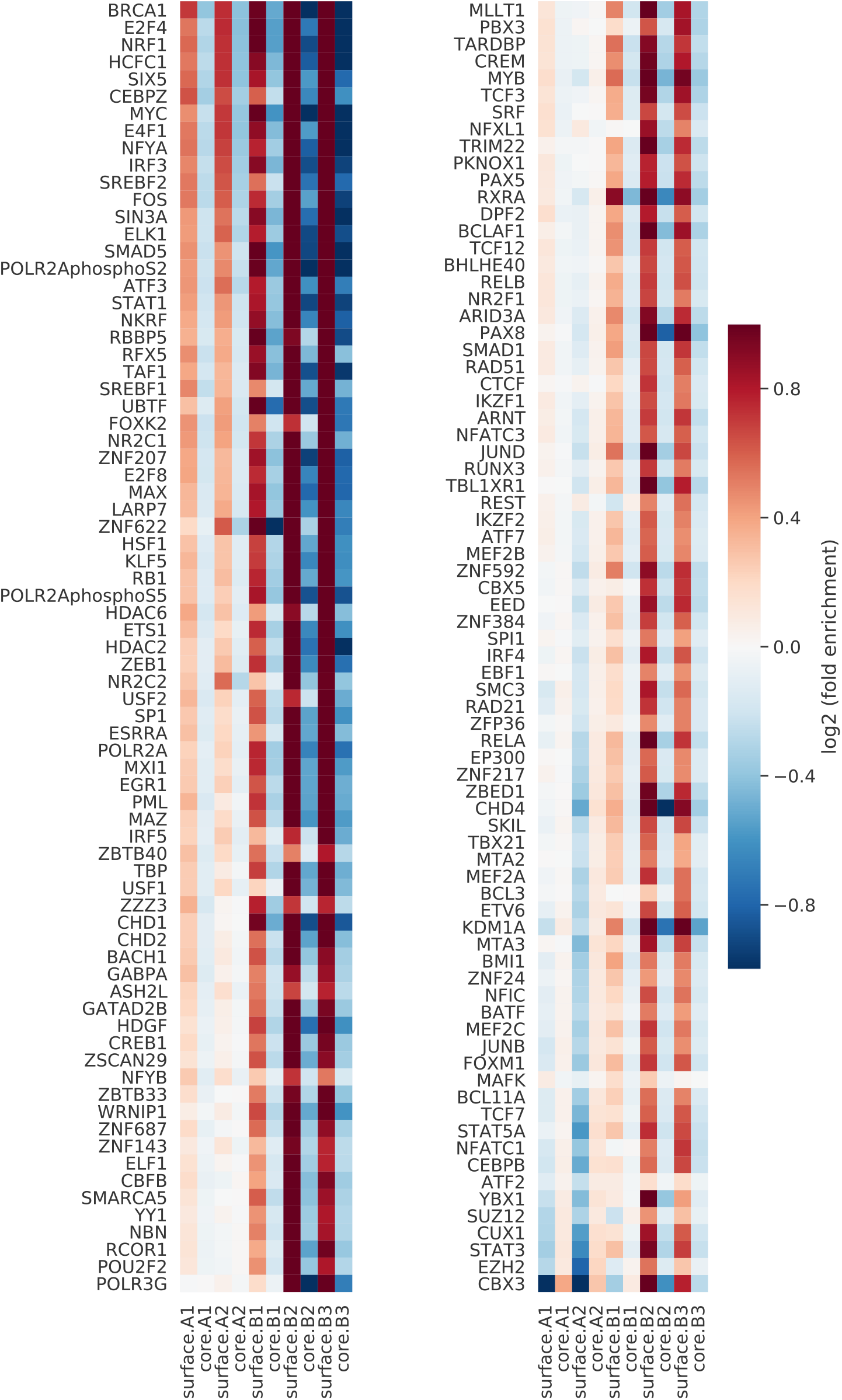
Heatmap showing log2(enrichment fold) of transcription factor binding sites in individual subcompartments on domain surfaces (odd-numbered columns) and domain cores (even-numbered columns) in GM12878.

**Figure S30:**
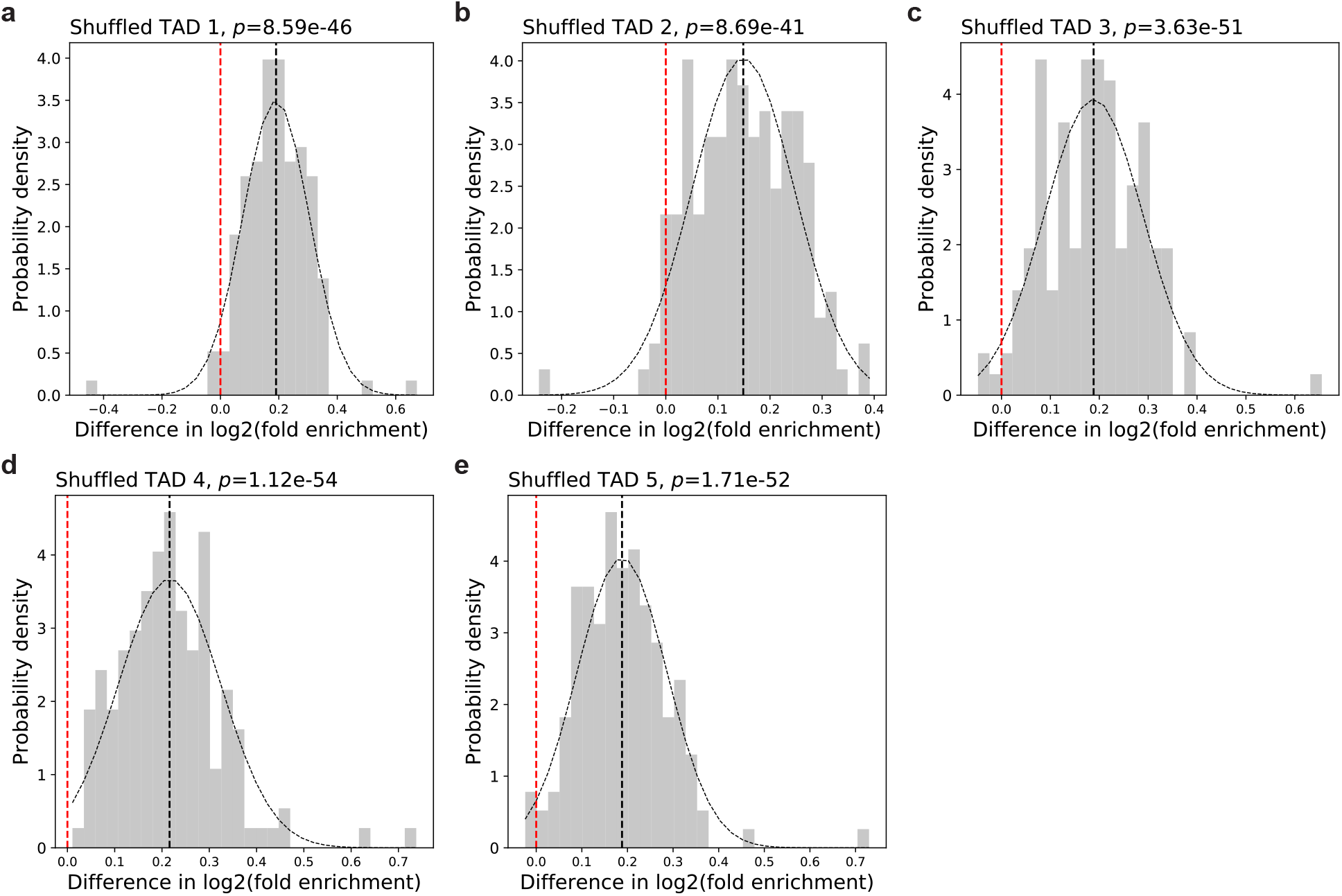
Distribution of difference in log enrichment fold of transcription binding sites on chromatin domain surfaces defined by intra-TAD ratio of real TADs and shuffled TADs. For each transcription factor, we calculated the log2 enrichment fold of transcription binding sites on domain surfaces defined by intra-TAD ratio of real TADs and shuffled TADs, respectively, and computed the difference (real TAD minus shuffled TAD). The distribution was plotted for all transcription factors. In all panels, black dotted curves show the fitted normal distribution. The vertical black dotted lines show the mean difference in log2(enrichment fold). The vertical red dotted lines show zero difference, as a reference. *p*-values are computed using the two-sided t-test.

**Figure S31:**
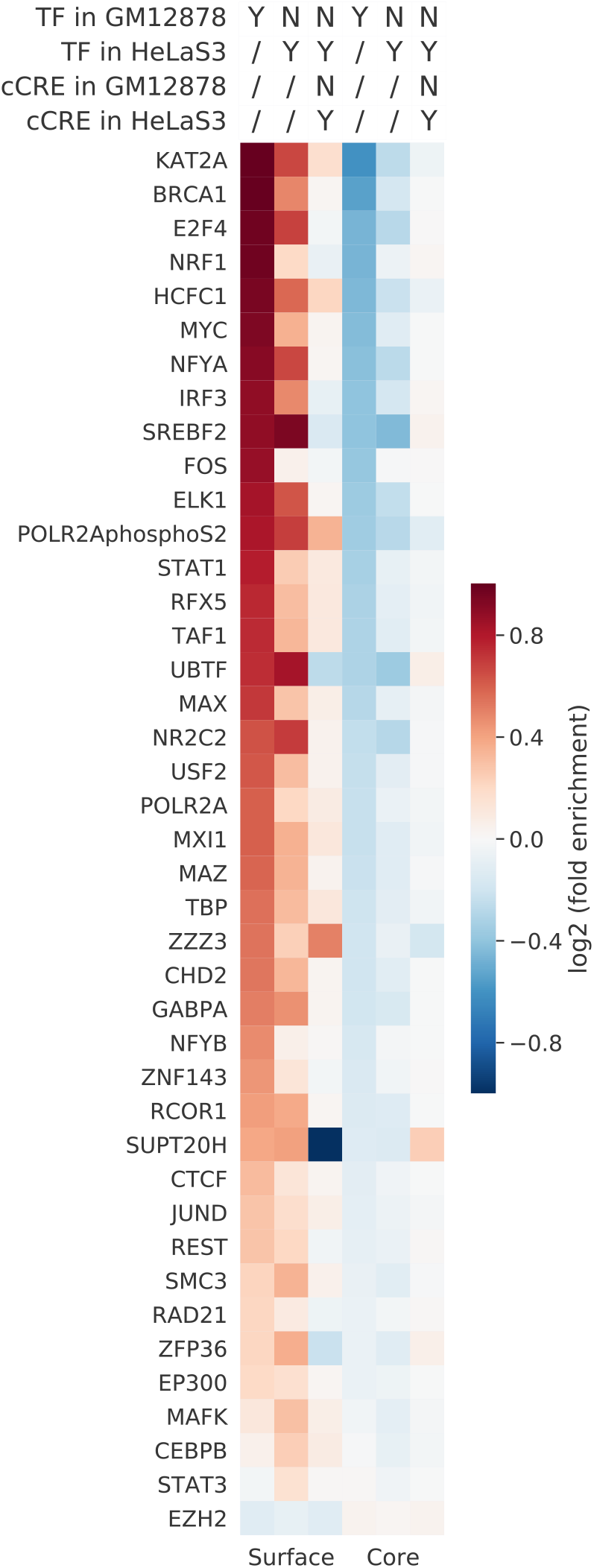
Heatmap comparing the locations of transcription factor binding sites and CREs in HeLaS3 and GM12878 relative to the TADs in GM12878. The first three columns show the enrichment of these locations on the GM12878 domain surfaces. The last three columns show the enrichment of these locations in the GM12878 domain cores. In the header, “Y” means only regions satisfying the condition are included, “N” means only regions not satisfying the condition are included, and “/” means the condition is not involved in defining the regions and thus no regions are excluded. The final set of regions is the intersection of the regions satisfying the different specified conditions.

**Figure S32:**
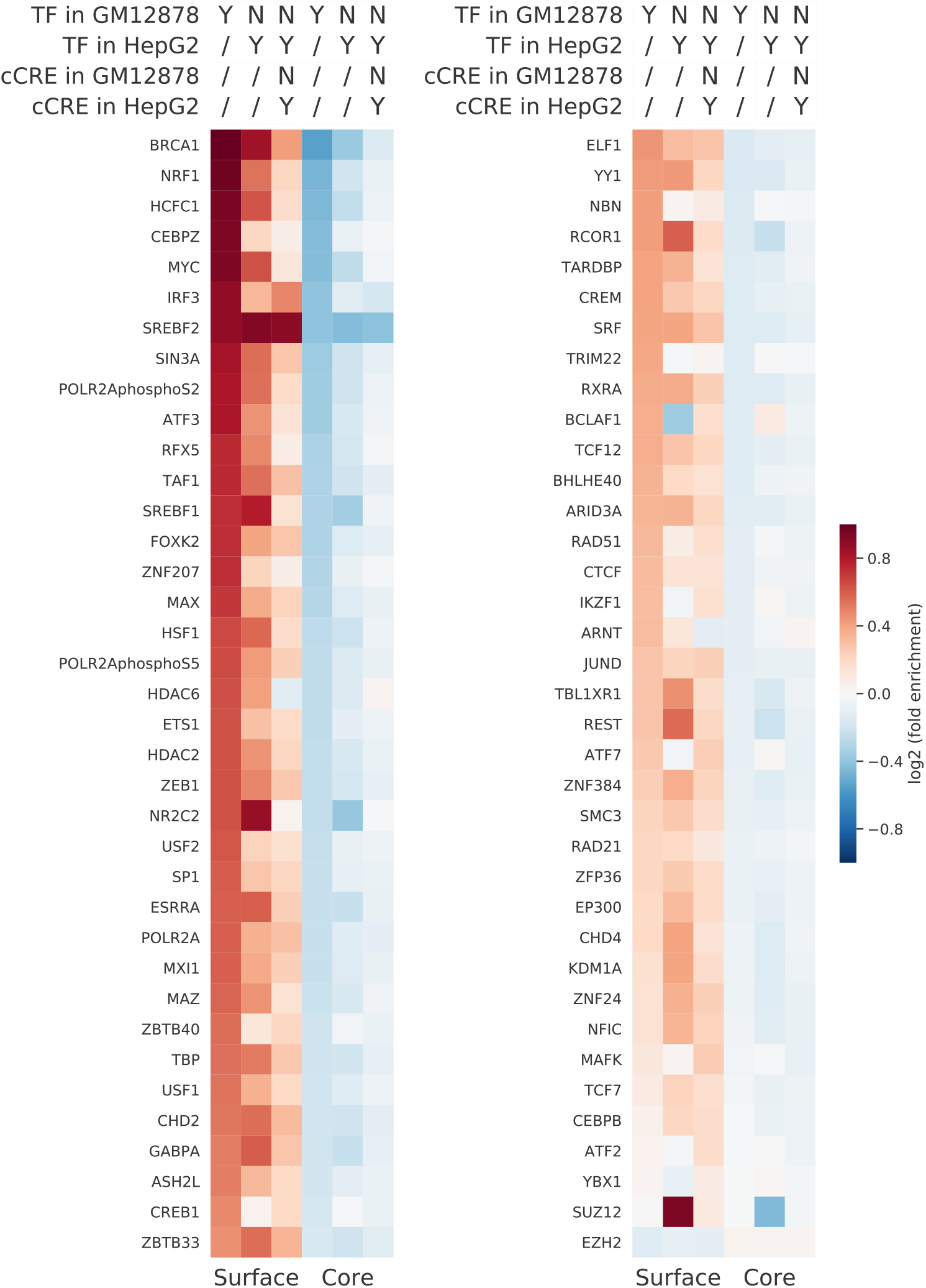
Heatmap comparing the locations of transcription factor binding sites and CREs in HepG2 and GM12878 relative to the TADs in GM12878. The first three columns show the enrichment of these locations on the GM12878 domain surfaces. The last three columns show the enrichment of these locations in the GM12878 domain cores. In the header, “Y” means only regions satisfying the condition are included, “N” means only regions not satisfying the condition are included, and “/” means the condition is not involved in defining the regions and thus no regions are excluded. The final set of regions is the intersection of the regions satisfying the different specified conditions.

**Figure S33:**
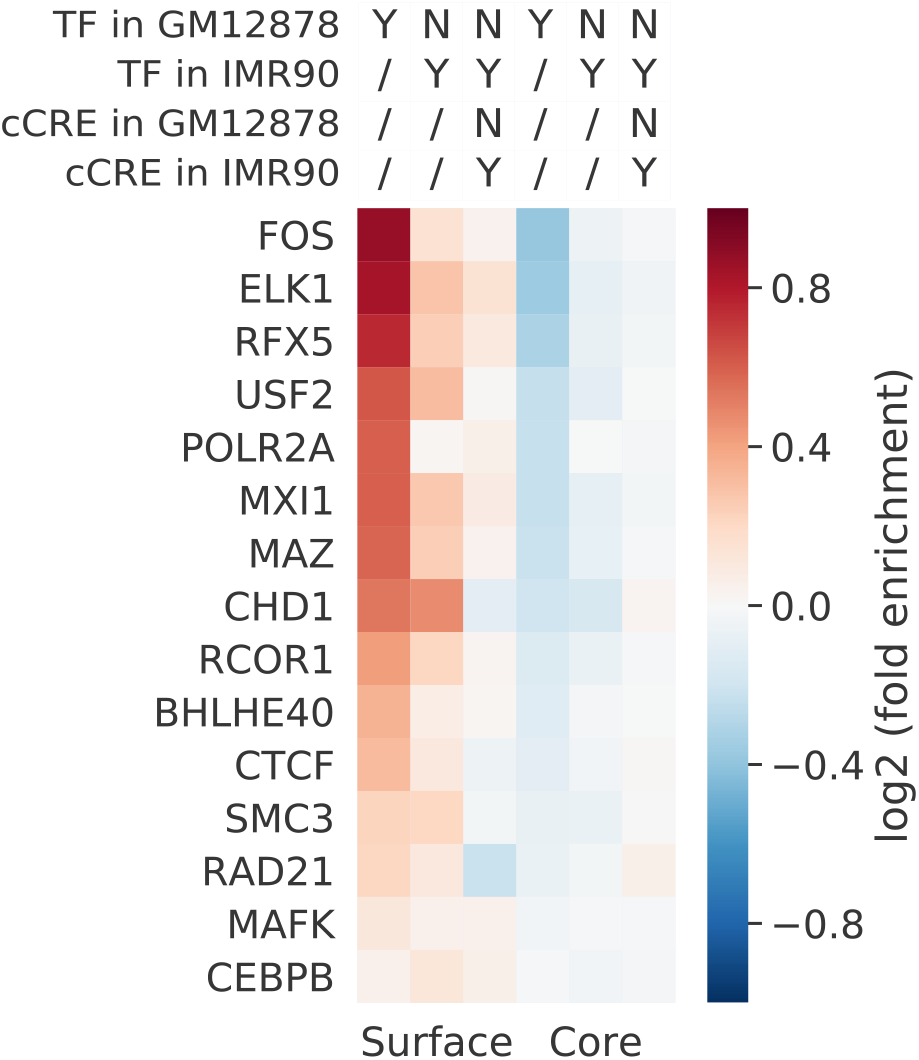
Heatmap comparing the locations of transcription factor binding sites and CREs in IMR90 and GM12878 relative to the TADs in GM12878. The first three columns show the enrichment of these locations on the GM12878 domain surfaces. The last three columns show the enrichment of these locations in the GM12878 domain cores. In the header, “Y” means only regions satisfying the condition are included, “N” means only regions not satisfying the condition are included, and “/” means the condition is not involved in defining the regions and thus no regions are excluded. The final set of regions is the intersection of the regions satisfying the different specified conditions.

**Figure S34:**
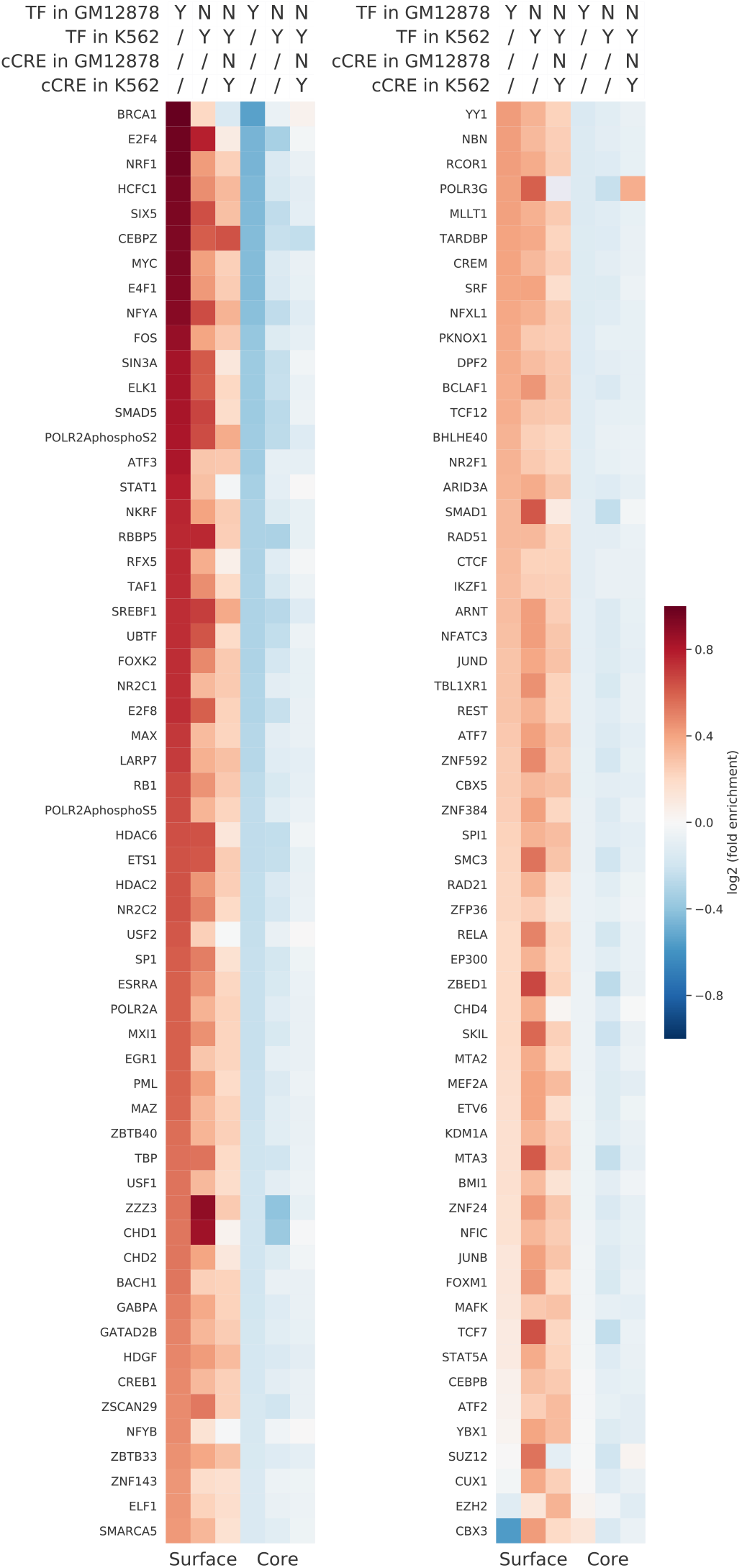
Heatmap comparing the locations of transcription factor binding sites and CREs in K562 and GM12878 relative to the TADs in GM12878. The first three columns show the enrichment of these locations on the GM12878 domain surfaces. The last three columns show the enrichment of these locations in the GM12878 domain cores. In the header, “Y” means only regions satisfying the condition are included, “N” means only regions not satisfying the condition are included, and “/” means the condition is not involved in defining the regions and thus no regions are excluded. The final set of regions is the intersection of the regions satisfying the different specified conditions.

**Figure S35:**
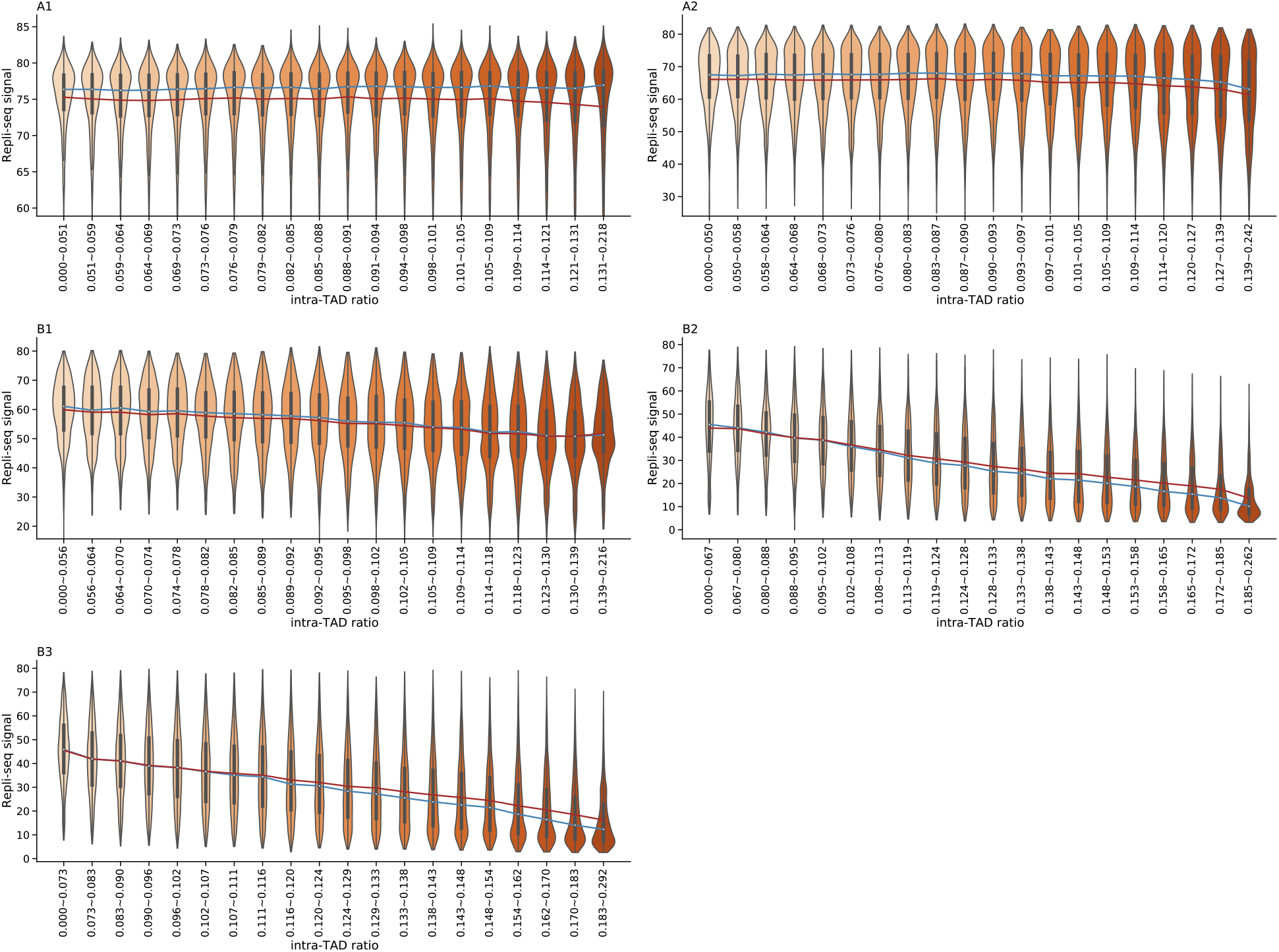
Violin plots of Repli-seq signals in groups with increasing intra-TAD ratio and similar number of bins in individual subcompartments. The blue and red line connects median and mean values, respectively, in each group.

